# ZC3HC1 is a structural element of the nuclear basket effecting interlinkage of TPR polypeptides

**DOI:** 10.1101/2021.12.30.474576

**Authors:** Philip Gunkel, Volker C. Cordes

**Author notes:** **Abbreviations:** aa, amino acid(s); AID, auxin-inducible degron; CLSM, confocal laser scanning microscopy; CRISPR/Cas9, Clustered Regularly Interspaced Short Palindromic Repeats/CRISPR-associated protein 9; CT, carboxy-terminus; FA, formaldehyde; FACS, fluorescence-activated cell sorting; IB, immunoblotting; IFM, immunofluorescence microscopy; KD, knockdown; KO, knockout; LNN, lamina-NPC-NB; MCL, master cell line; NB, nuclear basket; NB-s, NB-stabilising; NE, nuclear envelope; NPC, nuclear pore complex; NR, nuclear ring; NT, amino-terminus; NUP, nucleoporin; sdAbs, single-domain antibodies; sfGFP, superfolder green fluorescent protein; TR, terminal ring.

## Abstract

The nuclear basket (NB), anchored to the nuclear pore complex (NPC), is commonly thought of as built solely of protein TPR polypeptides, the latter thus regarded as the NB’s only scaffold-forming components. In the current study, we report ZC3HC1 as a second building element of the NB. Recently described as an NB-appended protein omnipresent in vertebrates, we now show that ZC3HC1, both *in vivo* and *in vitro*, enables in a step-wise fashion the recruitment of TPR subpopulations to the NB and their linkage to already NPC-anchored TPR polypeptides. We further demonstrate that the degron-mediated rapid elimination of ZC3HC1 results in the prompt detachment of the ZC3HC1-appended TPR polypeptides from the NB and their release back into the nucleoplasm again, underscoring the role of ZC3HC1 as a natural structural element of the NB. Finally, we show that ZC3HC1 can keep TPR polypeptides positioned even at sites remote from the NB, in line with ZC3HC1 functioning as a protein connecting TPR polypeptides.

## Introduction

The nuclear basket (NB) is a delicate fibrillar structure of eightfold-rotational symmetry appended to the nuclear side of the nuclear pore complex (NPC). Common to many eukaryotes and regarded as an NPC-appended entity probably present in almost all cell types in vertebrates, the prototypic NB is composed of rectilinear fibrils that emanate from the nuclear ring (NR) of the NPC, bifurcate at their distal ends, and laterally interconnect with their neighbouring fibrils, thereby forming a structure referred to as the NB’s terminal ring (TR). A range of diverse functions have been ascribed to the NB and to some of its attributed components in different cell types and species, including roles in perinuclear chromatin organisation, gene expression regulation and nucleocytoplasmic transport, but a universal, cell type-spanning function of this structure remains to be unveiled (Krull *et al*, 2010; Strambio-De-Castillia *et al*, 2010; Niepel *et al*, 2013; Snow & Paschal, 2014; Aksenova *et al*, 2020; Ashkenazy-Titelman *et al*, 2020; Bensidoun *et al*, 2021).

The NB scaffold in vertebrates is regarded as being built of numerous copies of a protein called TPR (e.g., Cordes *et al*, 1997; Frosst *et al*, 2002; Krull *et al*, 2004). This large protein features a long, coiled-coil-dominated amino-terminal domain that allows for forming the NB fibrils and an additional carboxy-terminal domain that, for the most part, is intrinsically disordered and thus considered highly flexible (e.g., Mitchell & Cooper, 1992; Hase *et al*, 2001). Vertebrate TPR and its homologs in other phyla are commonly considered the only scaffold-forming elements of these organisms’ NBs, to which several additional NB-resident proteins are merely appended. Among these are, in vertebrates, the Sumo protease SENP1 (e.g., Schweizer *et al*, 2013; Duheron *et al*, 2017), the cell cycle checkpoint regulators MAD1/Mad1p and MAD2/Mad2p (e.g., Lee *et al*, 2008; Schweizer *et al*, 2013), the components of the mRNA export complex TREX-2 (e.g., Umlauf *et al*, 2013; Wickramasinghe *et al*, 2014; Aksenova *et al*, 2020), and the ubiquitin E3 ligase COP1/RFWD2 (Yi *et al*, 2006; Ouyang *et al*, 2020). With these NB-appended proteins looked upon as neither required for NB assembly nor structural elements, they have been considered using the scaffold provided by TPR as either an operational platform for conducting their own tasks or as a storage place from which they are recruited to other subcellular locations upon demand.

Recently, protein ZC3HC1 has been presented as another TPR-interacting component of the NB, located at the TR of the NB structure, and, even though not an essential protein, it had been found occurring within probably most if not all cell types of different morphogenetic origins in which TPR is present as well (Gunkel *et al*, 2021). While formerly described as a nuclear protein with other functions and as engaging in interactions with other proteins (e.g., Ouyang *et al*, 2003; Bassermann *et al*, 2005, 2007; Klitzing *et al*, 2011; Illert *et al*, 2012; Gengenbacher *et al*, 2019) that are neither part of the NPC nor NB, ZC3HC1 had then been shown to not interact naturally with such proteins and instead be located nearly exclusively at the NB in all of several different cell types investigated (Gunkel *et al*, 2021).

Moreover, different from the other NB-resident proteins, ZC3HC1 deficiency had turned out causing a substantial proportion of TPR’s normally NB-associated amounts to then no longer be located there, with the latter then instead either remaining in a soluble state or no longer detectable at all (Gunkel *et al*, 2021). Inspection of different cell lines had further revealed that their NPC-associated amounts of TPR actually consisted of at least two, then similarly large subpopulations, with the one located at the NPC independent of ZC3HC1 and the other requiring the presence of this protein. However, it had remained unknown how ZC3HC1 would ultimately contribute to such positioning of TPR subpopulations at the NB, as the data available had not allowed for distinguishing unequivocally between different scenarios imaginable in which ZC3HC1 would either play some indirect or direct role. Moreover, regarding the latter case, ideas ranged from a role in the initial recruitment of TPR, or some other steps in the NB assembly process, to some unknown role in maintaining NB integrity.

Having scrutinised these conceived scenarios in the current study, we can now present ZC3HC1 as a second structural element of the NB. As such, ZC3HC1 functions as a component that allows for stable interconnections between different populations of TPR polypeptides. We reflect on how the additional ZC3HC1-dependent TPR polypeptides might allow for modulating, possibly also dynamically, some of the NB’s functional properties.

## RESULTS

Recently, we had found that the destruction of ZC3HC1 transcripts by RNAi or the disruption of all *ZC3HC1* alleles by CRISPR/Cas9n technology resulted in a substantial amount of the NB scaffold protein TPR being no longer positioned at the NB, and then instead often seen distributed throughout the nucleoplasm (Gunkel *et al*, 2021). However, it had remained unknown whether these findings correlated with a direct or indirect role of ZC3HC1, either in the initial recruitment or adhering of specific TPR subpopulations to the NB or in the subsequent stabilisation of their interactions at this site. In other words, with the data available until then, we only knew that the presence of ZC3HC1 was necessary for subpopulations of TPR to occur appended to the NB, in addition to TPR amounts already present at the NPC, but not why this was so.

In the current study, we addressed these questions and aimed to unravel which role ZC3HC1 might play in the recruitment and appendage of additional TPR polypeptides and in keeping them positioned at the NB. For the sake of convenience, we will, in the following, sometimes refer to those TPR polypeptides that occur anchored at the NPC in a ZC3HC1-independent manner as the TPR pool 1 (TPR 1 or T1) polypeptides. Accordingly, those located at the NE only in the presence of ZC3HC1, while otherwise soluble within the nuclear interior, will hence be referred to as the TPR polypeptides of pool 2 (TPR 2 or T2).

### ZC3HC1 is directly involved in the *in vivo* attraction of soluble TPR polypeptides that can be additionally appended to TPR already anchored at the NPC

One of the current study’s first goals was to gain insight into the course of events leading to the joint residency of ZC3HC1 and the T2 pool of TPR at the NB and to answer whether ZC3HC1 can promote the recruitment of such TPR 2 polypeptides to this site.

To this end, we first investigated whether the conspicuous nuclear amounts of soluble TPR that exist in some of the ZC3HC1 KO cell lines could be brought to being attached to the NE again if one would manage to provide sufficiently large amounts of newly synthesised ZC3HC1 rapidly. Ideally, such deployment of ZC3HC1 would occur by induced ectopic expression within a time span so short that no large amounts of TPR could be newly synthesised in the meantime and that it would instead primarily be the already existing soluble polypeptides of pool T2 that would be available as potential binding partners for the newly appearing ZC3HC1 polypeptides (for further considerations along this line, see also Supplemental Information 1).

For conducting such experiments, we used ZC3HC1 KO cell lines in which we had tagged all *TPR* alleles with superfolder GFP (sfGFP; Pédelacq *et al*, 2006) and in which a conspicuous nuclear pool of soluble sfGFP-tagged TPR polypeptides was then present (Supplemental Fig S1 and S2; our unpublished data). To create such cell lines, we had used the formerly isolated and characterised ZC3HC1 KO cells of lines HCT116 and HeLa (Gunkel *et al*, 2021), and eventually, we isolated cell lines of both HCT116 and HeLa that were homozygously expressing sfGFP-tagged TPR polypeptides. However, while yielding very similar results for all of those experiments that we had then performed with both cell lines for comparison, we used only the HeLa-originating cell lines for each one of the experiments that will be presented in the context of Figure 1 in the following, only sometimes complemented by the same type of experiment conducted with the HCT116 cell lines.

**Figure 1.**
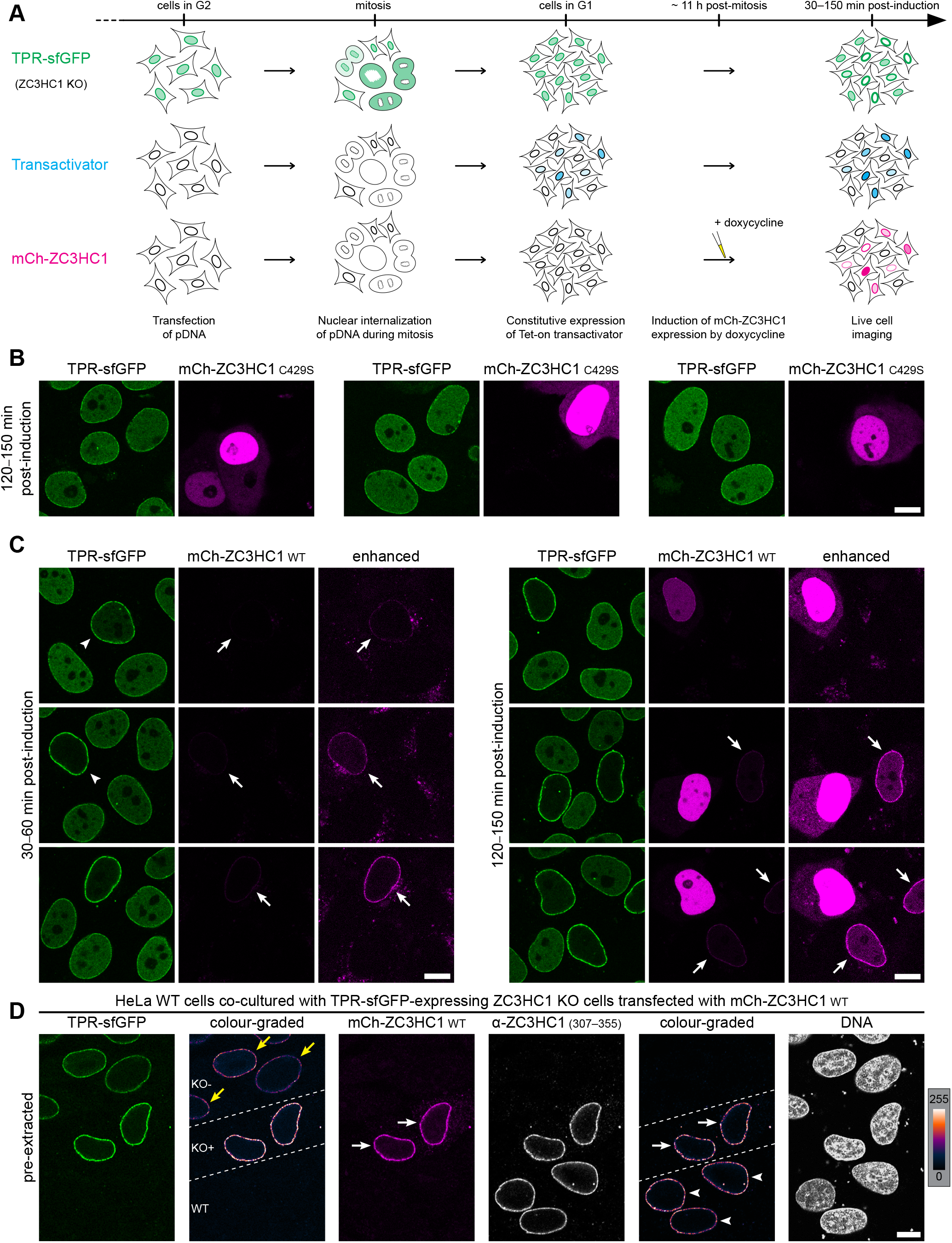
Ectopically expressed ZC3HC1 allows for depleting the nuclear pool of soluble TPR in ZC3HC1 KO cells by enabling its recruitment to the NE. **(A)** Timeline displaying the chronological order of procedures in this experiment schematically. Note that after transfection in G2, the all-in-one vectors encoding for the constitutively expressed transactivator and either a doxycycline-inducible WT or mutant version of mCherry-tagged *Hs*ZC3HC1 (see also Supplemental Fig S3) were considered incorporated into nuclei mostly during nuclear reassembly later in mitosis. This resulted in constitutive expression and gradual accumulation of the Tet-On transactivator directly from the G1 phase on, which allowed for inducing the actual reporter gene expression (here the WT version depicted) by adding doxycycline at about 11 h post-mitosis. **(B)** Live-cell fluorescence microscopy of TPR-sfGFP-expressing HeLa ZC3HC1 KO cells that had been transfected with an all-in-one vector encoding for the mutant version C429S of ZC3HC1 and that were inspected 120 to 150 min post-induction with doxycycline. Exemplifying micrographs show a selection of transfected cells with differing amounts of the ectopically expressed mutant protein. Note that even upon conspicuous expression of this TPR-binding-incompetent mutant, the nucleoplasmic pool of TPR-sfGFP remained unaffected, being indistinguishable in appearance from that of neighbouring cells that had not been transfected. Bar, 10 µm. **(C)** ZC3HC1 KO cells like those in B but transfected with an expression vector encoding for the mCherry-tagged intact WT version of ZC3HC1 capable of binding to the NE. These cells were inspected either 30–60 min or 120–150 min post-induction. Microscope settings were identical at the different time points, as was the degree of mCherry signal enhancement that was only done after image acquisition. Exemplifying micrographs show a selection of transfected cells with differing amounts of the ectopically expressed protein. Note that these images cannot claim to represent a proper time course experiment that allows for correlating time span with gradual protein accumulation within transfected cells at different time points, simply because plasmid copy numbers per cell likely vary significantly between cells. Nonetheless, though, note that already after short lengths of time, the nuclear pool of GFP-tagged TPR was notably diminished or no longer detectable at all (two exemplifying nuclei marked by arrowheads). The latter was even so in those transfected cells (several demarked by arrows) in which the amounts of ZC3HC1 synthesised till then were barely detectable via their mCherry tags at standard microscope settings, likely due to the relatively slow and here not yet completed mCherry maturation, which in turn also prompted the presentation of electronically brightness-enhanced images. Bar, 10 µm. **(D)** IFM of cell-cycle-synchronised cells that had been transfected with the all-in-one mCherry-ZC3HC1 expression vector and treated with doxycycline as outlined in 1A, yet having started with a mixed population of TPR-sfGFP-expressing ZC3HC1 KO cells and HeLa WT cells expressing ZC3HC1 and non-tagged TPR. Such mixed populations allowed for comparing the amount of endogenous ZC3HC1 at the WT cell’s NE relative to that eventually occurring at the ZC3HC1 KO cell’s NE, here focusing on those KO cells in which the mCherry signal intensity at the NE seemed to have reached a maximum. Cells presented here had been harvested 120 min post-induction. They had then been detergent-permeabilised before fixation, resulting in the removal of (i) the soluble pool of TPR still existing in those KO cells not transfected and of (ii) the surplus of soluble mCherry-ZC3HC1 for which no further binding sites existed at the NEs of the transfected cells, which in turn allowed for better assessment of signal intensities at only the NEs. The cells were then fixed with FA before being labelled with a *Hs*ZC3HC1 antibody and a sdAb specific for mCherry. Micrographs showing sfGFP-tagged TPR and antibody-labelled ZC3HC1 are also shown colour-graded to display differences in pixel intensities via a colour look-up table (LUT), with areas harbouring WT cells, two transfected KO cells (KO+), and some non-transfected KO cells (KO-) marked accordingly. Note that the intensity of ZC3HC1 immunolabelling at the WT cells’ NEs (marked by arrowheads in the colour-graded micrograph on the right side) and at the NEs of those ZC3HC1 KO cells that had been ectopically expressing mCherry-ZC3HC1 (white arrows) were very similar. As an aside, also note once again that TPR signal intensities at the NEs of those KO cells ectopically expressing ZC3HC1 (see KO+ cells in the colour-graded micrograph on the left) were notably higher than at the NEs of those neighbouring cells on the same coverslip that had remained non-transfected (KO-) and lacked mCherry fluorescence (yellow arrows). Bar, 10 µm.

Since we initially could not tell whether an sfGFP tag might affect TPR’s properties in the one or other manner, when positioned at either the protein’s amino-terminus (NT) or carboxy-terminus (CT), we created both versions of such ZC3HC1 KO cell lines in HeLa, i.e. such expressing only TPR-sfGFP (Supplemental Fig S1) and others only sfGFP-TPR (Supplemental Fig S2). Among the resulting collections of cell clones were such that steadily harboured large amounts of soluble TPR-sfGFP (Supplemental Fig S1) as well as such with similarly large amounts of soluble sfGFP-TPR (see further below). Having then used a TPR-sfGFP-expressing as well as an sfGFP-TPR-expressing cell line, both with a similarly large soluble pool of TPR, for experiments of the kind presented further below in Figure 1, and after having found both cell lines leading to very similar results, the representative data presented in the following stem from the TPR-sfGFP-expressing ZC3HC1 KO cell line only.

In order to allow for induced ectopic expression of ZC3HC1 in such ZC3HC1 KO cells harbouring the soluble pool of sfGFP-tagged TPR, we made use of the Tet-On system (Gossen & Bujard, 1992; Gossen *et al*, 1995) and cloned each one of two ORFs, encoding for different versions of ZC3HC1, into a bidirectional mammalian expression vector derived from the pTetOne expression vector (Heinz *et al*, 2011; Supplemental Fig S3). This setup allowed for constitutive expression of the Tet-On 3G transactivator and the doxycycline-inducible expression of each of the two ZC3HC1 versions.

The one ZC3HC1 expression vector encoded for the intact full-length WT protein. By contrast, the other one, representing a second control next to the common use of the “empty” vector, would express a ZC3HC1 mutant carrying a single amino acid (aa) substitution mutation (C429S) that we had found abolishing the protein’s ability to interact with TPR (Supplemental Fig S4). Both of these ZC3HC1 variants were tagged with mCherry to allow for live-cell imaging and later also for distinguishing between native and recombinant ZC3HC1 when inspecting mixed populations of transfected and non-transfected WT and KO cells that had been grown together on one coverslip (see below).

Next, we cell cycle-synchronised the KO cells and transiently transfected them with the plasmids in G2 in order to subsequently allow for nuclear uptake of the vector primarily during post-mitotic nuclear reassembly and commencement of the constitutive expression of the transactivator from early in G1 on (Fig 1A, and data not shown). Addition of doxycycline, which would then enable the transactivator to initiate transcription of the ZC3HC1 transgenes, followed later in interphase, at about 11 hours after mitosis (Fig 1A), and from then on, the fate of the nuclear pool of TPR-sfGFP, in the presence of then steadily increasing amounts of mCherry-ZC3HC1, was analysed by fluorescence microscopy of live cells at different time points (Fig 1B and 1C).

When inspecting those cells that had been transfected with either the “empty” expression vector (data not shown) or with the vector encoding the NE-binding-incompetent mutant version of ZC3HC1, we found the nucleoplasmic pool of TPR unaffected, with this being evident for all time points investigated. Moreover, even in those cells that eventually harboured conspicuous amounts of the C429S mutant, there was no increase in TPR signal intensities noticeable at their NEs (Fig 1B).

By contrast, when studying those cells that had been expressing the intact version of ZC3HC1, we found the gradually intensifying appearance of mCherry-ZC3HC1 at the KO cells’ NE, once mCherry had become detectable as a fluorescent protein (see below), to come along with a concomitant increase in the NE-associated TPR signal intensities. The latter had been accompanied by a seemingly steady diminishment of the nucleoplasmic pool of TPR, which eventually reached a point at which the pool of soluble TPR initially distributed throughout the nuclear interior was no longer visible (Fig 1C). Furthermore, in many of the transfected cells, all TPR pool 2 polypeptides were found attached to the NE already as early as between about 30 minutes and less than one hour after the addition of doxycycline, which was thus often faster than the time required in eukaryotic cells for the multistep maturation process that turns a newly synthesised mCherry polypeptide into a fluorescent protein (Merzlyak *et al*, 2007; Khmelinskii *et al*, 2012). Moreover, we estimated that during this short time span between induction of ZC3HC1 synthesis and image acquisition, only 2.5% to 5% of the cells’ total amount of TPR would have been newly synthesised (Supplemental Information 2). This, in turn, meant that the majority of TPR polypeptides recruited back to the NE had indeed been such that had already been around as soluble proteins before the synthesis of the new ZC3HC1 polypeptides.

Furthermore, the same type of experiments also performed with a TPR-sfGFP-expressing HCT116 cell line in which all *ZC3HC1* alleles had been knocked out yielded essentially identical results. In these cells too, we found all of the initially soluble TPR pool 2 polypeptides recruited back to the NEs of the transfected cells shortly after having induced expression of the WT ZC3HC1 protein, while in those cells only expressing the C429S mutant protein, these nucleoplasmic TPR polypeptides had remained soluble (our unpublished data).

However, even though rapid synthesis of high amounts of ZC3HC1 was not unrealistic, given that expression vectors often end up within a transfected cell’s nucleus in high numbers (Cohen *et al*, 2009; Glover *et al*, 2010), with imaginable theoretical ZC3HC1 copy numbers exceeding those of the TPR pool 2 polypeptides already after a short time (Supplemental Information 2), one question still needed to be addressed experimentally at this point.

So far, we could not tell whether the depletion of a nuclear pool of soluble TPR and its reattachment to the NE had been paralleled by ZC3HC1 having been attached to the same NEs in amounts proportional to those of TPR. In other words, it remained to be inspected whether NE-binding of the soluble TPR-sfGFP and mCherry-ZC3HC1 polypeptides might eventually reach a state in which both proteins’ total amounts at the ZC3HC1 KO cells’ NEs would be similar to those at the NEs of the WT cells, which would point at similar copy number relationships.

To address this question, we repeated the experiments described in Figure 1A and 1C, yet this time started with a mixed population of cells, namely the TPR-sfGFP-expressing HeLa ZC3HC1 KO cells together with HeLa WT cells naturally expressing the non-tagged TPR and non-tagged ZC3HC1 polypeptides. Such a mixed population was then transfected with the expression vector encoding for mCherry-ZC3HC1. Since the cells were never all transfected, this allowed for directly comparing the amounts of endogenous ZC3HC1 at the NEs of those WT cells that had remained non-transfected with the amounts of mCherry-ZC3HC1 at the NEs of those transfected ZC3HC1 KO cells in which all nuclear TPR-sfGFP had been reattached to the NE. While such comparison of ZC3HC1 amounts in transfected KO cells and non-transfected WT cells was via immunolabelling with ZC3HC1 antibodies, the mCherry-tagged ZC3HC1 polypeptides in the transfected cells were additionally labelled with mCherry-specific single domain antibodies (sdAb), to allow for detecting both the matured and still immature mCherry-tags within the specimens. Consequently, the use of such sdAbs, in combination with the KO cells being identifiable via their GFP-tagged TPR polypeptides, allowed for better identification of a transfected KO cell and for distinguishing it from both non-transfected and transfected WT cells.

This setup then allowed for unveiling that in the transfected KO cells, in which the pre-existing nuclear pool of sfGFP-TPR had been gradually and then quantitatively reattached to the NE, the NE-associated amounts of mCherry-ZC3HC1 had indeed risen concomitantly too. Moreover, in some of the transfected KO cells, the amounts of sfGFP-TPR eventually located at the NE even appeared doubled (Fig 1D), with these KO cells likely representing those in which the initially soluble pool, and thus the total amount of TPR, had been a bit larger than the population’s mean cellular amount. Especially noteworthy then, such quantitative re-recruitment of soluble sfGFP-TPR to the NE had been accompanied by the NE-association of mCherry-ZC3HC1 in amounts that eventually appeared indistinguishable from those of the non-tagged ZC3HC1 present at the NEs of the non-transfected WT cells (Fig 1D; see also Supplemental Information 2).

The outcome of these re-recruitment experiments *in vivo*, and some of the conclusions that could be drawn therefrom, could be summarised as follows. First, the actual re-recruitment of the soluble TPR polypeptides occurred seemingly proportional to NE-association of the recombinant mCherry-ZC3HC3 polypeptides, eventually resulting in some of the cells being in a steady state in which relative and absolute copy numbers of both proteins at the KO cell’s NE were similar to those of the endogenous proteins in the WT cell. Secondly, we could now essentially rule out a scenario in which ZC3HC1-dependent incorporation of TPR polypeptides would only be possible during the formation of entirely new NBs, with these impossibly assembled in large numbers in only 30–60 minutes. Thirdly, it was now clear that the NE-appendage of the ZC3HC1-dependent TPR pool 2 polypeptides could be temporally uncoupled from the ZC3HC1-independent NPC association of TPR. And finally, these findings meant that the synthesis of ZC3HC1 and TPR polypeptides also did not need to occur at the same time, proving that any potential early interaction between ZC3HC1 and TPR in the cytoplasm could at least be ruled out as being a prerequisite for their later appendage to the NB.

However, while now considered proven that ZC3HC1 polypeptides are directly involved in a process in which TPR polypeptides are recruited from a soluble nuclear pool and appended to those TPR pool 1 polypeptides that are already anchored at the NPCs in a ZC3HC1-independent manner, it still remained unknown at this point whether localisation of ZC3HC1 at the NB was due to direct interaction with the ZC3HC1-independent TPR pool 1 first, then followed by the recruitment of the second TPR pool, or whether ZC3HC1 was merely co-recruited together with the TPR pool 2 polypeptides. In this latter case, one could envision scenarios in which only the binding to such pool 2 polypeptides would allow ZC3HC1, once it had entered the nucleus, to adopt a conformation that would enable it to bind to TPR pool 1 polypeptides. Alternatively, it even appeared imaginable that binding to a ZC3HC1 polypeptide in solution would allow the TPR pool 2 polypeptides to adopt and maintain some distinct conformation, which only then would render them capable of interacting with the NPC-anchored TPR polypeptides.

In fact, we knew that soluble forms of ZC3HC1 and TPR can, in principle, be co-immunoprecipitated together (Gunkel *et al*, 2021), but we did not know whether such an interaction in solution would be a requisite for the two proteins’ subsequent binding to the NB. In other words, we did not know whether ZC3HC1 and the T2 polypeptides would need to bind concomitantly to the NB or whether such binding could also happen in a stepwise manner. Therefore, we set out to investigate next whether the binding of the ZC3HC1 and the TPR pool 2 polypeptides to the already NPC-anchored TPR pool 1 polypeptides could be uncoupled from each other.

### ZC3HC1 first bound to the NPC-anchored TPR pool 1 *in vitro*, allows for subsequent attraction and appendage of additional TPR pool 2 polypeptides

Addressing the topic of how ZC3HC1 contributes to the approximate doubling of the NE-associated TPR amounts had thus gone along with raising the following questions: Can ZC3HC1 bind directly to the NPC-anchored, ZC3HC1-independent pool 1 of TPR polypeptides, independently of the ZC3HC1-dependent soluble pool 2 of TPR? And, if this were the case, would these ZC3HC1 polypeptides, once bound to the ZC3HC1-independent pool of TPR, allow for the attraction of additional TPR polypeptides from a soluble second pool, as one might then expect?

Pursuing different *in vivo* and *in vitro* approaches to address the first question, namely whether ZC3HC1 on its own can bind to TPR pool 1, i.e. without any need for TPR pool 2 polypeptides being around, we eventually found that the different strategies all arrived at the same result. Since one of these experimental setups also allowed addressing some of the other still open questions in the same manner, only this particular approach is presented in the following.

The starting point for such kind of experiment were HeLa ZC3HC1 KO cells that had been grown in slide wells, normally used for live-cell imaging, to which they tightly adhered during all subsequent treatments. These cells, expressing TPR in its non-tagged native form, were detergent-permeabilised and subsequently washed in such a manner that even traces of soluble TPR pool 2 polypeptides were quantitatively removed, resulting in the NPC-lamina scaffold of the NEs (Supplemental Fig S5A) that were then only possessing the NPC-associated TPR pool 1 polypeptides. The latter could then be used as a platform that exhibited only minor residual autofluorescence in the 520–530 nm wavelength range and essentially none between 540–590 nm, and onto which one would then add, in the one or other combination, soluble mCherry-tagged ZC3HC1 polypeptides that had been obtained from other cells (Fig 2A).

**Figure 2.**
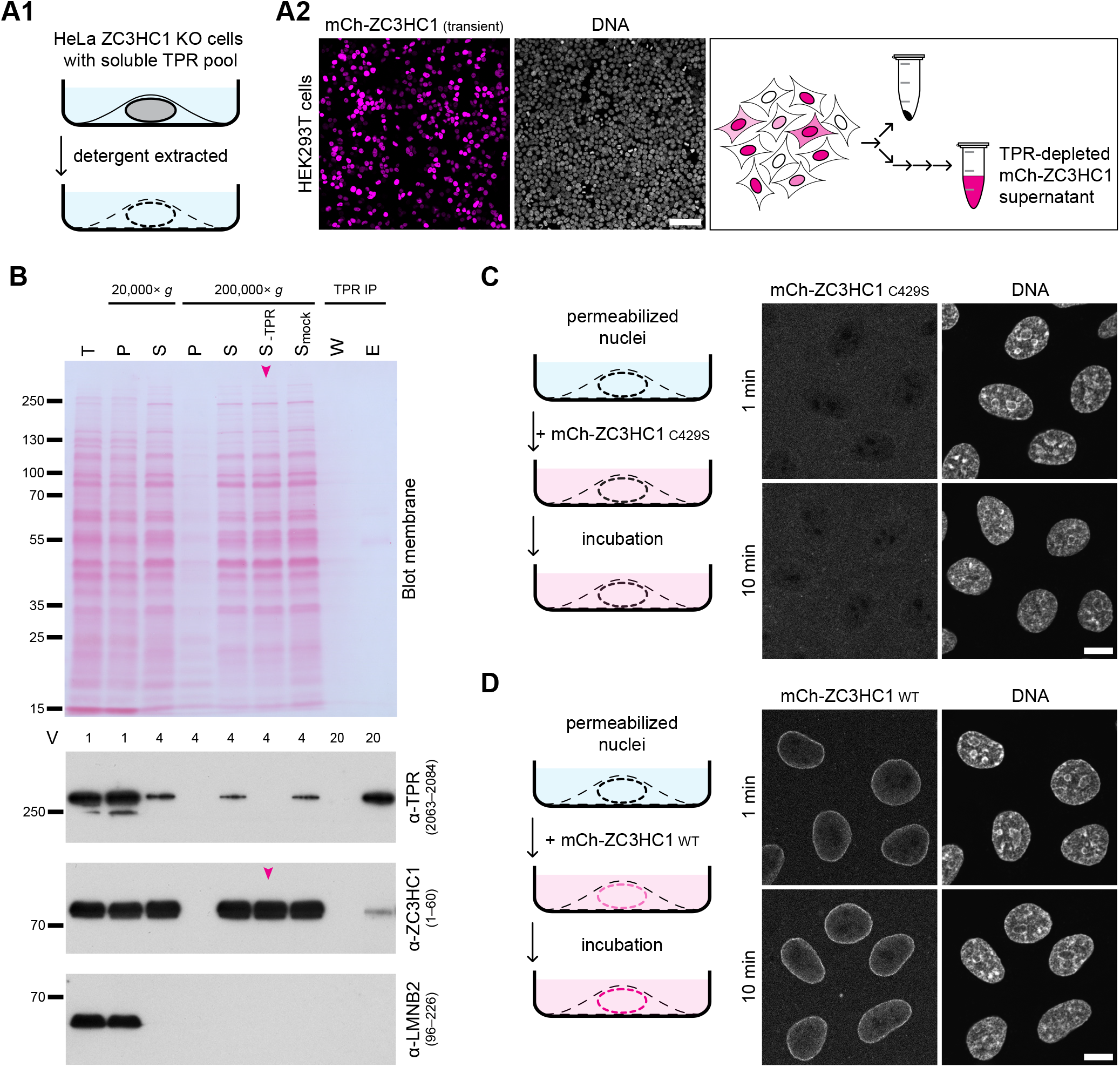
Ectopically expressed ZC3HC1 can bind to the ZC3HC1-independent pool of NPC-anchored TPR polypeptides *in vitro*. **(A)** Depiction of experimental materials and starting conditions, comprising (A1) HeLa ZC3HC1 KO cells permeabilised with TX-100 and cleared of soluble TPR polypeptides (for additional information, see Supplemental Fig S5), and (A2) supernatants of HEK293T cell extracts containing ectopically expressed variants of mCherry-ZC3HC1, obtained after cell permeabilisation with elevated concentrations of digitonin, high-speed centrifugation, and immunodepletion of TPR. The micrographs in 2A2 show representative live-cell images of the transiently transfected cell population schematically depicted on the right. Bar, 100 µm. **(B)** IBs of a selection of the cellular fractions obtained from the HEK293T cells that had been ectopically expressing mCherry-ZC3HC1. Loaded fractions included the total cell extract (T) and the digitonin-containing soluble cell extracts, obtained after 20,000× *g* and 200,000× *g* centrifugation, which contained minor amounts of soluble TPR (S 20,000 and S 200,000), the occurrence of which represented a common finding for HEK293T cells ectopically expressing large amounts of recombinant ZC3HC1. Furthermore, the loadings also included the corresponding pellets of these centrifugation steps (P 20,000 and P 200,000), the 200,000× *g* cell extract supernatant mock-depleted with magnetic beads only coated with Protein A (S_mock_ 200,000) and the same supernatant TPR-depleted with Protein A beads coated with *Hs*TPR antibodies (S_-TPR_ 200,000). Such latter supernatant cleared of TPR is here marked with an arrowhead and represented the mCherry-ZC3HC1-containing cell extract eventually used for the interaction experiments. For further comparison, additional lanes were loaded with materials released from the TPR beads during the second of two successive washing steps (W) and those finally eluted from these beads with SDS-containing sample buffer (E). Most lanes were loaded with the respective amount from the same number of HEK293T cells (4 V), while relative amounts of the total cell extract and the 20,000× *g* pellet material corresponded only to one forth thereof (1 V), whereas the W and TPR-IP fractions again represented five-fold higher relative amounts (20 V). Incubations with *Hs*TPR and *Hs*ZC3HC1 antibodies, and with LMNB antibodies for comparison, were on different parts of the Ponceau S-stained membrane shown here and on another one with identical amounts loaded. Note that essentially no TPR was detectable within the 200,000× *g* supernatant after the TPR depletion procedure. As an aside, also note that the fraction of immunodepleted TPR contained co-immunoprecipitated mCherry-ZC3HC1, which underscored the necessity to remove the minor pool of soluble TPR from the 200,000× *g* supernatant to avoid the experiment’s conclusiveness being spoiled by soluble TPR-ZC3HC1 assemblies. **(C, D)** Fluorescence microscopy of the detergent-extracted ZC3HC1 KO cells that had been incubated with TPR-free cell extracts, which had been adjusted to containing similar amounts of either, in 2C, the mCherry-tagged C429S mutant version of ZC3HC1 incapable of TPR binding, or, in 2D, the WT ZC3HC1 protein. Interaction experiments here shown as 2C and 2D had then been performed in parallel to each other in neighbouring wells, and images of the unfixed cells, in addition also stained with the DNA dye Hoechst 33342, had been acquired with identical microscope settings, at 1 and 10 min after the addition of cell extracts. Because of the necessity to omit anti-fade media and the fact that focussing on the equatorial plane of the extracted cells was more time-consuming than for intact or fixed cells, images taken at different time points were not from identical groups of cells but only from neighbouring groups of the same sample to avoid pronounced photobleaching. Moreover, reduced laser power, compared to that for IFM images, was chosen for the image acquisition of the *in vitro* assembly experiments, which was later followed by electronic brightness enhancement in the same proportional manner and identical extent for all of these images (see Material and Methods). Note that while no mCherry fluorescence appeared enriched specifically at the NEs treated with the C429S mutant, those NE scaffolds that had been incubated with the WT version for a similarly long time were by then clearly fluorescent. Bars, 10 µm.

The ZC3HC1 polypeptides to be added to these NE platforms in the course of such *in vitro* reconstitution experiments were the full-length intact version and, as a control again, the mutant version C429S incapable of TPR interaction. These two ZC3HC1 variants had been ectopically expressed in HEK293T cells, and from these, cell extracts were prepared that were essentially free of TPR. To this end, we applied a relatively gentle extraction procedure, using digitonin as the only detergent. With digitonin nonetheless allowing for NE perforation, when used at higher concentrations than those sufficient for plasma membrane permeabilisation, large enough amounts of the recombinant ZC3HC1 polypeptides, primarily located within the HEK293T cells’ nuclei, could be released from there, together with only minor amounts of soluble TPR (Fig 2B). However, even these relatively few TPR polypeptides, usually not detectable in non-transfected HEK293T cells and when present likely a consequence of ZC3HC1 overexpression (see also Gunkel *et al*, 2021), needed to be quantitatively removed in order to fulfil the requirements for this experimental setup and allow it to be sufficiently informative. Such clearance was achieved via immunodepletion, using a mixture of specific *Hs*TPR antibodies coupled to magnetic beads and eventually resulting in not even traces of TPR being detectable within the cell extracts anymore (Fig 2B).

Furthermore, even though the resulting concentrations of the mCherry-tagged WT and C429S mutant proteins were recurrently found to differ to some extent within the final cell extracts, a finding that was taken into consideration when later performing the actual experiments, they exceeded the concentrations of the soluble endogenous ZC3HC1 within these cell extracts by far (Supplemental Fig S5B and S5C; for approximations of intracellular concentrations, see Supplemental Information 3).

These TPR-free extracts were then separately added to each one of the slide wells with the NE scaffolds devoid of endogenous ZC3HC1 and TPR pool 2 polypeptides. The specimens were then immediately inspected by fluorescence microscopy and then further monitored for a certain period of time. Those to which the ZC3HC1 mutant C429S had been added did not show signs of mCherry fluorescence having been enriched specifically at the NE scaffolds within a predefined time span (Fig 2C). Unspecific binding to various subcellular structures, never observed with the mCherry-tagged WT ZC3HC1 polypeptides, was only observed after prolonged incubation (data not shown, but see also Supplemental Information 3).

By striking contrast, within seconds after adding the WT version of ZC3HC1 to a parallel batch of such permeabilised cells, mCherry fluorescence was noted specifically accumulating at the NEs, and after a few minutes, such NE-associated fluorescence had reached a seemingly constant brightness, indicative of a steady-state level of interactions between the ZC3HC1 polypeptides and some definite number of ZC3HC1 binding sites provided by the TPR pool 1 polypeptides (Fig 2D). Moreover, while the extracts for the experiments presented here had been adjusted to contain similar concentrations of either the mCherry-tagged WT or the C429S mutant version of ZC3HC1, essentially identical results were also obtained when using these proteins in a range of other concentrations too (data not shown).

Furthermore, upon washing these NEs with a detergent-free NB-stabilising (NB-s) buffer, in the following called assembly buffer, which allowed for removing all unbound mCherry fluorescence from the well, the NE-associated mCherry-ZC3HC1 was not similarly removed, further pointing at some ZC3HC1:TPR pool 1 interactions that could last for some time even in the absence of TPR pool 2 polypeptides (Supplemental Fig S6). However, we also noted that these ZC3HC1 interactions with the NE platforms only possessing TPR pool 1 were not for all of the appended mCherry-ZC3HC1 polypeptides similarly long-lasting. Some of the those initially appended to the NE scaffolds were found swiftly detaching when washing these NEs in assembly buffer, eventually resulting in only about half of the initially NE-appended amount of ZC3HC1 durably remaining bound to the TPR pool 1 polypeptides (Supplemental Fig S6). Such detachment of a certain amount of mCherry-ZC3HC1 differed from what one could observe with similarly detergent-extracted HeLa WT cells harbouring endogenous ZC3HC1 and both pool 1 and pool 2 of TPR. When such detergent-extracted WT cells were kept in the same assembly buffer, the NE-association of ZC3HC1, as part of the likely more complex assemblies involving ZC3HC1 and both pools of TPR, was not notably affected by similarly long incubations (our unpublished data, but see also further below).

Nonetheless, having demonstrated that ZC3HC1 can bind to the NE entirely independent of the TPR pool 2 polypeptides, we then asked whether these bound ZC3HC1 polypeptides could next allow for the recruitment of additional TPR polypeptides and for keeping them tethered there at the NE.

To this end, we harvested cell extracts from those HeLa ZC3HC1 KO cell lines that were expressing TPR only as sfGFP-tagged polypeptides and in which a major proportion of these proteins was occurring distributed throughout the nuclear interior as a soluble pool (Fig 3A), with us showing here the results with the extracts from the HeLa ZC3HC1 KO cell line in which the TPR polypeptides occur N-terminally tagged with sfGFP (Supplemental Fig S2). The resulting extract after high-speed centrifugation (Fig 3B), free of ZC3HC1 and NE fragments but containing large amounts of exclusively soluble sfGFP-tagged TPR (see also Supplemental Information 3), was then used for the following experiments.

**Figure 3.**
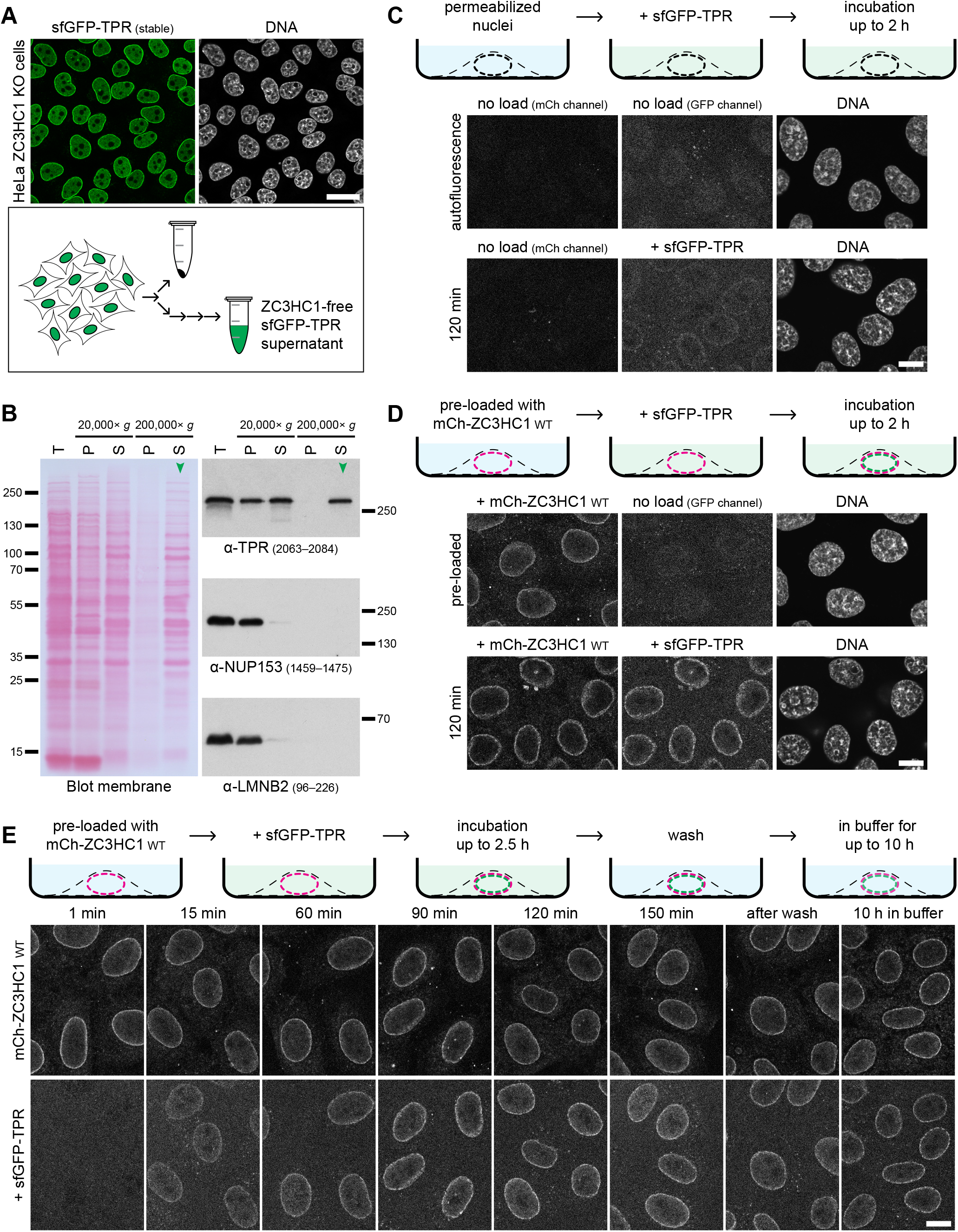
Ectopically expressed ZC3HC1 loaded onto the ZC3HC1-independent pool of NPC-anchored TPR can subsequently attract additionally provided TPR polypeptides, resulting in both types of proteins steadily appended to the NE. **(A)** Representative live-cell image of the sfGFP-TPR-expressing HeLa ZC3HC1 KO cell line, and schematic depiction of the high-speed supernatant of a detergent-free extract isolated from such cells, containing the soluble pool 2 polypeptides of sfGFP-TPR. Bar 25 µm. **(B)** IBs of a selection of the cellular fractions obtained from the sfGFP-TPR-expressing ZC3HC1 KO cells. Loaded fractions included the total cell extract (T), the soluble extracts obtained after the disruption of the cells by gentle sonication and subsequent centrifugation at 20,000× *g* (S 20,000) and 200,000× *g* (S 200,000), and the corresponding pellet fractions (P 20,000 and P 200,000). Lanes were loaded with amounts corresponding to the same number of HeLa ZC3HC1 KO cells. Immunolabellings for TPR, for the NPC component NUP153, and for the nuclear lamina component LMNB, the two latter proteins immunolabelled for comparison, were on different parts of the Ponceau S-stained membrane shown here and on an identically loaded duplicate. Note that the 200,000× *g* supernatant, marked with an arrowhead and representing the cell extract with the soluble sfGFP-TPR that we categorised as pool 2 polypeptides and eventually used for the interaction experiments, was virtually free of LMNB and NUP153. This finding indicated that the solution neither contained tiny NE and NPC fragments, as might have been suspected a consequence of the sonication process nor considerable amounts of NUP153, as the only other so far known TPR-binding protein that can mediate binding of TPR to distinct subpopulations of NPCs. **(C)** Fluorescence microscopy of the detergent-extracted ZC3HC1 KO cells in the complete absence of ZC3HC1, both before (no load) and after having then been incubated with the ZC3HC1-free but sfGFP-TPR-containing cell extract at RT (120 min). Note that the images of the not loaded cells labelled “mCh channel” and “GFP channel” actually reflected the degree of autofluorescence of detergent-extracted HeLa cells at different wavelengths. Note also that with the microscope settings chosen, which were kept the same also for corresponding images in 3D and 3E, essentially no autofluorescence was detected at the mCherry excitation wavelengths, while only some autofluorescence was notable at wavelengths that would excite GFP. Further note that after incubations of up to 120 min with the sfGFP-TPR extract, there was hardly any sfGFP fluorescence at some of the ZC3HC1-deficient NEs, or it was only marginally enhanced at others, beyond the autofluorescence background in this wavelength range. Bar, 10 µm. **(D)** Fluorescence microscopy of NE scaffolds like in C but first loaded with the mCherry-tagged WT version of *Hs*ZC3HC1, followed by the removal of the unbound surplus of mCherry-ZC3HC1 by brief washes with assembly buffer (pre-loaded specimen). Soluble sfGFP-TPR polypeptides had been added immediately after that, using the same sfGFP-TPR aliquot as for 3C, with interaction experiments shown as 3C and 3D performed almost simultaneously in parallel to each other in neighbouring wells, with the double-loaded specimens shown here also imaged after 120 min. Note that the sfGFP-tagged TPR polypeptides had specifically and conspicuously accumulated at the NE scaffolds that had been pre-loaded with mCherry-ZC3HC1. As an aside, it also needs to be mentioned that sfGFP-TPR signal intensities at the NEs of such permeabilised cells would not have been able to reach the levels in a WT cell expressing all pool 1 and pool 2 TPR polypeptides as tagged with sfGFP, simply because all pool 1 TPR polypeptides within the ZC3HC1 KO cells, here used as binding platforms, were untagged. Bar, 10 µm. **(E)** Fluorescence microscopy of initially ZC3HC1-deficient NE scaffolds that had been loaded with mCherry-ZC3HC1 first, then briefly washed in assembly buffer, immediately incubated in an sfGFP-TPR-containing cell extract for 150 min, again washed with buffer, and subsequently kept within such buffer for at least 10 h. Note that in this experiment, distinct from the one in 3D, the specimens were inspected at different time points, revealing gradual accumulation of the sfGFP-TPR polypeptides at the mCherry-ZC3HC1-loaded NEs until seemingly reaching a steady-state level of TPR at such NEs. Note further that such binding appeared to be rather persistent after having removed the cell extract, with only moderate reduction of the NE-associated sfGFP and mCherry fluorescence notable, even after having kept the specimens in cell extract-free buffer at RT overnight. Bar, 10 µm.

First, however, knowing that especially GFP fluorescence in cultured cells can be contaminated to some extent by cellular autofluorescence (Niswender *et al*, 1995), we regarded it as obligatory to assess the threshold beyond which GFP signals can be discriminated from autofluorescence. For this reason, we had inspected the detergent-extracted HeLa ZC3HC1 KO cells only in assembly buffer and in the absence of the fluorescent protein-containing cell extracts for the relative levels of residual autofluorescence at different wavelengths. Microscope settings, subsequently chosen for obtaining the original micrographs, and the degree of subsequent electronic signal enhancement by an identical multiplication factor, were then kept the same for the corresponding images here presented as Figure 3C and3D. With these settings, essentially no autofluorescence was detected in the original micrographs representing the mCherry excitation wavelengths, while some autofluorescence had to be taken into account as background at wavelengths that would excite GFP (Fig 3C).

When the NE scaffolds devoid of ZC3HC1 and TPR pool 2 polypeptides were then incubated with the sfGFP-tagged polypeptides only, this did not result in any pronounced accumulation of these polypeptides specifically at the ZC3HC1-deficient NEs, with hardly any or no sfGFP fluorescence detectable at some of these NEs, while at other NEs, the sfGFP levels turned out to be only marginally enhanced, relative to the autofluorescence (Fig 3C). Whether this reflected some weakish interactions between the sfGFP-tagged TPR polypeptides and other NE proteins, or perhaps even some weak direct interactions between the different TPR populations, could not be resolved in the current study.

Clearly, however, when the NE scaffolds had already been loaded with mCherry-ZC3HC1 beforehand, meaning that such ZC3HC1 had first been allowed to interact with the available TPR pool 1 binding sites, followed by removing the non-bound ZC3HC1 surplus and briefly washing these NE scaffolds, and only then incubating them with the sfGFP-tagged TPR polypeptides, the latter were found markedly accumulated at the NE (Fig 3D).

In this context, we do not want to miss out on mentioning that we also noted in the course of repeating such experiments, for corroborating data reproducibility, some variability in the amounts of sfGFP-TPR that we finally found appended to such mCherry-ZC3HC1-loaded NEs. While the marked accumulation of sfGFP-TPR only at the mCherry-ZC3HC1-loaded NEs, in comparison to those not loaded with mCherry-ZC3HC1, was evident in all experiments, the NE-associated GFP signal intensities finally observed were sometimes found to differ. To some extent, such variability appeared to reflect an inverse correlation between (i) the length for which the ZC3HC1-loaded NE scaffolds had been washed before loading the sfGFP-tagged polypeptides and (ii) the amounts of sfGFP-TPR subsequently bound to these NEs. However, while perhaps pointing at different subpopulations of NB-bound mCherry-ZC3HC1 polypeptides (see also Supplemental Fig S6) being both required for the recruitment and anchorage of the TPR pool 2 polypeptides, a systematic, thoroughly controlled analysis of this possibility will need to be a topic of future research.

In the current study, though, we could already conclude that without ZC3HC1 first appended to the pool 1 polypeptides of TPR, a subsequent lasting appendage of TPR pool 2 polypeptides in considerable amounts could not possibly occur in this setting. In fact, when (i) mCherry-ZC3HC1-loaded NEs that had then been briefly but nonetheless efficiently washed, and (ii) corresponding NEs that had not been loaded with mCherry-ZC3HC1, had been incubated with the same sfGFP-TPR extract in parallel, i.e. in wells side by side as had been done for the specimens presented in Figure 3C and 3D, only the ZC3HC1-loaded ones were found capable of reproducibly attracting the TPR polypeptides in substantial amounts.

Attempting also to assess how lasting and robust these *in vitro* interactions might be, now between both pools of TPR and ZC3HC1, we performed further experiments in which the mCherry-ZC3HC1-loaded NE scaffolds, once they had been allowed to attract steady-state levels of sfGFP-TPR, were again washed with buffer and then kept therein for many hours (Fig 3E). While fluorescence was sometimes found very moderately reduced after prolonged dwelling times, the NE-associated levels of sfGFP- and mCherry-fluorescence were nonetheless recurrently found still prominent, and then indistinguishable from the same experiment’s not yet washed sfGFP-TPR-loaded NEs, even after 10–12 hours of incubation in the cell extract-free buffer at RT (Fig 3E, and data not shown). This finding suggested that the interplay between the ZC3HC1 polypeptides and the pool 1 and 2 sfGFP-TPR polypeptides had resulted in an arrangement that allowed for a lasting positioning at the NE.

Altogether, we interpreted these *in vitro* data in such a way that the presence of ZC3HC1 polypeptides, once attached to the NPC-appended pool 1 of TPR, allows for the subsequent recruitment of further TPR pool 2 polypeptides and promotes their interaction with those of pool 1. Beyond that, we even felt tempted to already reckon ZC3HC1 as a structural element that would at least stabilise the interactions between the different TPR populations at the NB or even function as a direct interconnector between them.

In actuality, however, we even then could not yet exclude another scenario reminiscent of the relationship between the NPC-anchored TPR polypeptides and the NPC protein NUP153, as will be outlined further below. ZC3HC1, in such a scenario, would enable the establishment of some higher-order arrangements between the different TPR populations, similar to NUP153 enabling the NPC appendage of TPR subpopulations, but then, just like NUP153, be no longer required for maintaining the integrity of such structures, once they had been assembled. This possibility thus still needed to be addressed next.

### ZC3HC1 *in vivo* is directly required for keeping TPR subpopulations attached durably to those TPR polypeptides have been anchored at the NPC independently of ZC3HC1

In the absence of unequivocal evidence for whether or not ZC3HC1 might also be required for maintaining some specific arrangements between TPR polypeptides at the NB, we could not yet declare ZC3HC1 beyond doubt also a construction element of the NB. Such uncertainty was due to the circumstance that formerly observed phenotypes, such as the appearance of soluble TPR pools following the destruction of the ZC3HC1 transcripts by RNAi or the CRISPR/Cas9n-mediated disruption of the *ZC3HC1* alleles (Gunkel *et al*, 2021), had only manifested themselves time-delayed. Namely, either after the RNAi-mediated transcript elimination had at some point eventually resulted in notably diminished cellular amounts of ZC3HC1 or after having accomplished the unavoidably lengthy process of identifying and culturing ZC3HC1 KO cell clones (Gunkel *et al*, 2021), which in both cases meant that such cells had to pass through mitosis before manifesting a phenotype. However, since all NBs are disassembled at the outset of mitosis and then need to be reassembled again from soluble components in the daughter cells, it had not been possible to unambiguously distinguish between a direct or indirect role of ZC3HC1 in either the recruitment or appendage of the TPR pool 2 polypeptides or in keeping them bound to the NB.

Therefore, even though it had been tempting to imagine that ZC3HC1 might be a protein with some structural role at the NB, we were aware that we lacked experimental evidence that would have allowed for direct and immediate correlations between the presence or absence of ZC3HC1 on the one hand and subcellular positioning of the ZC3HC1-dependent pool of NB-associated TPR on the other. And without such evidence, it was not possible to unequivocally exclude scenarios in which the formerly observed ZC3HC1 deficiency phenotypes would merely reflect some indirect contributions of ZC3HC1 to the one or other process leading to TPR pool 2 polypeptides being eventually positioned at the NB.

While the recruitment and appendage experiments here presented as Figures 1 to 3 had then allowed for a close spatiotemporal correlation between the presence of ZC3HC1 and resulting NB-positioning of TPR pool 2, we still needed to converge to a similarly conclusive spatiotemporal correlation between ZC3HC1 deficiency and resulting phenotypes with regard to the fate of the ZC3HC1-dependent pool of TPR, once it had been appended to the NB.

Since we regarded degron-based technologies as possibly best suitable for achieving elimination of ZC3HC1 in such a rapid manner that the cells would no longer need to pass through mitosis before phenotypes could manifest themselves, we created yet further CRISPR/Cas9-edited cell lines. One of them was meant for the inducible degradation of the then degron-tagged ZC3HC1 polypeptides, and the other, for comparison, would similarly allow for degradation of TPR.

To this end, we created a versatile insertion cassette, encoding for a double-tag comprising an auxin-inducible degron (AID)-tag (Li *et al*, 2019) and sfGFP. The former, we positioned within a loop of sfGFP in which we had found this AID-tag (auxin-responsive protein IAA7, aa37-104, in the following called mIAA7) to be fully functional as a degron while only marginally attenuating sfGFP’s ability to fluoresce (Supplemental Fig S7A, our unpublished data, and further below).

In addition, we had integrated another cassette, which allowed for constitutive expression of the auxin signalling F-box 2 protein AFB2 gene from *Arabidopsis thaliana* (Li *et al*, 2019), into the genome of the HCT116 cell line. Apart from being nearly diploid and having used it as one of several representative cell lines in the prequel study on ZC3HC1 (Gunkel *et al*, 2021), we chose HCT116 as the first cell line for performing such degron-induced degradation experiments also because it already had been proven suitable for AID-tagging and auxin-inducible target degradation (Natsume *et al*, 2016).

As the integration site for the *At*AFB2 cassette, we had chosen the adeno-associated virus integration site 1 (AAVS1) safe-harbour locus (Supplemental Fig S7B) and eventually isolated an *At*AFB2-expressing HCT116 clone that we found suitable as the master cell line (MCL, Supplemental Fig S7C) for subsequently integrating the cassette with the AID-tag within loop 9 of sfGFP (in the following referred to as sfGFP^L9mIAA7^) into the alleles of any target gene of interest.

Using the sfGFP^L9mIAA7^ cassette to target the *ZC3HC1* and *TPR* alleles in this HCT116 MCL, we eventually isolated homozygous cell lines in which either all TPR (Supplemental Fig S8) or all ZC3HC1 polypeptides (Supplemental Fig S9) possessed such a sfGFP^L9mIAA7^-tag. Furthermore, when comparing by immunofluorescence microscopy (IFM) the NE-attached amounts of the tagged polypeptides with those of the corresponding tag-free polypeptides appended to the progenitor cell line’s NEs, these amounts appeared very similar, if not seemingly indistinguishable, in most cells (Figs 4A1 and 5A1), indicating that this tag still allowed for largely quantitative NB-association of both TPR and ZC3HC1. While minor differences in the total amounts, occasionally observed after having tagged the 5’-end of some genes, including *TPR*, were considered in the context of another study in which we determined the corresponding proteins’ copy numbers at the NPC and NB (our unpublished data), we found such minor variations not notably affecting the outcome of the here presented degradation experiments.

Next, we triggered TPR degradation in the sfGFP^L9mIAA7^-TPR cell line by adding auxin and thereby initiating *At*AFB2-mediated recruitment of the proteasome to its target, which resulted in essentially all TPR degraded within less than 90 minutes (Supplemental Fig S10A, and further below). In addition, this had come along with ZC3HC1 displaced from the NE and then only present in soluble form within the nucleoplasm (Fig 4). Thus, being in line with the outcome of former TPR RNAi experiments, which had already indicated that TPR represents the only anchor point for ZC3HC1 at the nuclear periphery (Gunkel *et al*, 2021), the current result corroborated this conclusion, confirming that all ZC3HC1 polypeptides appended to the NE are only located there because of their interactions with TPR.

**Figure 4.**
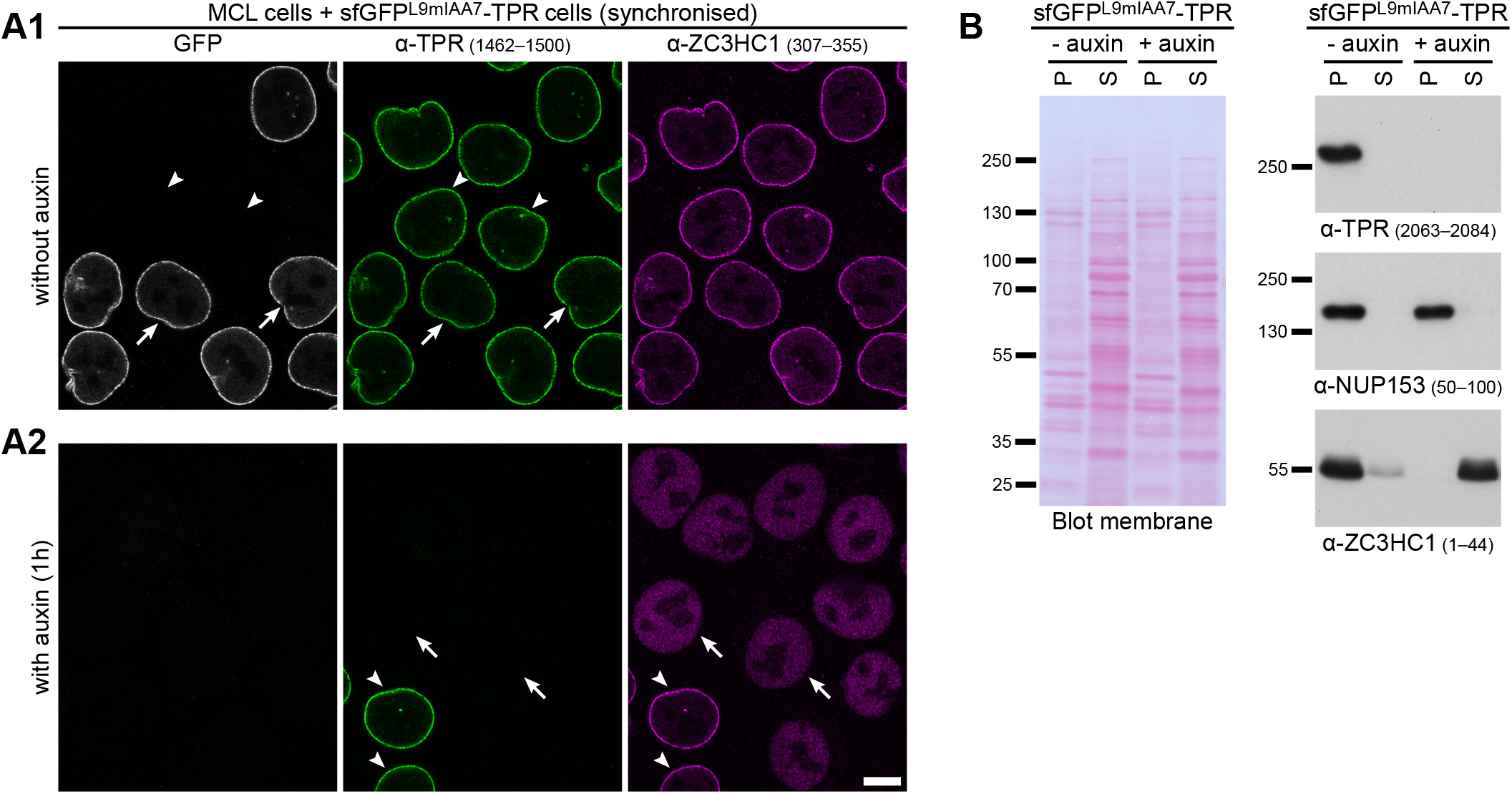
Auxin-induced TPR degradation in a homozygous sfGFP^L9mIAA7^-TPR cell line results in the detachment and solubilisation of NB-bound ZC3HC1. **(A)** IFM of cells from the HCT116 progenitor MCL expressing the naturally tag-free TPR and cells of the homozygous HCT116 progeny line, in which all TPR polypeptides were N-terminally tagged with sfGFP^L9mIAA7^. Cells had been co-cultured and synchronised as mixed populations together on the same coverslip, followed by an additional incubation of one hour in the absence (A1) or presence (A2) of auxin. Specimens were then double-immunolabelled for TPR and ZC3HC1 and analysed in parallel, using identical microscope settings. Arrowheads mark the nuclei of some MCL cells, i.e. those expressing the tag-free version of TPR, while arrows point at some of the progeny cells’ nuclei, i.e. those with the sfGFP-tagged TPR. Note first, in 4A1, that the amounts of sfGFP^L9mIAA7^-tagged TPR appended to the NEs did not notably differ from the amounts in the neighbouring tag-free cells. Further note, in 4A2, that auxin-treatment had resulted in the complete elimination of any visible NE-associated GFP and TPR immunostaining in those cells that had been expressing the tagged version of TPR, while the MCL cells’ untagged TPR polypeptides had remained unaffected. Moreover, note, in particular, that the elimination of the tagged TPR had been accompanied by ZC3HC1 being no longer detectable at the NE but distributed throughout most of the nuclear interior. Bar, 10 µm. **(B)** IB of cell extracts obtained from the sfGFP^L9mIAA7^-TPR HCT116 MCL cells after having treated these with auxin, or only with DMSO, for one hour. Similar numbers of cells of the two differently treated batches had then been fractioned in parallel, resulting each in a fraction of soluble proteins (S) and a corresponding pellet fraction (P) also containing the NPCs and normally, i.e. in the absence of auxin, the complement of NB proteins. Equal portions of each fraction’s whole amount were then loaded for SDS-PAGE and IB. Incubations with *Hs*TPR and *Hs*ZC3HC1 antibodies, and *Hs*NUP153 antibodies for comparison, were on different parts of the Ponceau S-stained membrane shown here and on another one with identical loadings. Note that TPR was no longer detectable after auxin treatment. Moreover, ZC3HC1, mainly part of the LNN-enriched pellet fraction of cells not treated with auxin, had been released into solution upon auxin-induced TPR degradation, while the exclusive presence of NUP153 within the NPC-NB-enriched fraction had remained unaffected.

Of further note, TPR degradation had also resulted in the detachment of other known TPR binding partners from the NE, here exemplified by IFM for MAD1, GANP and SENP1 (Supplemental Fig S11). Strikingly, though, while the detachment of some NB components, like ZC3HC1, was found to correlate strictly temporarily with TPR’s degradation, this was not the case for others, here exemplified by GANP. At early time points, at which essentially all TPR had been degraded, and virtually all ZC3HC1 polypeptides had been concomitantly detached from the NE and released into a soluble nucleoplasmic pool, almost all GANP polypeptides were still found appended to the NE (Supplemental Figure S12), and from there, they then only gradually detached over time. Such degron-based investigations thus allowed for assigning NB-associated proteins to distinct categories, with ZC3HC1 exemplifying those proteins whose NB residency in proliferating culture cells appears to be solely TPR-dependent, and others like GANP that appear to engage in versatile interactions with yet other proteins at the nuclear periphery, with some of the latter even on their own capable of keeping proteins like GANP positioned at the NE for some time. Nonetheless, in the end, all of the proteins so far regarded as ratified NB components, including GANP, had in common that they could not lastingly remain positioned at the NPC when TPR was absent (Supplemental Figure S11; and our unpublished data).

By contrast, proteins formerly proposed as ZC3HC1 binding partners (e.g., Ouyang *et al*, 2003; Bassermann *et al*, 2005, 2007; Klitzing *et al*, 2011; Illert *et al*, 2012; Gengenbacher *et al*, 2019) were not among the proteins initially present and then released from the NE upon TPR degradation (our unpublished data), with this in agreement with recent data (Gunkel *et al*, 2021). Furthermore, some formerly proposed NB components, like, e.g. NUP98, were not found detached even in minor amounts either, and this similarly also held for most of the other nucleoporins (NUPs), i.e. the other proteins of the NPC proper and its cytoplasmic appendices (our unpublished data). Merely for one of TPR’s binding partners at the NPC, namely for NUP153, and NUP50 as one of the other binding partners of NUP153, did we note some minor reduction in their NE-associated amounts after several hour-long incubations following the auxin-induced degradation of TPR (Supplemental Figure S11; our unpublished data), again in line with former studies in which TPR and the NB had been eliminated by different means (e.g., Hase & Cordes, 2003; Aksenova *et al*, 2020; Gunkel *et al*, 2021).

The result regarded as even more informative, though, was obtained after having triggered with auxin the degron-mediated degradation of ZC3HC1, which resulted in essentially complete degradation within less than 90 minutes. Most strikingly, such disappearance of ZC3HC1 was accompanied by roughly about half the total amount of NE-appended TPR being no longer attached to the nuclear periphery (Fig 5A2). Instead, a conspicuous amount of TPR was then found distributed within the nucleoplasm (Fig 5A2).

The more precise quantification of the signal intensities for the residual amounts of immunolabelled TPR at the ZC3HC1-deficient NEs then yielded another finding that we regarded as of note, namely that such a reduction as the result of the induced proteasomal degradation of ZC3HC1 amounted to actually more than half and sometimes up to about 60% of the total NE-associated amounts of TPR present before ZC3HC1 degradation (Fig 5B). This degree of reduction differed moderately, but seemingly reproducibly, from the diminishment by only about 40–50% that one could determine for the NE-associated amounts of TPR after having achieved ZC3HC1 deficiency by RNAi or in ZC3HC1 KO cells of different cell lines, and, in particular, in HCT116 (Gunkel *et al*, 2021; our unpublished data). However, we did not regard these differences as inconsistent with each other, as discussed further below.

**Figure 5.**
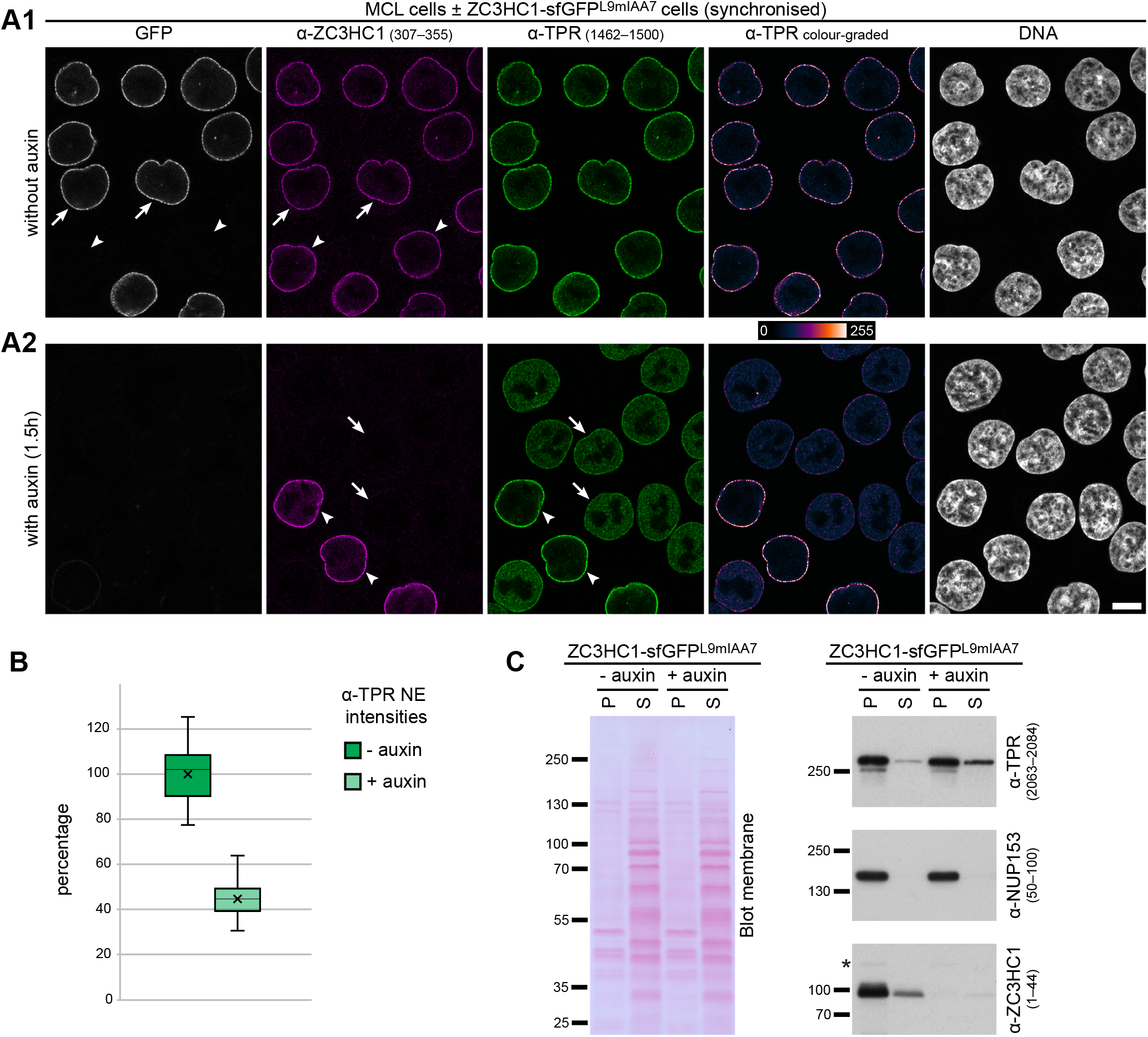
Auxin-induced rapid and quantitative ZC3HC1 degradation in a homozygous ZC3HC1-sfGFP^L9mIAA7^ cell line results in the immediate detachment of large amounts of NB-positioned TPR. **(A)** IFM of cells from the HCT116 progenitor MCL expressing the naturally tag-free ZC3HC1 and cells of the homozygous HCT116 progeny line, in which all ZC3HC1 polypeptides were C-terminally tagged with sfGFP^L9mIAA7^. Cells had been co-cultured and synchronised as mixed populations together on the same coverslip, followed by an additional incubation of 90 min in the absence (A1) or presence (A2) of auxin. Specimens were then double-immunolabelled for ZC3HC1 and TPR and analysed in parallel, using identical microscope settings, with staining for TPR also shown colour-graded to display differences in pixel intensities via a colour LUT. Arrowheads mark the nuclei of some MCL cells, i.e. those expressing the tag-free version of ZC3HC1, while arrows point at some of the progeny cells’ nuclei, i.e. those with the sfGFP-tagged ZC3HC1. Note first, in 5A1, that the amounts of sfGFP^L9mIAA7^-tagged ZC3HC1 appended to the NEs did not notably differ from the amounts in the neighbouring tag-free cells. Further note, in 5A2, that auxin-treatment had resulted in the elimination of essentially all visible NE-associated GFP and ZC3HC1 immunostaining in those cells that had been expressing the tagged version of ZC3HC1, while the MCL cells’ untagged ZC3HC1 polypeptides had remained unaffected. Moreover, note, in particular, that the elimination of the tagged ZC3HC1 had been accompanied by a conspicuous reduction in the immunolabelling intensity for TPR at the NE and a significant amount of TPR then distributed throughout most of the nuclear interior instead. Bar, 10 µm. **(B)** Quantification of signal yields for immunolabelled TPR at the NEs of the same mixed population of HCT116 MCL cells expressing the tag-less and sfGFP^L9mIAA7^-tagged ZC3HC1, after the addition of auxin. Randomly chosen NE segments for quantifications via ImageJ were essentially from all labelled cells in equatorial view within several randomly chosen images from the same specimen that also provided the micrograph for 5A2. Box plots display the relative signal intensity values, with the arithmetic means marked by x, with the ones for the cells not treated with auxin set to 100%, and with the SDs provided. Note that the mean TPR signal yield for the KO cells’ ZC3HC1-free NEs was only about half the WT cells’ corresponding value. **(C)** IB of cell extracts obtained from the sfGFP^L9mIAA7^-ZC3HC1 HCT116 cells after having treated these with DMSO or with auxin for 90 min. Applying conditions maintaining NB integrity, cells had been extracted with TX-100 like for Figure 1, resulting in fractions of soluble (S) and non-soluble proteins (P), with equal portions of each fraction’s whole amount then loaded for IB. Incubations with *Hs*TPR and *Hs*ZC3HC1 antibodies, and *Hs*NUP153 antibodies for comparison, were on different parts of the Ponceau S-stained membrane shown here and on another one with identical loadings. A cross-reaction unrelated to ZC3HC1 is marked by an asterisk. Note that ZC3HC1 had been largely degraded after the 90-minutes treatment with auxin, accompanied by a minor increase in the TPR amount detectable within the soluble cell fraction, while the presence of NUP153 within the pellet fraction had remained unaffected.

In its overall microscopic appearance, the redistribution of notable amounts of TPR from the NE into the nucleoplasm, which had come very rapidly into being, appeared essentially indistinguishable from what we had already seen in ZC3HC1 KO cells and WT cells after several days of ZC3HC1 RNAi. However, in at least one aspect, the current result differed from the two latter cases in which most soluble TPR likely had never been NB-associated and could readily be extracted by our standard cell fractionation protocols using detergent-containing NB-s buffers (our unpublished data, but see also Fig 1D and Supplemental Fig S1F): When applying the same fractionation conditions, the initially NB-associated and then, after degron-mediated ZC3HC1 elimination, detached TPR pool 2 was hardly extractable as soluble polypeptides. We regarded it possible that such renitent extractability might have to do with the detached TPR-containing ensembles perhaps not wholly disassembled and to some extent being still somewhat blocky instead. Alternatively, we could also imagine it might have to do with specific properties that these TPR molecules had acquired as NB-associated polypeptides (our unpublished data), with only the NB-appended molecules modified in a manner that allows for specifically interacting, then at the nuclear periphery only, with other nuclear binding partners. While this would prevent any newly synthesised and still soluble, not yet NB-associated TPR from prematurely engaging in such interactions, the modified TPR polypeptides, once detached from the NBs, might then bind to their partners also deeper in the nuclear interior, at sites where such interactions were actually meant not to happen.

Whichever of these assumptions might hold though: having tested, i.a., (i) different NB-s buffers, (ii) several variations of the standard fractionation conditions that did not noteworthily affect NB integrity in the control cells, and (iii) prolonged incubations with auxin, the thereby extractable amounts of the NB-detached TPR polypeptides had always been similarly moderate only (Fig 5C; and our unpublished data).

The main finding of these degron experiments, though, namely the essentially instantaneous detachment of large amounts of TPR from the NE concomitant to the loss of ZC3HC1, could now no longer be explained by several other imaginable scenarios that RNAi and gene disruption experiments had not been able to exclude (e.g. Gunkel *et al*, 2021, and as discussed further below). Instead, the current finding firmly argued for some direct role of ZC3HC1 in keeping subpopulations of TPR appended to the NB also *in vivo*, with ZC3HC1 acting thereby as a structural element that would either enable and stabilise some distinct direct interactions between TPR polypeptides or even function as a linker between them.

Such ZC3HC1-dependent interconnecting of TPR polypeptides did not even need to occur at the NB proper, as revealed by an experiment combining NUP153 RNAi and degron-mediated ZC3HC1 degradation (Fig 6). Knockdown of NUP153, a nucleoporin anchored to the NPC’s NR and with some role in the NPC appendage of the TPR pool 1 polypeptides, was already known to cause local subcellular accumulations of TPR in either cytoplasmic or nuclear foci, which formerly had been referred to as aggregates (Hase & Cordes, 2003). With such foci now also found positive for ZC3HC1 (Fig 6, and our unpublished data), and with such TPR-containing foci never seen upon KD or KO of ZC3HC1 (e.g. Gunkel *et al*, 2021; our unpublished data), we wondered whether ZC3HC1 might even play a direct role in keeping TPR agglomerated in such foci in the absence of NUP153. To address this question, we performed NUP153 RNAi in the ZC3HC1-sfGFP^L9mIAA7^ cell line. In the resulting NUP153-deficient cells, and in the absence of auxin, this again came along with TPR and ZC3HC1 mostly no longer NPC-appended but co-localising in multiple foci instead, with most of these here located in the cytoplasmic compartment (Fig 6). Strikingly then, upon treatment with auxin, apparently resulting in the degradation of most of the initially foci-positioned ZC3HC1, the foci too were essentially no longer visible. Their disappearance, in turn, was accompanied by the apparent release of the initially there-located TPR into solution, the nuclear import of the then soluble TPR polypeptides, and their distribution throughout the nucleoplasm in the absence of NUP153 and ZC3HC1 (Fig 6).

**Figure 6.**
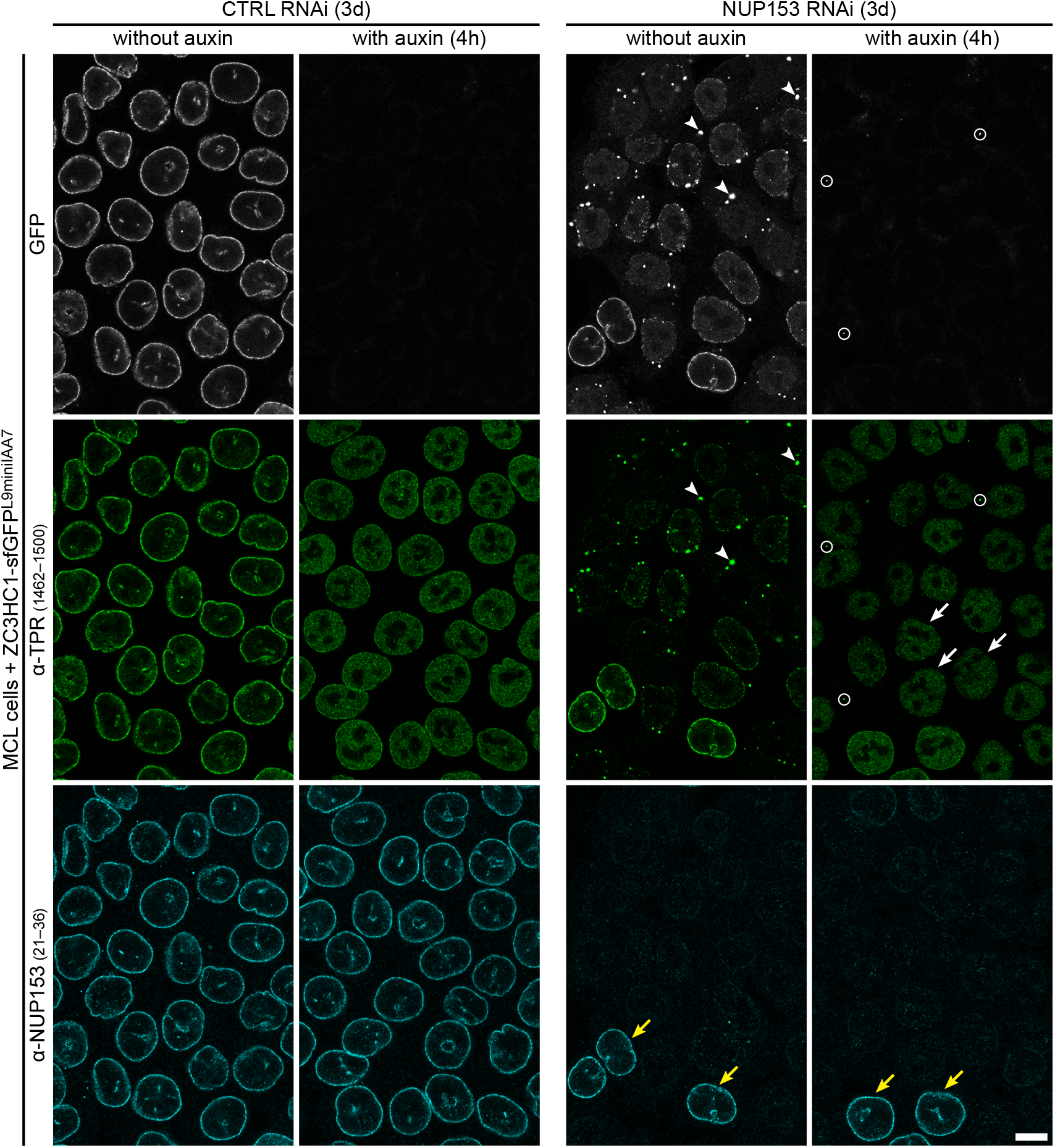
Ensembles of degron-tagged ZC3HC1 and TPR, remote from the NB due to NUP153 deficiency, are rapidly disassembled upon auxin-induced ZC3HC1 degradation, which results in TPR polypeptides unleashed again. IFM of sfGFP^L9mIAA7^-ZC3HC1 HCT116 cells treated with non-target control (CTRL) or NUP153 siRNAs and, at day 3 post-transfection, with DMSO or with auxin for 4 h, with all specimens then analysed in parallel, using identical microscope settings. Upon NUP153 RNAi, only traces of NUP153-staining were seen at most of the cells’ NEs; the usually bright NE-staining for NUP153 was only visible in cells that had remained non-transfected (some marked by yellow arrows), here shown as a reference. In the NUP153-deficient cells, NE staining for TPR and ZC3HC1 was conspicuously reduced, with the latter two proteins then co-localising, in the absence of auxin, in numerous, here primarily cytoplasmic foci instead (some marked by arrowheads). Note that ZC3HC1 and such foci were hardly detectable anymore (some small remnants encircled) after auxin-treatment, with TPR then appearing distributed throughout the NUP153- and ZC3HC1-deficient cells’ nucleoplasm instead (some marked by white arrows). Bar, 10 µm.

Altogether, these experiments had proven that ZC3HC1 is a structural element that allows for the interconnection of large amounts of TPR and is essential for keeping such TPR ensembles place-bound, irrespective of whether they are located at their natural sites at the NB or even at artificial ones elsewhere within the cell.

## Discussion

### ZC3HC1 as a novel, second structural element of the nuclear basket

In the current study, we have demonstrated that protein ZC3HC1 is a bona fide structural element of the NB. Such finding was the outcome of experiments by which we have dissected the process leading to the NB residency of the ZC3HC1-dependent TPR polypeptides, namely by systematically scrutinising whether ZC3HC1 is involved (i) in recruiting such TPR amounts to the NB, (ii) in appending them to TPR polypeptides already present at the NPC, and (iii) in then keeping the additional ones stably NB-associated.

The combination of different experimental approaches that had allowed for addressing the temporal order of the interactions underlying the reciprocal dependence of ZC3HC1 with TPR included assembly experiments, both *in vivo* and *in vitro,* by which these interactions could be studied in a stepwise manner, and the inducible, rapid and complete degron-mediated degradation of ZC3HC1, to assess its contribution for maintaining the TPR subpopulations’ residency at the NB. All these approaches had in common that they had allowed for a close spatiotemporal correlation between the addition or removal of ZC3HC1 and the resultant impact on the ZC3HC1-dependent pool of NB-associated TPR.

While former findings based on RNAi-mediated ZC3HC1 KD and *ZC3HC1* gene disruption had already revealed that the presence of ZC3HC1 in cultured cells is required for subpopulations of TPR to occur appended at the NB (Gunkel *et al*, 2021), it had remained elusive how ZC3HC1 would contribute to such a positioning of TPR pool 2 polypeptides. Since the formerly observed ZC3HC1 deficiency phenotypes had manifested themselves only time-delayed, with the cells first having run through mitosis and thus through the processes of NB disassembly and reassembly, it had not been possible to unequivocally attribute ZC3HC1 a role in either the recruitment and appendage or the stable positioning of such additional TPR amounts at the NB. Such uncertainty and caution in data interpretation had not been unfounded since we had also noted in our additional lines of research that other proteins appeared only transiently involved in some stages of the NB assembly process (our unpublished data). While some of these interactions might be required for adopting specific TPR conformations and NB arrangements as a prerequisite for further assembly steps, none of these other proteins has been classified as a structural component of the NB so far, unlike ZC3HC1.

However, the reason for exercising even more caution was the example of the multifaceted relationship between TPR and the NPC protein NUP153. This latter protein was back then the only other of TPR’s so far published direct binding partners that had been shown to play a role in the NB assembly process, either directly or indirectly involved in recruiting TPR pool 1 polypeptides and in attaching these to those NPCs that are assembled either after mitosis or in interphase (Walther *et al*, 2001; Hase & Cordes, 2003; Mendjan *et al*, 2006; Sabri *et al*, 2007; Mackay *et al*, 2010; Umlauf *et al*, 2013; Vollmer *et al*, 2015; Larrieu *et al*, 2018; Supplemental Information 4). On the other hand, though, NUP153 had turned out being seemingly no longer required by TPR once the latter had been anchored to the NPC’s NR (Lussi *et al*, 2010; Duheron *et al*, 2014; our unpublished data), which eventually also appeared in line with NUP153 and TPR having different anchor points at the NR (Supplemental Information 4). However, since NUP153 RNAi phenotypes relating to TPR-NUP153 interactions at different points of the cell cycle had seemingly also been varying to some extent, depending on cell types, on the RNAi experiment’s duration, and on the time points of inspection, the various cause-effect relationships observed upon such NUP153 RNAi had to be interpreted too with some caution until recently. Eventually, clarification only was provided by a groundbreaking study (Aksenova *et al*, 2019, 2020), in which induced rapid degradation of degron-tagged NUP153, thus providing an immediate phenotype, had allowed proving that NUP153, while required for NB formation and processes leading to NPC-appendage of TPR, was indeed not essential for keeping the NPC-anchored TPR polypeptides positioned there.

Hence, such dispensability of NUP153 for keeping bound TPR polypeptides in place, in addition to it not being an NB protein itself and not depending on TPR for its own binding to the NPC, clearly distinguished NUP153 from ZC3HC1. In point of fact, the detachment of essentially all TPR pool 2 polypeptides from the NE upon the degron-mediated rapid elimination of ZC3HC1 manifested the latter, unlike NUP153, as a structural component of the NB.

We regard this latter conclusion as not in conflict with the current study’s finding that the degron-mediated ZC3HC1 degradation led to somewhat more TPR detached from the NE than had been noted as absent upon ZC3HC1 RNAi or in ZC3HC1 KO cells (Gunkel *et al*, 2021; our unpublished data). Instead, we consider it conceivable that the rapid elimination of ZC3HC1 could have even affected the NPC association of some of the TPR 1 polypeptides that are usually appended to the NPC independently of ZC3HC1. Even without an experimental data-based explanation currently available for these moderate differences, we can imagine, for example, that a proteasome degrading an AID-tagged ZC3HC1 might also consume the one or other TPR pool 1 polypeptide if some of the degron-tagged ZC3HC1 polypeptides might be very tightly bound to them. We regard such a scenario as not entirely unreasonable, given former findings that a degron-tagged nanobody can confer degradation of its non-tagged binding partner if tightly bound to it (e.g., Daniel *et al*, 2018). On the other hand, however, such co-degradation of non-tagged binding partners does not appear to happen for again other pairs of non-tagged and degron-tagged interaction partners (e.g., Beer *et al*, 2019), and we can, alternatively, also imagine that the TPR pool 1 polypeptides and the residual NB scaffold, once rapidly stripped of all ZC3HC1 and TPR pool 2 polypeptides, might be less durable and more prone to disintegration than NBs built from the ground up in the absence of ZC3HC1.

However, the results of the current study’s assembly experiments also indicated that ZC3HC1 is not merely a structural element required for keeping TPR 2 polypeptides positioned at the proliferating cells’ NBs but also needed for the actual recruitment and appendage of such TPR polypeptides. On first thought, this might appear unavoidable and thus self-evident for a structural protein required for maintaining the integrity of higher-order protein ensembles because one could imagine its presence, of course, also a prerequisite for the other proteins’ successful recruitment and appendage. However, even though a distinction between recruitment, appendage and maintenance might appear superfluous, and challenging to attest for a structural protein anyhow, we nonetheless regard it appropriate to carefully distinguish between such different contributions in the context of the NB’s assembly and maintenance of its higher-order arrangements. To us, the latter appear more complex and dynamically variable than as one might have formerly anticipated.

In fact, we can easily imagine, for example, the existence of additional proteins that are not essential for the NB assembly process and the initial appendage of additional amounts of TPR to the NB but that, in addition to ZC3HC1, can nonetheless play a role too in maintaining the additional TPR polypeptides’ relative positions, once they have been appended to the NBs. Among the candidates conceivable are, for example, various nucleic acid-binding proteins, including DNA-binding proteins, that one can find detached from nuclease-treated NEs together with TPR, ZC3HC1 and other known NB proteins upon the disintegration of NBs in different cell types (our unpublished data). We can now conceive a scenario in which the one or other protein capable of interacting with both TPR and nucleic acids might qualify for additionally stabilising the NB via lateral interactions with neighbouring chromatin, possibly even in a manner varying between different cell types. Again, however, we currently do not regard such other proteins, which too might be involved in maintaining the perinuclear positions of the NB-associated TPR subpopulations, as qualifying themselves as additional structural elements of the NB; again, unlike ZC3HC1.

### A methodological toolbox for NB research

The type of *in vivo* and *in vitro* assembly experiments presented in the current study might now turn out helpful in answering some of the still open questions regarding the contributions of different proteins, including NB components and other proteins like NUP153, to specific steps in the NB assembly process. For example, in the case of NUP153, one could imagine creating NEs free of NUP153 and NBs, via degron-mediated co-elimination of NUP153 and TPR, which could then be used for the stepwise re-addition of recombinant versions of NUP153 and NB components. Furthermore, we regard approaches of such kind, in which NE scaffolds are used as assembly platforms for ectopically expressed WT proteins and mutant versions thereof, as also suitable for providing insight into how the stepwise interactions take place between further NPC and NB components and other NB-interacting proteins. While it will not be possible to generate CRISPR/Cas9-edited KO cell lines for every NPC, NB, and NB-associated protein of interest, simply because some of them are essential, NEs deficient of specific target proteins can also be acquired by their RNAi-mediated KD or their degron-mediated degradation, once the corresponding CRISPR/Cas9-edited cell lines are available.

We can also imagine using the NE scaffolds, then with complete NBs, as platforms suited for screening and identifying proteins that only interact with the NB as a structural entity. Since circumstantial evidence suggests (our unpublished data) that some transiently NB-interacting proteins can only interact with some of the NB’s components when the latter are part of the NB’s higher-order structural arrangements, such interactions would not be readily identifiable by conventional affinity chromatographic approaches or other means based on pairwise interactions only, like, e.g., yeast two-hybrid methodology.

Furthermore, the resulting cell lines can be versatilely used after having tagged the NB research-related genes of interest with the here presented sfGFP^L9mIAA7^ cassette. Apart from the induced rapid degradation of the tagged proteins, to study resulting phenotypes, such lines will also allow for the standardised use of GFP-specific single-domain antibodies (sdAbs), not only for the immunoprecipitation of the tagged proteins but also for their visualisation at higher resolution by super-resolution microscopy.

Beyond that, we propose using such sfGFP^L9mIAA7^-expressing cell lines, particularly when corresponding cell lines of different germ layers and tissue origins are available to be used in parallel, to gain further insight into the standard and cell-type-specific repertoire of proteins that interact with the NB and its appended structures. Such distinction will be achievable, i.a., by mass spectrometry, when inspecting the protein composition of the lamina-NPC-NB (LNN)-enriched materials isolated from such cells before and after having induced the degradation of different NPC and NB proteins, as it has here been exemplified by the degron-mediated degradation of TPR and the resulting loss of NB proteins like ZC3HC1 from such LNN materials. Systematically investigating this not only in one but several cell lines, including correspondingly CRISPR/Cas9-edited stem cells and thereby applying not only one but several different LNN isolation procedures in parallel, might eventually provide an inventory of those proteins that only transiently interact with the NB while enriched at these sites in a steady state nonetheless. Furthermore, systematically comparing populations of quiescent and cell-cycle-synchronised cells should also provide insight into possibly cell cycle-phase-specific interactions between the NB components and other proteins. Finally, such an approach should also contribute to further distinguishing between universal and cell-type-specific NB-resident proteins. Altogether, such subtractive proteomics based on degron-mediated target protein elimination would complement and exceed a related approach of comparing NEs with and without NBs, in which the latter had been detached by physicochemical means from the NEs of *Xenopus* oocytes, resulting in the initial identification of ZC3HC1 as an NB-resident protein (Gunkel *et al*, 2021).

### The open question of how and how many of the non-essential ZC3HC1 polypeptides keep how many TPR polypeptides in which type of arrangements where appended to the NB, and why so

The current study does not yet permit to unambiguously tell how ZC3HC1 exerts its function as a structural element required for keeping the TPR pool 2 polypeptides bound at the NB, as the data here do not yet allow for unambiguously distinguishing between different scenarios. On the one hand, ZC3HC1 could be a protein whose interaction with TPR pool 1 polypeptides would force the latter into a specific conformation, only thereby forming a platform that would allow for direct interactions between TPR pool 1 and pool 2 polypeptides. Alternatively, one could also imagine ZC3HC1 as a protein forming a socket or flange that too would allow for some direct interaction between the TPR pool 1 and pool 2 polypeptides, with the latter mounted on the former, or as a protein stabilising in yet another manner some direct interactions between the TPR polypeptides. On the other hand, though, it is, of course, also tempting to imagine ZC3HC1 as an interconnector positioned between the different populations of TPR polypeptides, with such a setup then not necessarily requiring any direct contact between the TPR pools 1 and 2. However, which of these scenarios, not all of which would be mutually exclusive, will eventually hold, will have to be the topic of studies still to come.

Such studies will also need to delineate those parts of ZC3HC1 and TPR that enable them to engage in such a relationship. In addition, they will have to provide the total copy numbers for those ZC3HC1 and TPR polypeptides that engage in such interactions at the NB, as only such numbers will eventually allow sketching which type of arrangements and constellations among these proteins at the NB will be numerically possible at all.

Furthermore, since current evidence already points at different subpopulations of TPR that occur NB-associated, their copy numbers, too, need to be known. While the current study has been referring to at least two major TPR subpopulations, called T1 and T2 for simplification, data from further experiments, in which we have systematically disassembled the NB stepwise in more detail, have hinted at even further subpopulations within those of T1 and T2. For example, in the case of the T1 pool, this could mean that it might actually be composed of two equally large subpopulations, T1a and T1b (Gunkel *et al*, 2021; our unpublished data). Similarly, several pieces of circumstantial evidence, both in the recent study (Gunkel *et al*, 2021) and the current one (Supplemental Fig S6), have also been pointing at the existence of ZC3HC1 subpopulations that appear to engage in different types of interactions with TPR and the NB, with each ZC3HC1 subpopulation perhaps having a distinct task while nonetheless acting cooperatively, or instead sequentially, in the TPR assembly processes. Further support for such notions also comes from the data of systematic cell fractionation and NB disassembly experiments (Gunkel *et al*, 2021; our unpublished data), which, too, point at NB-associated ZC3HC1 subpopulations with differing dissociation characteristics, and at some point, the copy numbers for each of these different subpopulations will need to be unriddled as well.

Another question among the central ones regarding the NB’s architecture in the proliferating human cell is how the ZC3HC1-recruited TPR populations are arranged and positioned relative to those TPR polypeptides that are anchored to the NPC independently of ZC3HC1. In the one scenario, one can imagine both pools representing components of the same structure commonly regarded as the prototypic NB, with both the T1 and T2 polypeptides perhaps even longitudinally aligned. On the other hand, it is very tempting to envision the T2 polypeptides as appended to the NB in such a manner that would allow them to project away from the NB and reach out further into the nuclear interior, similar to arrangements that have been described for NBs in *Xenopus* oocytes (e.g., Gunkel *et al*, 2021). However, to address this and further questions regarding the NB’s assembly blueprint and final architecture, one will need to dissect the NB assembly process in further molecular detail by also using yet further mutant versions of ZC3HC1 and TPR. Moreover, one will now also need to microscopically visualise the arrangement of the different TPR subpopulations, relative to each other and ZC3HC1, and use, therefore, the emerging novel methodologies in super-resolution microscopy that will allow for resolving these proteins’ positions relative to the NPCs in different human cell lines.

The question, though, that we still regard perhaps the most pressing one is why evolution has decided to stick with ZC3HC1 in a wide range of species, even though the protein appears to be dispensable for all those species in which the corresponding homolog hitherto has been eliminated experimentally (Gunkel *et al*, 2021, and references therein). For example, such dispensability holds for also *Saccharomyces cerevisiae* Pml39p, a protein that binds to the yeast’s TPR homologs Mlp1p and Mlp2p (Palancade *et al*, 2005) and that we found to be the budding yeast’s sole homolog of ZC3HC1 (Gunkel *et al*, 2021; our unpublished data). Moreover, even evolution itself has sometimes allowed for discarding an initially present ZC3HC1 homolog again, with this having happened and seemingly still happening within some subphyla, orders, or even families of organisms. Such evolutionary fate of ZC3HC1 can be reconstructed, for example, in insects, in which almost all orders, including Diptera and thus also *Drosophila*, lack an apparent ZC3HC1 homolog, while unambiguously still present in some species of other insect orders and while functionally likely intact ZC3HC1 homologs are omnipresent in all other subphyla of the Arthropoda (Gunkel *et al*, 2021; our unpublished data).

For those organisms in which ZC3HC1 acts as a structural NB element that would allow for appending additional TPR polypeptides, this raises the question of what the latter might be good for and, conversely, why the ZC3HC1-deficient organisms have adapted to getting along without such additional amounts of TPR, provided that such species have not come up with some protein functionally analogous to ZC3HC1 or some other mechanism of appending more TPR to their NBs.

Recently, several conceivable tasks that one could assign to the ZC3HC1-dependent populations of TPR polypeptides have already been outlined (Gunkel *et al*, 2021). For example, apart from potential contributions to the NB’s overall structural stability and involvement in various other tasks, one scenario regarded ZC3HC1 as a protein that would allow for increasing the number of TPR polypeptides for the sake of these then acting as either transient or more lasting binding sites at the NB for yet other molecules, with the copy numbers of such additional TPR polypeptides perhaps even modifiable upon demand (Gunkel *et al*, 2021). Among the TPR-interacting candidate molecules in such a scenario, one could then easily imagine proteins involved in transcription regulation, replication, and perinuclear chromatin organisation. Such type of interactions would be in keeping with the NB and TPR in metazoans and *Arabidopsis* (e.g., Mendjan *et al*, 2006; Skaggs *et al*, 2007; Vaquerizas *et al*, 2010; Zhang *et al*, 2016; Myers *et al*, 2016; Pérez-Garrastachu *et al*, 2017; Yang *et al*, 2017; Raich *et al*, 2018; Boumendil *et al*, 2019; Krull *et al*, 2010; Su *et al*, 2018; Cohn *et al*, 2020; Uhlířová *et al*, 2021; Kosar *et al*, 2021; Wu *et al*, 2021; Aleman *et al*, 2021; our unpublished data), and the Mlp1p/Mlp2p and Mlp1p/Mlp2p-associated proteins in budding yeast, having been found engaging in interphase in different types of interactions with NB-neighbouring chromatin and a subset of genes (e.g., Casolari *et al*, 2004, 2005; Dieppois *et al*, 2006; Cabal *et al*, 2006; Schmid *et al*, 2006; Luthra *et al*, 2007; Brickner *et al*, 2007; Tan-Wong *et al*, 2009; Bermejo *et al*, 2011; Texari *et al*, 2013; García-Benítez *et al*, 2017), with only some of the latter here listed.

Of course, though, this scenario too, like all other ones trying to explain the add-on of additional TPR polypeptides, instantaneously leads us back to the question of why ZC3HC1-deficient organisms like *Drosophila*, in which interactions between TPR and the autosomes and male X-chromosome occur nonetheless, with *Drosophila*’s TPR homolog Mtor also directly involved in transcription regulation (e.g., Mendjan *et al*, 2006; Vaquerizas *et al*, 2010; Raich *et al*, 2018; Aleman *et al*, 2021), would not require more TPR appended to their NBs, and why they do well or perhaps even better without them. The answers to such fundamental questions might eventually provide us with an explanation for why evolution has decided that we vertebrates should rather hang on to ZC3HC1 as a second building element of the NB.

## Materials and Methods

### Antibodies

All specifications regarding antibodies used in this study are provided in Supplemental Table S1. Among these are novel *Hs*TPR antibodies, all raised in rabbits, that were used for immunoaffinity depletion of TPR from human cell extracts (see below). One of these TPR antibodies was obtained after immunisation with a chemically synthesised peptide, corresponding to *Hs*TPR aa 2147-2163, that was phosphorylated at S2155. The others were obtained after immunisation with recombinant fragments of *Hs*TPR followed by isolating subpopulations of peptide-specific antibodies from the resulting sera, using series of overlapping peptides for affinity chromatography, as recently described for collections of *Xl*TPR peptide antibodies (Gunkel *et al*, 2021).

### Cell culture, auxin treatments, RNAi, synchronisation, and transfections

All cell lines used in this study, including those generated by CRISPR/Cas9-editing, and their growth conditions, including types of media used, are listed in Supplemental Table S2. Cell cycle synchronisation was performed as described (Gunkel *et al*, 2021). To initiate auxin-induced degradation, we first added the auxin indole-3-acetic acid (I5148, Sigma-Aldrich) from a 500 mM stock solution in DMSO, freshly prepared or frozen for no longer than one week and then thawed, to the cell-free cell culture medium at a concentration of 0.5 mM, followed by incubation at 37°C and 5% CO2 for 30–60 min. This pre-warmed, auxin-supplemented medium was then used to replace the medium of the cells. Concomitantly, a pre-warmed medium supplemented with auxin-free DMSO was applied to cells representing the uninduced control population. Transfections of HCT116 cells with small interfering RNAs (siRNAs; Silencer Select negative controls #1 and 2 (#4390843 and #4390846) and Silencer Select siRNAs targeting *Hs*NUP153 (s19374 [CGAAAAUCUCUCUACCGAU] and s19376 [CAGUCUAA ACUACGAAAUA]), Ambion Inc., Austin, TX, USA) were performed as described (Gunkel *et al*, 2021). For the transfections of HeLa with mammalian expression vectors presented in Figure 1, we used PolyJet (SignaGen Laboratories, Frederick, MD, USA) as the transfection reagent, following the manufacturer’s instructions. For transfections of HeLa and HCT116 cells in the course of the gene-editing procedures, see further below. For large scale transfections of the adherent HEK293T cells for the ectopic expression of proteins to be used for the NB *in vitro* assembly experiments, we followed a transfection protocol (Reed *et al*, 2006) using linear 25 kDa polyethylenimine (PEI; Polysciences Europe GmbH, Hirschberg, Germany). We adjusted this protocol for 10 cm culture dishes with about 50-70% cell density, with these and further modifications already described (Gunkel *et al*, 2021), yet with the current study’s cells harvested 16–21 h after the initial transfection.

### Immunofluorescence microscopy and live-cell imaging of cultured cells

For standard IFM, cells were fixed for 30 min with 2.4% of freshly prepared and methanol-free formaldehyde (FA) in PBS, followed by quenching with 50 mM NH_4_Cl in PBS, and permeabilisation with 0.25% Triton X-100 (TX-100) in PBS for 5 min. In cases in which detergent-permeabilisation was carried out before FA-fixation (Fig 1D), cells were treated with 0.25% TX-100 in a temporarily NB-stabilising (tNB-s) buffer for 3 min, the latter here composed of PBS to which MgCl_2_ was freshly added to a final concentration of 10 mM, followed by fixation in tNB-s buffer, and quenching with NH_4_Cl in PBS, yet omitting subsequent treatments with detergents. Blocking and antibody incubations were performed with 1% BSA in PBS, with Hoechst 33342 (1 µg/ml) for DNA staining added during the incubation with fluorophore-coupled secondary antibodies. The immunolabelled specimens were mounted in SlowFade Gold or Diamond Antifade Mountant (Invitrogen, Carlsbad, CA, USA), the latter used for FA-fixed GFP-tagged specimens and the subsequent inspection of GFP signals. For live-cell imaging, cells were grown on 4-well or 8-well ibidi glass-bottom (#1.5H) µ-slides (ibidi, Martinsried, Germany), using either Leibovitz’s L-15 medium (Sigma-Aldrich) or the live-cell imaging medium FluoroBrite DMEM (GIBCO), and an improved version of SiR-Hoechst (Bucevičius *et al*, 2019) kindly provided by Gražvydas Lukinavičius for live-cell DNA staining. All cells, i.e. the immunolabelled ones and the intact live cells, were inspected with a Leica TCS SP5 or SP8 confocal laser-scanning microscope (Leica Microsystems, Wetzlar, Germany) equipped with a 63x oil immersion objective (Leica Microsystems).

### Cell fractionation and immunoblotting

Fractionation of confluent, not synchronised populations of HCT116 cells, to obtain, for IB, a fraction of soluble proteins released upon extraction with TX-100 and a non-soluble, LNN-enriched fraction, was in principle as described (Gunkel *et al*, 2021), applying conditions, with only minor modifications, that allow for maintaining the interactions of TPR and ZC3HC1 with the NBs and the LNN-enriched material. In brief, washed cells of lines HCT116 MCL and sfGFP^L9mIAA7^-TPR, that had been treated with auxin or not, were sedimented by 800× *g* centrifugation for 3 min, followed by resuspending the cells in 20–22°C-warm NB-s solution containing 41.5 mM KCl, 8.5 mM NaCl, 5 mM MgCl_2_, 2.5 mM EGTA, 2 mM DTT, 10% sucrose, 20 mM HEPES, pH 7.5, 0.25% TX-100, and cOmplete EDTA-free protease inhibitor cocktail (Roche), followed by incubation for 4 min and centrifugation at 20,000× *g* and RT for 3 min. Cells of line HCT116 ZC3HC1-sfGFP^L9mIAA7^ were handled similarly, except for incubating the batches of ZC3HC1-sfGFP^L9mIAA7^ cells, after having been treated in parallel with and without auxin, in the NB-s buffer containing TX-100 for 15 min. The fractionations of HeLa and HEK293T cells for the assembly assays presented in Figures 2 and 3 were performed as explained further below, and SDS–PAGE and IB as described recently (Gunkel *et al*, 2021).

### *In vitro* assembly experiments with ZC3HC1-deficient NEs and fluorescent protein-tagged ZC3HC1 and TPR polypeptides

For preparing TPR-free cell extracts with either the mCherry-tagged WT or C429S mutant version of ZC3HC1, up to about 1.4×10^8^ adherent HEK293T cells had been transiently transfected with corresponding expression vectors (see Supplemental Table S3) and harvested 16–21 h later. Cells were sedimented by a 3 min centrifugation at 1,000× *g* and resuspended in an NB-s buffer, here called assembly buffer, composed of 88 mM KCl, 22 mM NaCl, 10 mM MgCl_2_, 10 mM potassium sodium phosphate (4:1 molar ratio of K_2_HPO_4_/KH_2_PO_4_: Na_2_HPO_4_/ NaH_2_PO_4_), pH 6.8. The cells were then recentrifuged at 1,500× *g* and RT for 2 min, resuspended, at a volume ratio of about 1:4 between pellet volume and solution, in assembly buffer supplemented with cOmplete Mini EDTA-free protease inhibitor cocktail (Roche) and 0.05% digitonin, for permeabilisation at RT for 3 min. They were subsequently centrifuged at 20,000× *g* and RT for 2 min, followed by recentrifuging the resulting supernatants at 200,000× *g* at 4°C for 10 min. The then obtained solutions were rotated at RT for 20 min together with Protein A-coupled magnetic Dynabeads (Invitrogen, Carlsbad, CA, USA) that had been loaded beforehand with a collection of five different rabbit antibodies targeting different parts of *Hs*TPR (see Supplemental Table S1). Loading of these magnetic beads with TPR antibodies had been performed in the assembly buffer containing 0.02% Tween-20 at RT for 15 min, followed by several washes in the same buffer. For the immunodepletion of such soluble HEK293T extracts, then generally corresponding to still about 5–9×10^7^ cells, a total of up to about 18–21 μg of anti-TPR IgGs were used, with each of the five different TPR antibodies contributing different amounts, ranging between 2.9 to 5.8 μg. The antibodies’ corresponding total copy numbers coupled to the magnetic beads had been calculated to exceed the maximally expectable total numbers of soluble TPR dimers within the extracts at least 200-fold. Following the magnetic removal of the beads, the solutions were directly used for the interaction experiments; but later on, they were always controlled by IB to have been devoid of TPR. On the other hand, the ZC3HC1-free cell extracts containing sfGFP-tagged TPR were obtained from one of the sfGFP-TPR-expressing HeLa ZC3HC1 KO cell lines in which substantial amounts of TPR normally occurring NE-associated in WT cells are distributed throughout the nuclear interior in a soluble form instead. To this end, about 1×10^8^ cells from sub-confluent populations, i.e. from such, in which cell sizes were still notably larger than in highly confluent populations of HeLa, were first washed in assembly buffer. They were then sedimented by centrifugation at 1,000× *g* for 3 min, resuspended in assembly buffer, and recentrifuged for 2 min at 1,000× *g*. Next, they were resuspended in assembly buffer supplemented with cOmplete Mini EDTA-free protease inhibitors, at a volume ratio of about 1.25:1 between buffer and pellet volume, the latter though still containing buffer from the preceding washing step, so that the actual ratio between buffer and total cell volume came rather close to 1.8:1. All steps were performed at RT. About 5×10^7^ cells as 325 µl suspensions in 1.5 ml safe-lock tubes (Eppendorf AG, Hamburg, Germany) were ruptured by brief pulsed sonication at RT using the ultrasonic homogeniser SONOPULS mini20, equipped with an MS 2.0 sonotrode (Bandelin Electronic GmbH, Berlin, Germany), applying conditions (sonotrode immersion depth of 1 mm, amplitude setting 50%, pulsating mode, a 15-sec total length of operation, with pulses of 0.5 sec separated by pauses of 1 sec), which had been heuristically determined by series of trial-and-error experiments to result in an adequate, i.e. a probably complete rupturing of the suspended HeLa cells’ nuclei without causing disruption of the NEs primarily into fragments no longer sedimentable at 20,000× *g*, while at the same time allowing for minimising energy input and keeping the mean rise in temperature within the suspension below two degrees. The cell suspensions were then centrifuged at RT and 20,000× *g* for 2 min, followed by re-centrifugation of the resulting supernatant at 4°C and 200,000× *g* for 10 min. Supernatants obtained in such a way, free of any detectable NE fragments but containing soluble sfGFP-TPR polypeptides, were used for the assembly assays straight away. Note that for time reasons, the preparation of such freshly prepared HEK293T and HeLa cell extracts for their direct use in the assembly experiments required two persons’ work efforts in parallel. For preparing the NEs as assembly platforms, HeLa ZC3HC1 KO cells were grown on 8-well ibidi µ-slides with ibiTreat bottom (ibidi), and populations of 50–70% cell density were then permeabilised with detergent just prior to the addition of the cell extracts. To this end, the cells were incubated for about 2 min to maximally 3 min in assembly buffer containing 0.25% TX-100 and Hoechst 33342 (1 µg/ml), accompanied by gentle rocking, followed by three washes with detergent-free assembly buffer, with the first wash step for 2 min still containing the DNA dye, then omitted from the next steps. All subsequent incubations with the mCherry-ZC3HC1- and the sfGFP-TPR-containing solutions, and all washes, using the detergent-free assembly buffer supplemented with cOmplete Mini EDTA-free protease inhibitors, were performed at RT. After having loaded the nuclei with the recombinant polypeptides, the specimens were inspected with a Leica TCS SP8 confocal laser-scanning microscope (Leica Microsystems, Wetzlar, Germany), with all images of mCherry and sfGFP fluorescence, respectively, then acquired with identical microscope settings and reduced laser power, as compared to IFM images, to minimise pronounced bleaching in the absence of anti-fade media. Signal brightness of all raw images was then enhanced electronically, which was done in the same manner for each series of corresponding images. To this end, all corresponding colour images of either mCherry or sfGFP were first assembled into one image, then converted to the 8-bit greyscale of 0–255. Each pixel value of this composite image was then multiplied by the same multiplication factor, using for this purpose the Multiply command in the Math Submenu of the ImageJ/FiJi software (version 2.0.0-rc-64/1.51t, National Institutes of Health, USA). With such constant multiplication of pixel intensities being the only image processing conducted for the *in vitro* assembly experiments, and with less than 0.05% of each image’s pixels having reached a value of 255, this procedure allowed the signal intensity relationships between the electronically brightness-enhanced images to remain essentially the same as between the corresponding raw images beyond their background zero values.

### Genomic editing in human cell lines

Deletion of ZC3HC1 gene segments in human cell lines, using CRISPR/Cas9 technology and following the Cas9 double-nickase approach with pairs of single guide RNAs (sgRNA) (Ran *et al*, 2013a), was as described in detail recently (Gunkel *et al*, 2021). This approach was also used for tagging the TPR and ZC3HC1 alleles with the ORF for sfGFP or variants thereof. Novel pairs of sgRNAs, with sequences complementary to the genomic regions at which the integration was to occur (for sequences, see Supplemental Table S4; also see Supplemental Figs S1, S2, S8, S9), were once again designed by using the CRISPR Design tool formerly provided online by the Zhang laboratory (Hsu *et al*, 2013). This again was followed by the cloning of these sgRNAs into the bicistronic Cas9n expression vector pSpCas9n(BB)-2A-Puro (PX462) V2.0 (Ran *et al*, 2013b) kindly provided by Feng Zhang (Addgene plasmid #62987; http://n2t.net/addgene:62987; see also Supplemental Table S3), with further details as described (Gunkel *et al*, 2021) but here using only one pair of sgRNAs and thus two sgRNA/Cas9n vectors, instead of two pairs, per integration site. Cultured cells were then transfected with the GFP-donor plasmid and the two target site-specific sgRNA/Cas9n vectors, using PolyJet (SignaGen Laboratories) for the transfection of HeLa cells, and FuGene HD (Promega Cooperation, Madison, WI, USA) for HCT116 cells, according to the manufacturer’s instructions. Three to four days after transfection, the sfGFP-expressing cells were enriched by sorting via flow cytometry, using a Bio-Rad S3e cell sorter (Bio-Rad Laboratories, Hercules CA, USA). In some cases, this step was repeated with the first enriched population of GFP-positive cells to sort for the brightest ones, often representing those in which the sfGFP ORF had been integrated into all of the target gene’s alleles. These enriched populations were then seeded into 10 cm culture dishes at a low cell density to allow for the growth of monoclonal colonies that could be manually picked and transferred into 15-well µ-slides (µ-slide angiogenesis ibiTreat bottom; ibidi) for further growth. The clones were screened by live-cell imaging, assessing sub-cellular GFP localisation and comparing the different clones’ NE-associated GFP brightness under identical microscope settings. The brightest NE-positive clones were selected for colony expansion, followed by further analysing them by comparative IFM and IB with the corresponding progenitor cell line. In addition, all monoclonal populations of the CRISPR/Cas9-edited KO and GFP-expressing cell strains were analysed by genomic PCR and DNA sequencing of subcloned PCR products (for primers, see Supplemental Table S5).

For generating the HCT116 MCL with constitutive *At*AFB2 expression, HCT116 WT cells were transfected with the sgRNA/Cas9 vector (kindly provided by Masato Kanemaki) together with the integration vectors for *At*AFB2-Myc and *At*AFB2-Myc-NLS (based on a vector kindly provided by Elina Ikonen) using FuGene HD transfection reagent (Promega). Puromycin (5 µg/mL) was added for the selection of transfected cells that were grown until monoclonal colonies could be manually picked. Clones were screened by IFM for the presence and subcellular distribution of the Myc-tagged *At*AFB2. The finally selected MCL was used for further integration of degron-tags in combination with sfGFP (see above).

## Acknowledgements

We acknowledge Gražvydas Lukinavičius, Elina Ikonen, Masato Kanemaki, and Feng Zhang for kindly providing research reagents and Dirk Görlich for sharing equipment and advice. In addition, we thank Thomas Güttler and Georg Krohne for their critical reading of the manuscript.

## Author contributions

Conceptualization, V.C.C. and P.G.; data curation, P.G. and V.C.C.; formal analysis, P.G. and V.C.C.; investigation, P.G. and V.C.C.; methodology, P.G. and V.C.C.; project administration, V.C.C. and P.G.; resources, V.C.C.; supervision, V.C.C.; validation, P.G. and V.C.C.; visualization, P.G.; writing—original draft preparation, V.C.C. and P.G.; writing—review and editing, V.C.C. and P.G. Both authors have read and agreed to the submitted version of the manuscript.

