## Supplemental Data for "ZC3HC1 is a structural element of the nuclear basket effecting interlinkage of TPR polypeptides"

Running title: Nuclear basket architecture and ZC3HC1

#### Supplemental Information, Figures, Tables and References

#### Supplemental Information

**Supplemental Information 1. Motivation for a TPR *in vivo* re-recruitment experiment.**

In the early course of our investigation (Gunkel *et al*, 2021), we had noted that ZC3HC1 deficiency, either by RNAi or gene disruption, had always come along with a very conspicuous reduction in the NE-associated amounts of TPR, and often with also a nucleoplasmic pool of soluble TPR polypeptides. The latter observation instantaneously raised the question of whether these soluble TPR polypeptides could be reused for being appended to the NE, provided that ZC3HC1 would become available again. Or, alternatively, whether a soluble pool of TPR, either detached from the NE or, more likely, never having been attached to it at all but having strayed through the nucleus instead, would irreversibly no longer be useable for any incorporation into the NB.

One could conceive various possibilities of how such a latter situation could come about. For example, in the case of the NB-detached TPR polypeptides, one could imagine that they might have acquired distinct features whilst they were part of the NB, with these though preventing the reattachment of such TPR polypeptides to the NB once they had been detached from it. Such features could include post-translational modifications required to perform certain tasks or adopt specific conformations at the NB.

In the other scenario again, in which the soluble TPR polypeptides would not yet have been part of an NB, one could imagine that only the newly synthesised TPR polypeptides would be adapted for a ZC3HC1-dependent NB-association process. In the course of such a process, theTPR polypeptides would need to pre-assemble with ZC3HC1 early on, perhaps otherwise irreversibly folded or again post-translationally modified in such a way that NB appendage would no longer be possible. And in yet another scenario, one could imagine that the NB-association of the ZC3HC1-dependent TPR polypeptides would even occur only during an NB formation step that required co-assembly with the ZC3HC1-independent TPR pool.

Thus, if the one type of scenario would hold, only TPR polypeptides newly synthesised together with ZC3HC1 could be appended to a residual NB composed of the ZC3HC1-independent TPR polypeptides only. And in the other, even less permissive scenario, the newly synthesised TPR polypeptides would be incorporated only into those NBs that are newly assembled from the ground up. In other words, if the soluble TPR polypeptides were indeed useless for ectopically expressed ZC3HC1 that was newly synthesised in a ZC3HC1 KO cell line, it would take a long time, then likely ranging from at least many hours to possibly even several cell cycles, before most NPCs would be equipped again with their normal amounts of TPR. Such notion had also taken available evidence into account that did not indicate any notably different TPR synthesis rates in the WT and the ZC3HC1 KO cells, even not upon the ectopic expression of ZC3HC1 (our unpublished data). Such finding had not come as a major surprise since HeLa was known to lack dosage compensation in gene expression (e.g., Landry *et al*, 2013).

In again other words, even if ectopic ZC3HC1 expression in a formerly ZC3HC1-deficient cell would rapidly lead to those ZC3HC1 amounts usually required for having the additional TPR amounts NB-appended, it would not provide much clarification as to whether and how ZC3HC1 might directly contribute to the TPR polypeptides' recruitment if the latter would first need to be newly synthesised in large amounts for some time.

However, in the alternative, quite different scenario, the already existing pool of soluble TPR in such a ZC3HC1 KO cell line would be reusable, provided that ZC3HC1 were made available again. In this case, only the amount of ZC3HC1 would be determining the rate of such NB re-association of TPR polypeptides. In fact, if it were possible to have a sizeable nucleoplasmic pool of soluble TPR polypeptides rapidly re-recruited and appended to the NE, concomitantly to the reappearance of ZC3HC1, and to hence rule out a substantial contribution by newly synthesised TPR polypeptides, one could rate such a finding as evidence for ZC3HC1 playing a direct role in the process that leads to the recruitment of a TPR subpopulation and its attachment to the NE.

**Supplemental Information 2. Assessment of the prerequisites for a TPR re-recruitment experiment *in vivo*.**

To investigate whether ZC3HC1 plays a direct role in recruiting TPR polypeptides to the NE *in vivo*, we performed the *in vivo* experiment presented in Figure 1. Priorly, however, we performed some estimating calculations and control experiments in order to assess whether the experimental setup could, in principle, deliver results of potential diagnostic value. For these estimations, we took the following assumptions and already available pieces of information into account, with some of them considering the amount of TPR to be re-recruited to the NEs (points 1–4). The others addressed the potential copy numbers of ZC3HC1 required for such a TPR re-recruitment process and whether such numbers could realistically be provided by the ectopic expression of ZC3HC1 in the ZC3HC1 KO cells within only a short time window (points 5 and 6).

(1) About half of the NPCs within a proliferating cultured cell are assembled towards the end of mitosis, by a process called post-mitotic NPC assembly (e.g., Doucet & Hetzer, 2010; Rothballer & Kutay, 2013; Otsuka & Ellenberg, 2018; Kutay *et al*, 2021). The NBs are then appended to these NPCs very early in G1, with nearly if not all of the post-mitotically assembled NPCs then regarded as equipped with an NB, at least in a tumour cell, like HeLa. From then on, additional NPCs and NBs are assembled by a process referred to as the interphase assembly of NPCs, which occurs rather steadily at a seemingly relatively constant rate throughout most

of the interphase (Dultz & Ellenberg, 2010; Maeshima *et al*, 2010), until reaching the cell's maximal number of NPCs and NBs at the end of G2.

(2) In the HeLa WT cells used in the current study, in which barely any TPR occurs at sites other than the NE, we assumed that the total number of TPR polypeptides occurring within an ordinary cell in late G2 would not likely exceed 270,000, among which there would be only a few TPR polypeptides not bound to the NB (see also Gunkel *et al*, 2021). The largest share of this total number, which we had been able to only estimate at the time when conducting the experiments leading to Figure 1, was the result obtained after having multiplied the approximate number of about 4,000 NPCs in HeLa cells in late G2 (e.g., Maul *et al*, 1972; Ribbeck & Görlich, 2001) with a presumed number of 64 TPR polypeptides associated per NPC on average. Back then, this hypothetical number of 64 represented an assumption based also, but not exclusively, on the following consideration: Even earlier, 32 TPR polypeptides had been determined to occur appended to the NPCs of HeLa nuclear envelopes (Ori *et al*, 2013). These NES, though, had been isolated under conditions that we generally refer to as NB-destabilising, as they can cause up to about half of the NB-appended TPR polypeptides to be detached from such envelopes (see also Gunkel *et al*, 2021). Consequently, we early on considered it possible that, on average, about twice as many TPR polypeptides might occur NE-appended within the HeLa cells' NB-stabilising (NB-s) *in vivo* conditions.

(3) When taking no TPR protein turnover into account, for reasons of simplification, and assuming that most of the TPR polypeptides can be reused after mitosis, the 135,000 TPR polypeptides that will have been distributed to each of the two daughter cells during mitosis will then be largely used up for the post-mitotic assembly of the about 2,000 NBs in early G1, once the same number of NPCs have been assembled slightly earlier. 67,500 of these TPR polypeptides will thereby be such whose association will depend on the presence of ZC3HC1.

From then on, the about 135,000 additional TPR polypeptides required for those NBs to be assembled in interphase, including the 67,500 copies whose NE association is ZC3HC1-dependent, will have to be newly synthesised in each daughter cell during its transition through interphase. Since TPR synthesis appears to occur rather steadily throughout most of the time in interphase, as there is essentially no stockpile of TPR that awaits to be used for NB assembly in interphase, and since the total length of the interphase of the HeLa sublines used for the current study ranges between 19–20 hours, one could estimate that about 6,800-7,000 TPR polypeptides would be synthesised within such a HeLa cell per hour, which thus would represent about 5% of the cell's total TPR amount synthesised during interphase. This amount, in turn, would be equivalent to what would be required for about 100-110 NBs newly assembled within the same time, and again, about 3,400-3,500 of these TPR polypeptides would be used for the ZC3HC1-dependent assembly step.

(4) In the ZC3HC1 KO version of the same progenitor cell line then, the 2000 post-mitotic NPCs would lack the 67,500 TPR polypeptides usually appended in the presence of ZC3HC1, and, in principle, every additional hour into interphase, this number of soluble TPR polypeptides would increase by another 3,400-3,500 copies, until finally reaching the number of 135,000 copies, by the end of G2, whose usual NE-association depends on ZC3HC1.

However, we had already observed by then that the actual amount of TPR in the soluble nucleoplasmic pool could vary notably between different HeLa ZC3HC1 KO sublines and even between individual ZC3HC1 KO cells in some of these KO lines (for details, see Gunkel *et al*, 2021). In the HeLa ZC3HC1 KO cell line used in the current study, we determined the soluble pool of TPR polypeptides to represent, on average, only about 85% of those TPR amounts that are appended to the NPCs in a ZC3HC1-independent manner (e.g., Supplemental Fig S1F). However, also in this HeLa line, some variability could be observed between the individual

ZC3HC1 KO cells, with some harbouring soluble TPR in amounts likely similar to those still appended to the NEs (data not shown, but see also Fig 1D).

(5) At this early point of our investigation, we did not yet know the absolute numbers of ZC3HC1 needed for about half the cell's total amount of TPR to occur NE-appended in HeLa cells, with these numbers eventually determined only later (our unpublished data, to be presented elsewhere). However, quantitative mass spectrometry of the NBs of *Xenopus* oocytes had already indicated by then that ZC3HC1 copy numbers per NB were notably lower, i.e., probably less than half the number of NE-associated TPR polypeptides (Gunkel *et al*, 2021, and our unpublished data). Reckoning that the NBs in the oocytes, at least in the earlier stages of oogenesis, and in HeLa cells are not so fundamentally different with regard to their composition, we thus considered it possible that for re-appending the mean soluble pool of about 115,000 TPR polypeptides back to the NE of a ZC3HC1 KO cell in late G2, less than 115,000 ectopically expressed ZC3HC1 polypeptides, regarding this number as an upper limit value, should suffice, provided that they were indeed involved in such a TPR recruitment process. Since we finally decided to harvest the TPR-sfGFP-expressing ZC3HC1 KO cells at time points no later than 13.5 hours after mitosis (see Fig 1A), and since we were at the same time aiming at re-recruiting back the largest possible amounts of soluble TPR, i.e. those present in cells in which soluble and NE-associated amounts were similar, the total amount of soluble TPR that would then be needed to be re-recruited would comprise 113,400 (i.e., 67,500 + 13.5×3,400). This number, in turn, meant that one would need to provide up to but not more than about 113,000 ectopically expressed ZC3HC1 polypeptides, newly synthesised within a short time, with the latter defined as a period ranging from 30 minutes to certainly not exceeding 150 minutes, in order not to compromise the diagnostic value of the experimental setup.

(6) When addressing the question if it were realistic to reach such ZC3HC1 copy numbers within, for example, one hour after having induced the mCherry-tagged protein's ectopic

expression, we already knew that large amounts of recombinant ZC3HC1 could be rapidly synthesised in transfected HEK293T cells (Gunkel *et al*, 2021). For the latter, we had already approximated the polypeptide copy numbers of ectopically expressed FP-tagged ZC3HC1 to commonly reach more than 100,000 and sometimes exceed 150,000 per hour per transfected cell; with such numbers readily achievable following the transient transfections with mammalian FP-ZC3HC1 expression vectors of about 7 kb in size and the constitutive expression of the recombinant ZC3HC1 polypeptides (Gunkel *et al*, 2021; and our unpublished data). Then, when similarly assessing expression levels in HeLa cells, we found these to be sufficiently high enough, too, for the type of experiment in planning. Even though transfection efficiencies and probably also copy numbers of vectors per cell were lower than for HEK293T, we found that the achievable expression levels within HeLa ZC3HC1 KO cells, transiently transfected with the about 6.5 kb large all-in-one FP-ZC3HC1 expression vector and its strongly inducible Tet promoter, allowed for reaching copy numbers of several tens of thousands of recombinant ZC3HC1 polypeptides per transfected ZC3HC1 KO cell already within less than an hour post-induction. Thereby, these numbers reached and sometimes clearly exceeded the amounts of native ZC3HC1 commonly present in HeLa WT cells (data not shown). Furthermore, such findings were not inconsistent with our initial estimations merely based on published knowledge: Taking mean transcription rates in HeLa cells of 60 nt/sec (Fuchs *et al*, 2014) into account, we had calculated a minimal number of 90 mCherry-ZC3HC1 transcripts per hour and plasmid. Taking further into account the translation rates in different human cell types (e.g., Ingolia *et al*, 2011; Yan *et al*, 2016), and that dozens to hundreds of rounds of translation per single mRNA reportedly occur in cells of lines like HEK293T and U-2 OS (Yan *et al*, 2016), we found that numbers calculated to be in the range of several thousand polypeptides per expression vector per hour were not unreasonable. Finally, it was already known that mammalian expression vectors could end up within the transfected cells' nuclei, like in those from HeLa cells, in large numbers, with such vector copy numbers reportedly ranging from many dozens to thousands per nucleus (see, e.g., Cohen *et al*, 2009; Glover *et al*, 2010).

**Supplemental Information 3. Further considerations relating to experimental benchmarks and data interpretation of the TPR:ZC3HC1:TPR *in vitro* assembly experiment.**

In order to compare starting conditions of the *in vivo* and *in vitro* interaction experiments presented as Figures 1 to 3, and to estimate in how far concentrations of recombinant proteins within the eventually used cell extracts for the *in vitro* assembly experiments might differ from those within the intact cells, we pursued both calculational and experimental approaches.

The calculational one for approximating the mean concentration of soluble sfGFP-TPR within the nuclear interior of a HeLa ZC3HC1 KO cell *in vivo* and of the soluble sfGFP-TPR polypeptides considered assembly-competent within the cell extracts used for the *in vitro* interaction experiments was based on the following considerations and initial values, originating from own findings, published evidence, model assumptions, or estimates. Among these was (i) a mean number of $3\times10^3$ NPCs per HeLa cell in interphase, when from an asynchronous, non-confluent adherent population, and (ii) a 40–45% proportion of all TPR polypeptides within the ZC3HC1 KO cells occurring in a soluble form in interphase, which in turn meant a calculated average of about $8\times10^4$ soluble TPR polypeptides per cell in interphase (see also Supplemental Information 2). In addition to these potentially assembly-competent TPR polypeptides, we (iii) considered it possible that the extracts could also contain several thousand phosphorylated TPR polypeptides, simply because the extracts also included mitotically phosphorylated, and thus assembly-incompetent TPR polypeptides that stemmed from a small number of mitotic cells. The latter had not all been removed during the washes of the adherent populations before subsequent fractionation, with such mitotic cells perhaps still representing a few percent of the total number of cells of the asynchronous populations. In these mitotic ZC3HC1 KO cells, also those TPR polypeptides that had been anchored to the NPCs independently of ZC3HC1 would occur in soluble form, next to the anyhow soluble TPR polypeptides in this cell line. However, assuming that the phosphorylated TPR polypeptides would not compete with the non-phosphorylated ones and thus not interfere with the NB assembly process, we did not take them into account. Furthermore, we (iv) considered the mean total nuclear volume, enclosed by the inner nuclear membrane, to amount to about 700 $\mu m^3$ (Monier *et al*, 2000), of which (v) a fraction of $\leq$300 $\mu m^3$ was estimated to comprise areas freely accessible for soluble proteins, i.e. nucleosol-filled areas neither occupied by the nucleoli nor the various kinds of nuclear bodies, nor nucleosomes and higher-order chromatin arrangements, nor the nuclear lamina and its associated proteins.

Based on these numbers, we calculated a mean concentration of soluble sfGFP-TPR within the nucleosol of the ZC3HC1 KO cells in interphase of about $2.9\times10^5$ polypeptides per picoliter. By contrast, the concentration of the soluble, allegedly assembly-competent sfGFP-TPR polypeptides within the cell extracts used for the *in vitro* interaction experiments presented in the current study was calculated to amount to about $1\times10^4$ to $1.6\times10^4$ polypeptides per picoliter, after having taken into account a total of $8\times10^4$ soluble TPR polypeptides per cell, the total cell numbers commonly used for sonication in buffer solution, and the resulting volumes of the soluble cell fractions obtained after centrifugation (see Material and Methods). As an aside, this value of an about 18- to 29-fold dilution was similar to an about 14-fold dilution, calculated independently of any TPR copy numbers, with the latter dilution values obtained by just setting off against each other (i) the HeLa cell's nucleosol of an estimated 0.3 picoliters (see above), (ii) the HeLa cell's cytoplasmic volume of about 1.6 picoliters (Guillaume-Gentil *et al*, 2016), as part of the HeLa cell's mean total cellular volume of 2.6 picoliters (Zhao *et al*, 2008) including all its insoluble matter, and (iii) a buffer to total cell volume ratio of about 1.8:1 used for preparing such cell extracts (see Material and Methods).

In order to complement these calculational approaches of approximation and to assess the final concentration of the soluble sfGFP-TPR polypeptides within the cell extracts also experimentally, we immunoblotted different amounts of the sfGFP-TPR-containing soluble cell extracts from the ZC3HC1 KO cells, loaded next to total cell extracts as well as next to lamina-NPC-NB (LNN) materials (see Gunkel *et al*, 2021) enriched from defined numbers of sfGFP-TPR-expressing HeLa WT and ZC3HC1 KO cells. Again based on a mean number of $3\times10^3$ NPCs per HeLa cell and at the same time assuming (Supplemental Information 2) that it might be that, on average, 64 and 32 TPR polypeptides, respectively, are located at the NPC of a WT and a ZC3HC1 KO cell, this then allowed for comparing IB signal intensities for such predefined numbers of TPR polypeptides with those present in the soluble cell extracts used for the *in vitro* assembly experiments. Remarkably, such IB signal intensity comparisons allowed determining a concentration of $1\times10^4$ soluble TPR polypeptides per picoliter, representing a value very similar to the one that had been determined by calculational approximation. In other words, in the *in vitro* assembly experiments presented in Figure 3, the sfGFP-TPR polypeptides were used about 30-fold diluted compared to their alleged *in vivo* concentration within the nucleosol of the ZC3HC1 KO cells.

In contrast to the relatively straightforward approximation and comparison of the *in vivo* and *in vitro* amounts of soluble sfGFP-TPR, we had to introduce some simplifying premises when attempting, merely by calculational estimations, to compare the *in vitro* concentrations of mCherry-ZC3HC1, in those extracts to be used for the *in vitro* reconstitution experiments, with some hypothetical values for the *in vivo* concentrations of ectopically expressed mCherry-ZC3HC1 that would be required for reaching a state of equilibrium at the NB.

With hardly any information available till then regarding the ZC3HC1:TPR ensembles' rate constants of association and dissociation, with such constants possibly even differing between subpopulations of ZC3HC1 and TPR and their likely assembly intermediates along the assembly pathway, we aimed at only crudely approximating via highly simplified model assumptions a higher-range value for the *in vivo* concentrations of the ectopically expressed mCherry-ZC3HC1 polypeptides in the Hela ZC3HC1 KO cells presented in Figure 1. The hypothetical value was then to be compared with the easy to determine actual mCherry-ZC3HC1 concentrations within the HEK293T cell extracts that were then to be used for the experiments presented in Figures 2 and 3.

One of the problems was that upon ectopic expression within the HeLa ZC3HC1 KO cells, the recruitment of a large amount of mCherry-ZC3HC1 to the NE appeared to occur shortly after the newly synthesised proteins had been imported into the nucleus, i.e. even before mCherry maturation had been completed (data not shown). Altogether, this rendered it even more difficult to assess any critical concentration of ZC3HC1 polypeptides for NB association from among those steadily changing concentrations within the transfected cells *in vivo* and then compare such value with the single concentration of mCherry-ZC3HC1 polypeptides within the cell extract obtained from an entire population.

Nonetheless, in order to simplify matters and allow for an assessment of whether the *in vitro* concentrations of mCherry-ZC3HC1 obtainable by the mild cell fractionation procedures required for this kind of experiment would be in the range of concentrations encountered in the living cell when first reaching a state of equilibrium between the mCherry polypeptides and the NBs, we prescribed the entirely fictitious situation. In this scenario, the number of mCherry-ZC3HC1 polypeptides first had to reach a distinct *in vivo* concentration that would be stoichiometric to that of the soluble pool of sfGFP-TPR, in order to only then allow for quantitative recruitment of all soluble ZC3HC1 and sfGFP-TPR polypeptides back to the NE.

In those HeLa KO cells that appeared expressing the highest copy numbers of mCherry-ZC3HC1 per time (some presented in Fig 1), such a concentration had been reached, at the latest, within 30 minutes after induction, since all soluble TPR polypeptides had by then been recruited to the NE.

Since we already assumed that at least eight but less than 32 ZC3HC1 polypeptides would be required for the recruitment of 32 TPR polypeptides back to the NE (for the corresponding rationale, see Supplemental Information 2), and since the mean number of soluble sfGFP-TPR polypeptides within the non-transfected cell's nucleus was about $8.6 \times 10^4$ (see further above), we could conclude that at least $2.15–4.3 \times 10^4$ mCherry-ZC3HC1 polypeptides had been synthesised and imported into the nucleus by then. At least such high numbers of polypeptides newly synthesised in such short time were not at all unrealistic, but in principle could even be exceeded, when considering (as outlined in Supplemental Information 2) (i) plasmid numbers per nucleus not uncommon for human tumour cells upon transient transfection (Cohen *et al*, 2009; Glover *et al*, 2010), (ii) transcription speed, (iii) translation speed and (iv) polysome formation within mammalian cells.

Further assuming that the nucleosol-filled areas accessible for soluble proteins within the HeLa ZC3HC1 KO cell nuclei had a mean volume of about 300 $\mu m^3$ (see further above), we calculated the theoretical concentration of such mCherry-ZC3HC1 polypeptides therein to have reached about $7 \times 10^4$ to $1.4 \times 10^5$ polypeptides per picoliter after 30 minutes; always provided, for this hypothetical scenario, that they had not been already gradually removed from the soluble pool by binding to the NE.

The procedure of how to experimentally determine the amounts of mCherry-tagged ZC3HC1 polypeptides within the cell extracts obtained from the transiently transfected HEK293T cell populations, and then, deducible therefrom, also within a transfected cell on average, has in principle already been outlined earlier (Gunkel *et al*, 2021). This approach, using the same cell line and mCherry-ZC3HC1 expression vector also used for the current study, included the essentially lossless, quantitative immunoprecipitation of all mCherry-tagged polypeptides from such extracts, using RFP-specific sdAbs, followed by determining the total amounts of such “nanotrapped” mCherry-ZC3HC1 as it has been described (Gunkel *et al*, 2021). The resulting copy numbers, recurrently determined, had turned out to be relatively proportional to transfection efficiencies and periods of constitutive expression. Now, in the current study, with HEK293T transfection efficiencies of at least 50–70%, periods of 16 to 21 hours during which expression was allowed to take place, average cell population sizes of about $0.7\times10^8$ per 0,5 ml of extract finally to be used for the *in vitro* assembly experiments, yet with only about 25–30% of the cells' total amount of mCherry-ZC3HC1 released into solution (see, for example, Fig 2B) during the brief permeabilisation with digitonin (see Material and Methods), this resulted in final mCherry-ZC3HC1 concentrations, determined by different means, of either $4.7\times10^3$ to at most $1.6\times10^4$ polypeptides per 0.3 picoliters, or of $1.6\times10^4$ to $1.9\times10^4$ copies per 0.3 picoliters. As such, these copy numbers were either close to the same magnitude or one order of magnitude below the estimated total of $4.8\times10^4$ native ZC3HC1 polypeptides that, *in vivo*, would need to have passed through the approximated 0.3 picoliters of freely accessible nucleosol of a HeLa WT cell with its average of $3\times10^3$ NPCs, before eventually having ended up attached at the NE.

However, since *in vivo* incorporation of ZC3HC1 polypeptides turned out to be a continuous process, with no physiological need of first amounting to concentrations as high as in the here prepared cell extracts, the concentrations of mCherry-ZC3HC1 polypeptides within the latter likely exceeded those required anyhow. While it appeared to take slightly longer until all ZC3HC1 attachment sites at the NEs were occupied when cell extracts with the WT version of mCherry-ZC3HC1 had been diluted about 8-fold, for adjusting concentrations to that of the C429S mutant (data not shown, but see Fig 2 and Supplemental Fig S5), the eventually reached occupancy of binding sites, then seemingly in steady state, appeared to be similarly quantitative as with the non-diluted cell extract.

Apart from these considerations concerning copy numbers, we also pondered on the finding that the C429S mutant form of ZC3HC1 was so much less abundant in the final 200,000× *g* cell extract than the WT protein and eventually interpreted it as follows. Since transfection efficiencies with both expression vectors, and corresponding expression levels, appeared to be rather similar, and since relatively more of the mutant than the WT protein was found sedimented by the 20,000× *g* and 200,000× *g* centrifugations, we considered the mutant protein not probably folded and thus prone to aggregation or unspecific binding to other molecules, resulting in more readily sedimentable materials. This interpretation also appeared to be in line with our other observation, namely that the C429S mutant version was found to non-specifically bind, after prolonged incubation, to a variety of subcellular structures within the pre-extracted cells; a feature never observed with the mCherry-tagged WT protein. Among such structures tainted with mCherry-ZC3HC1 C429S were, i.a., interphase chromatin and mitotic chromosomes, kinetochore microtubules and the cytokinetic bridge, and the NE. However, in contrast to when the WT protein was bound to the NE, the mutant protein was not found to attract sfGFP-TPR, neither when located at the NE nor any other structures. Such unspecific binding due to flawed folding might also provide an explanation for why other mutant versions of ZC3HC1, including various deletion mutants, had been reported in the past to engage in interactions with a range of proteins other than TPR and why we could not confirm such interactions being of physiological relevance (Gunkel *et al*, 2021).

**Supplemental Information 4. The multifaceted relationship between TPR and its binding partner NUP153, as one reason for exerting caution when interpreting the relationships between TPR and ZC3HC1.**

Former caution of not frankly assigning ZC3HC1 a direct role in recruiting or keeping TPR polypeptides appended to the NB (Gunkel *et al*, 2021) stemmed from long-existing uncertainties regarding how to interpret the complex relationship between TPR and one of its other binding partners, namely the NPC-appended nucleoporin NUP153.

This latter protein is required for TPR polypeptides of pool 1 to end up attached to the NPCs, both to those assembled directly after mitosis and those formed in interphase (e.g., Walther *et al*, 2001; Hase & Cordes, 2003; Sabri *et al*, 2007; Mendjan *et al*, 2006; Mackay *et al*, 2010; Umlauf *et al*, 2013; Vollmer *et al*, 2015; Larrieu *et al*, 2018), yet with this being so for different reasons. When deficiency of TPR's binding partner NUP153 was achieved by quantitative immunodepletion of NUP153 from *Xenopus* egg extracts that had been made competent for post-mitotic nuclear assembly, or by RNAi-mediated knockdown of NUP153 in living cells, NPCs were still readily assembled, both within such extracts and in the NUP153 RNAi cells towards the end of mitosis (e.g., Walther *et al*, 2001; Hase & Cordes, 2003; Mackay *et al*, 2010; Vollmer *et al*, 2015). TPR, by contrast, could then no longer be found located at such post-mitotic NPCs at all, or only in trace amounts (e.g., Walther *et al*, 2001; Hase & Cordes, 2003; Vollmer *et al*, 2015), thus pointing at some direct role of NUP153 in TPR's recruitment and in mediating the process leading to TPR's appendage to the post-mitotic NPC.

On the other hand, the finding that TPR was also no longer recruited and appended to the NE in interphase when NUP153 was absent likely reflected a more indirect effect because the assembly of the NPC itself is a NUP153-dependent process in interphase (Vollmer *et al*, 2015), which is probably also one of several reasons why NUP153 is an essential protein (e.g., Hart *et al*, 2015; https://www.mousephenotype.org/data/genes/MGI:2385621). The resulting absence of newly assembled interphase NPCs would then simply mean that those TPR polypeptides that are newly synthesised in interphase would find no novel anchor sites.

However, apart from its direct and indirect roles in the recruitment and attachment of TPR to the NPC, it was uncertain for long whether NUP153 would be needed primarily for the NPC's and NB's assembly processes or whether it would also be required for maintaining the integrity of the once assembled NB (e.g., Walther *et al*, 2001). It actually appeared imaginable early on that other NPC proteins, and possibly even other proteins, unknown at that time, could contribute to maintaining and stabilising TPR's localisation at the NB (Hase & Cordes, 2003).

Later findings then, which stemmed from experiments in which cells had been studied at shorter periods after having triggered the NUP153 RNAi process, had indicated that NUP153 might indeed be dispensable for the integrity of a once assembled NB and no longer required by TPR once the latter had been anchored to the NPC (Lussi *et al*, 2010; Duheron *et al*, 2014; our unpublished data). However, while again hinting at some other proteins keeping TPR anchored durably to the NPC, some uncertainty remained with these findings too, as they stemmed from experiments in which most cellular NUP153 had been eliminated, yet, as has been pointed out (Lussi *et al*, 2010), likely not all of it. At the same time, it was still unknown if, and to which extent, a once assembled NB would disassemble again when only some of its fibrils would no longer bind to the NPC whilst others were still attached. Therefore, it did not appear justified at the time to shelve a scenario that would have been equivalent to that of a car wheel that remains attached to a car's axle, even when all but one of the wheel nuts are gone.

Moreover, other data had even suggested that not all parts of those NUP153 polypeptides that pre-existed within a cell before a protein synthesis shutdown needed to be regarded as being degraded and disappearing at the same pace over time. In fact, some NUP153 fragments, namely its NPC- and TPR-binding domain, had been noted to sometimes persevere notably longer than all other parts of the protein, possibly for some remaining purpose (Krull *et al*, 2010, and our unpublished data).

On the other hand, however, while searching for NUP153's and the NB's binding sites at the NPC, we had noted, among a complexity of possibly cooperative interactions, some that were incompatible with NUP153 being an exclusive binding partner for TPR at the NPC proper. Instead, NUP107 too appeared to be one of TPR's binding partners directly at the NPC's NR, in line with findings and conclusions drawn in another study (Kim *et al*, 2014). Furthermore, NUP133 turned out to be in line for a likely second one, which also turned out compatible with yet other findings (Kim *et al*, 2014; Souquet *et al*, 2018). NUP153, on the other hand, turned out to be anchored to at least NUP133 and another protein of the NR's Y-complex, namely NUP160. In fact, we had found NUP153 capable of robustly interacting with NUP160 via a C-terminal segment (NUP160 aa 912-1436), and, in addition, it could bind similarly well to NUP133 aa 479-1156 (our unpublished data). Furthermore, we found this latter interaction between NUP133 and NUP153 abolished when severing NUP133 at sites between aa 934 and 950, and these data, too, were again compatible with related findings (Souquet *et al*, 2018). Altogether, the evidence available to us appeared pointing at an arrangement in which NUP153 polypeptides, anchored to the Y-complex, seem to flank a NUP107- and probably also NUP133-anchored composite NB fibril formed by several TPR polypeptides. As such, the NUP153 polypeptides could, well conceivable, contribute to also stabilising the NB fibril's position via interaction with part of TPR's NBD. On the other hand, though, it also appeared evident in the abovementioned constellation that such interactions between NUP153 and TPR would not be the only ones involved in maintaining the fibril's position once the TPR polypeptides had been appended to the NR.

However, at least in our opinion, the needful clarification regarding NUP153's contribution to maintaining, or not, TPR's positioning a the NPC was only recently provided by a groundbreaking study (Aksenova *et al*, 2020). In the latter, all NUP153 alleles of a human cell line had been CRISPR/Cas9-edited in such a manner that all NUP153 polypeptides synthesised in these cells were degron-tagged, which allowed for their induced degradation and complete elimination within only a few hours. This approach thus permitted more unequivocally correlating the resulting NUP153 deficiency to the effects it was causing at the different time points of the cell cycle. In summary, this approach made it possible to prove that NUP153, while required for NB formation and a process leading to NPC-appendage of TPR, is not the protein required for keeping these TPR pool 1 polypeptides anchored to the NPC.

In the current study, a similar approach, using a cell line expressing degron-tagged ZC3HC1, has now allowed for demonstrating that ZC3HC1, in contrast to NUP153, is required to maintain the NB attachment of the TPR pool 2 polypeptides.

#### Supplemental Figures

**Supplemental Figure S1. Characterisation of a HeLa ZC3HC1 KO cell line expressing TPR C-terminally tagged with sfGFP.**

**(A)** Schematic depiction of the human *TPR* gene and the nucleotide and deduced aa sequences corresponding to exon 51, which in the current study was one of the two target sites of CRISPR/Cas9n-conferred insertion of the ORF for sfGFP into *TPR*. Such tagging was conducted in a HeLa ZC3HC1 KO cell line already characterised in detail earlier (Gunkel *et al*, 2021). The pair of single guide RNA (sgRNA) sequences used for site-specific knock-in are shown in blue lettering except for each of the protospacer adjacent motifs (PAM) highlighted in magenta. In addition, the deduced aa sequences of the non-tagged WT progenitor and the tagged progeny cell line at the transitions between sfGFP-tag and TPR are provided for comparison.

**(B)** Genotyping of the on-target integration of the sfGFP ORF via PCR amplification of genomic regions, with up to four *TPR* alleles (4*) present in the HeLa cell line P2. For primer sequences, see Supplemental Table S5. The PCR products, shown after agarose gel electrophoresis, stemmed from the original, tag-less (0/4*) progenitor HeLa ZC3HC1 KO cell line and the homozygous TPR-sfGFP ZC3HC1 KO cell line in which all *TPR* alleles have been tagged (4/4*). The arrowhead points at the PCR product that includes the 717 bp-long sfGFP ORF. Note, as an aside, that cytological analysis of HeLa P2 had allowed for unambiguously detecting only three copies of chromosome 1, whereas our other data for HeLa P2 indicated the existence of 4 alleles of *TPR*, located at 1q31.1 and here referred to as 4*, with these findings in accord with studies in which 1q had been determined to occur in four copies in other lines of HeLa (e.g., Landry *et al*, 2013; Adey *et al*, 2013).

**(C)** IB of total cell extracts from the tag-less HeLa ZC3HC1 KO progenitor and the same progeny line expressing TPR-sfGFP presented in S1B with all *TPR* alleles tagged. Note that the IB and corresponding Ponceau S-stained membrane shown on the left side lacked the proteins below 70 kDa due to a low-percentage polyacrylamide gel and SDS-PAGE intentionally prolonged for achieving better separation of the non-tagged and tagged TPR polypeptides, with molecular masses of about 267 kDa and 295 kDa, respectively, which were detected by antibodies against TPR. By contrast, SDS-PAGE for the IB and its Ponceau S-stained membrane shown on the right side was performed under standard conditions, followed by incubation with a monoclonal antibody against GFP, to confirm the almost complete absence of truncated versions of sfGFP-tagged TPR and free sfGFP. Traces of the latter (asterisk) could only be detected after prolonged exposure. Arrowheads mark the bands that represent the tagged full-length TPR polypeptides.

**(D)** IFM of HeLa ZC3HC1 KO progenitor cells, expressing non-tagged TPR, that had been grown together on one coverslip with the ZC3HC1 KO progeny cell line expressing all TPR as C-terminally sfGFP-tagged polypeptides. Cells had been cell cycle-synchronised to the G2 phase and then immunolabelled for TPR. Note that in the TPR-sfGFP cell line, the overall intensity of immunolabelling at the NE was very similar to that in the tag-less progenitor line; indicating that the tagged polypeptides' copy numbers at the NE remained largely similar to those in the original cell line and, thus, that the sfGFP tag did not notably affect the NPC-binding of the ZC3HC1-independent population of TPR polypeptides. Also, note that in this particular progeny cell line, the overall appearance of its nucleoplasmic TPR pool, i.e. the one no longer NE-associated, was also similar to the corresponding pool in the progenitor cell line. Bar, 10 µm.

**(E)** Live-cell fluorescence microscopy of cell cycle-synchronised HeLa ZC3HC1 KO cells expressing TPR-sfGFP. Images were taken from populations in late G2, mitosis, and early in

G1. Note that a conspicuous nucleoplasmic pool of TPR-sfGFP is already visible early in G1, indicating that those TPR polypeptides that are seen similarly conspicuously distributed throughout the G2 cells' nuclei would have been readily re-imported into the reassembled nuclei after mitosis. Therefore, such nucleoplasmic TPR does not merely represent polypeptides newly synthesised within the daughter cells. As an aside, also note that TPR in mitosis was not detectable at other structures, like kinetochores, and remained seemingly evenly distributed throughout the mitotic cell's cytoplasm instead. Bar, 10 µm.

**(F)** IB of exemplary cell fractions, obtained from HeLa WT and ZC3HC1 KO cells expressing TPR-sfGFP, to illustrate the existence of a soluble TPR pool in the ZC3HC1 KO cell line. We had chosen this particular ZC3HC1 KO cell line as one among others in which a prominent pool of TPR had been found to persist, distributed throughout the nucleoplasm, even after numerous rounds of cell passaging. This homozygous cell line was the one then used for all of the current study's experiments that involved a HeLa ZC3HC1-KO cell line expressing the C-terminally tagged version of TPR-sfGFP. The corresponding WT cell line, naturally expressing the non-tagged endogenous ZC3HC1 while also homozygously expressing TPR-sfGFP, will be presented in detail in another study. Here, it is only shown for underscoring that a significant, readily extractable soluble pool of TPR is a characteristic of the ZC3HC1 KO cell line. Fractionation of both cell lines was performed in parallel and in a manner allowing for essentially all TPR to remain NE-associated in the ZC3HC1-positive WT cells, which was achieved by using an NB-s buffer that allowed for maintaining the ZC3HC1:TPR interactions at the NE (Gunkel *et al*, 2021). Lanes were loaded with the soluble proteins released during the first and second of two successive fractionation steps (S1 and S2), with the non-soluble LNN-enriched pellet fractions (P), and with the total cell proteins (T) of the non-fractioned cells for comparison. All loadings corresponded to materials obtained from the same number of cells. Immunolabelling with antibodies for TPR and ZC3HC1 was on the upper and lower halves of the Ponceau S-stained membrane shown here. The asterisk marks a minor TPR degradation product. Note that for the KO cells, a major proportion of their total amount of TPR was detectable within their soluble materials (green arrowhead). By contrast, the WT cell line harboured hardly any soluble TPR, with only some trace amounts detectable, possibly including such disassembled in mitotic cells, as the latter had not all been removable by the cells' initial washings.

**Supplemental Figure S2. Characterisation of a HeLa ZC3HC1 KO cell line expressing TPR N-terminally tagged with sfGFP.**

**(A)** Schematic depiction of the human *TPR* gene and the nucleotide and deduced aa sequences corresponding to exon 1, which in the current study was the other target site of CRISPR/Cas9n-conferred insertion of the ORF for sfGFP into the *TPR* alleles in a HeLa ZC3HC1 KO cell line established earlier (Gunkel *et al*, 2021). The pair of sgRNA sequences used for site-specific knock-in are shown in blue lettering except for each of the PAM sequences highlighted in magenta. In addition, the deduced aa sequence of the non-tagged WT progenitor and the tagged progeny cell line at the transitions between sfGFP-tag and TPR are provided for comparison.

**(B)** Genotyping of the on-target integration of the sfGFP ORF via PCR amplification of genomic regions, with four *TPR* alleles present in the HeLa cell line P2. The PCR products, shown after agarose gel electrophoresis, stemmed from the original, tag-less (0/4*) progenitor HeLa ZC3HC1 KO cell line and the homozygous sfGFP-TPR ZC3HC1 KO cell line in which all *TPR* alleles have been tagged (4/4*). The arrowhead points at the PCR product that includes the 717 bp-long sfGFP ORF.

**(C)** IB of total cell extracts from the tag-less HeLa ZC3HC1 KO and the progeny line expressing sfGFP-TPR presented in S2B. SDS-PAGE and IB were performed like for Supplemental Figure S1C. Arrowheads mark the bands representing the sfGFP-tagged TPR polypeptides. The asterisk marks a common TPR degradation product lacking most of the protein's C-terminal domain (e.g., Krull *et al*, 2010; Gunkel *et al*, 2021).

**(D)** IFM of HeLa ZC3HC1 KO progenitor cells, expressing non-tagged TPR that had been grown together on one coverslip with the ZC3HC1 KO progeny cell line expressing all TPR as N-terminally sfGFP-tagged polypeptides. Cells had been cell cycle-synchronised to the G2 phase and then immunolabelled for TPR. Note that the overall intensity of immunolabelling at the NE in the sfGFP-TPR cell line was similar to that in the tag-less progenitor line. Further note that in this progeny cell line, the overall appearance of its nucleoplasmic TPR pool, too, was similar to the corresponding pool in the progenitor ZC3HC1 KO cell line. Bar, 10 µm.

**Supplemental Figure S3. A Tet-On system for doxycycline-inducible ectopic expression of mCherry-tagged versions of ZC3HC1.**

Schematic depiction of the design and functional principle of the mammalian all-in-one expression vector, based on the pTetOne expression system (Heinz *et al*, 2011; https://www.takarabio.com/learning-centers/gene-function/inducible-systems/tet-inducible-systems/tet-one-technology-overview). The latter allows for constitutive expression of the Tet-On 3G transactivator and, in the current study, for doxycycline-inducible expression of the WT or mutant versions of mCherry-tagged ZC3HC1.

**Supplemental Figure S4. An amino acid substitution within the zinc finger domains of ZC3HC1 abolishes its interactions with TPR and the nuclear basket.**

The intactness of the two zinc finger domains of ZC3HC1 is required for its binding to TPR and the NB, as will be presented in detail elsewhere. Here, we already exemplify this by a fluorescence loss in photobleaching (FLIP) experiment, which allowed for comparing the subcellular localisation of the FP-tagged intact protein with its mutant version harbouring the

C429S substitution upon their ectopic expression in HeLa WT and HeLa ZC3HC1 KO cells. When conspicuously overexpressed, the mutant protein and the intact WT polypeptides mainly accumulated within the nucleus as a mobile surplus pool, once binding sites at the NE had been saturated. Bleaching these FP-tagged polypeptides within the cells' nuclear interior then allowed visualising those ZC3HC1 polypeptides that were more lastingly appended to the NE. This approach was chosen to detect polypeptides at the NE that might have remained undetectable due to their signal intensities not exceeding those concomitantly accumulating within the nuclear interior.

**(A, B)** Fluorescence microscopy of populations of transiently transfected HeLa WT cells (A) and HeLa ZC3HC1 KO cells (B), expressing either the intact ZC3HC1 protein, C-terminally tagged with EGFP, or the corresponding C429S mutant, which differs from the WT version by the single aa substitution only. The marked rectangular areas in the pre-bleach image were subjected to pulses of full laser power for 2 min, followed by the acquisition of post-bleach images 1 min after the bleaching, with these images then also shown brightness-enhanced, and with such electronic enhancement only carried out after the image acquisition. Note that the bleaching of the surplus of intact and mutant ZC3HC1 polypeptides deeper within the nuclear interior had resulted in seemingly quantitative elimination of all nuclear EGFP fluorescence and that the NEs of those cells ectopically expressing the intact version of ZC3HC1 were the only structures of the nucleus still fluorescent, pointing at a steady and lasting *in vivo* interaction between the NE and a certain amount of ZC3HC1. By contrast, the ZC3HC1 mutant harbouring C429S was not capable of durably binding to the NE in the ZC3HC1 KO cells, i.e., even not in the absence of any potentially competing intact ZC3HC1 polypeptides. As an aside, also note that the bright cytoplasmic foci (some marked by arrows), seen in HeLa WT and ZC3HC1 KO cells ectopically expressing the vast amounts of the intact ZC3HC1 polypeptides, represented accumulations of ZC3HC1 that were brightly immunostainable for TPR too (data not shown), with such piled-up material evidently no longer amenable for nuclear import. By contrast, some far less signal-intense cytoplasmic foci, only occasionally observed in the C429S-transfected cells, were recurrently found negative for TPR (data not shown). However, how these latter C429S-containing foci (some marked by arrowheads) can come about was not investigated in further detail. We deem it likely, though, that they reflected the mutant polypeptides' tendency, due to their flawed folding, for unspecific interactions with various cellular structures and proteins that do not represent natural binding partners of the intact ZC3HC1 protein (see also Supplemental Information 3). Along this line, most of the less intense cytoplasmic EGFP fluorescence, especially in some of the C429S-expressing cells, originated from sites that appeared dispersed all over the cytoplasmic areas. Therefore, while the bright foci in the cells expressing the intact ZC3HC1 polypeptides unambiguously reflected the mutual immobilisation of subpopulations of ZC3HC1 and TPR polypeptides, as will be presented in detail elsewhere (but also see Figure 6), the attenuated nuclear import of some of the C429S mutant's amounts appeared likely due to some general "stickiness" of this not correctly folded polypeptide. Bar, 10 µm.

**Supplemental Figure S5. The appearance of the NE scaffolds and the relative abundance of ectopically expressed ZC3HC1 used for the assembly experiments.**

The characterisation of the components used for the assembly experiments presented in Figures 2 and 3 also included the visualisation of the morphology of the detergent-permeabilised HeLa ZC3HC1 KO cells, used as the platforms for such *in vitro* assembly experiments. In addition, we regarded it as necessary to assess the cell extracts' amounts of the ectopically expressed mCherry-tagged versions of ZC3HC1 relative to those of the non-tagged endogenous protein.

**(A)** Brightfield and fluorescence microscopy of HeLa ZC3HC1 KO cells before and after extraction with Triton X-100 (TX-100). Cells that had been grown on ibidi µ-slides were washed with assembly buffer and then treated with or without 0.25% (v/v) TX-100 in assembly buffer for 2 min, followed by further washes with assembly buffer. In the course of comprehensive serial tests, such treatment with the detergent had been found sufficient for removing all soluble TPR pool 2 polypeptides from the ZC3HC1 KO nuclei while leaving those of pool 1 bound to the NE. DNA-staining of the cells shown here had been achieved by adding SiR-Hoechst to the culture medium one hour prior to the detergent extraction. Bar, 10 µm.

**(B)** IB of the total proteins of HEK293T cells that had either been mock-transfected or transiently transfected with the expression vector for the mCherry-tagged WT version of ZC3HC1. For estimating amount relationships between the endogenous and tagged version of ZC3HC1, assuming that the tagged and untagged polypeptides were similarly well transferred onto the membranes during Western blotting, the total extracts from the mCherry-ZC3HC1-expressing cells had been loaded as serial dilutions (100 to 0.8%) in parallel. Representative loadings shown here were from cell populations harvested 20 h post-transfection, with a transfection efficiency of approximately 60%. The Ponceau S-stained membrane shown here was used for simultaneously immunolabelling the endogenous and recombinant forms of ZC3HC1, with the former marked by an arrow and the latter with differently-sized arrowheads. Note that the total amount of mCherry-ZC3HC1 in such cell extracts, after having been ectopically expressed overnight, commonly exceeded the cells' endogenous levels approximately 25- to 30-fold, which corresponded in those cells actually transfected to an about 40-fold excess of recombinant over endogenous ZC3HC1 on average. Further note that some amount of the recombinant ZC3HC1, N-terminally tagged with mCherry, was present in a shortened version, here marked by the smaller-sized arrowheads, and since detectable even in the total cell extracts, this meant that such shortening had not been caused during cell fractionation. With no IB-suitable antibody against the very C-terminus of *Hs*ZC3HC1 currently available to us, we could not exclude that some of the shortened polypeptides might reflect C-terminal truncations of ZC3HC1, which might then affect the proper functioning of ZC3HC1 (our unpublished data). However, we regard these shortened polypeptides as more likely explained by the following: The mCherry progenitor protein, *Ds*Red, is known to pass through a state during maturation that is sensitive to heating in SDS-containing protein sample buffer, which results in the protein's cleavage and a major cleavage product that is about seven kDa smaller than the non-cleaved mature protein (Gross *et al*, 2000). Since such an additional "maturation" band has also been described for mCherry (e.g., https://nano-tag.com/products/rfp-selector), we regard it well possible that the here marked minor population mCherry-ZC3HC1 polypeptides, with these too a few kDa smaller than the full-length version of these chimeric polypeptides, reflected a cleaved mCherry tag rather than a ZC3HC1 truncation.

**(C)** IB of the soluble proteins released from digitonin-permeabilised HEK293T cells that had been ectopically expressing either the WT or the C429S mutant version of mCherry-tagged ZC3HC1. Cell extracts loaded for SDS-PAGE had been obtained after 200,000× *g* centrifugation and immunodepletion of trace amounts of soluble TPR. Materials in lanes marked as 1V stemmed from about the same number of permeabilised cells after these had been transiently transfected, in parallel, with the corresponding expression vectors. The WT version-containing extract was also loaded as dilutions (0.5 and 0.1V) to facilitate comparing the amounts of the WT and C429S version relative to each other. Note that the concentrations of the mCherry-tagged WT and C429S mutant proteins were recurrently found to differ to some extent in the final cell extracts, with the mutant version recurrently present in lower amounts, in line with it having turned out to be more prone to unspecific interactions, aggregation, and degradation. This finding was taken into consideration when performing the actual *in vitro* assembly experiments, which also included mutant protein-containing extracts from an about 8-fold higher number of cells than were required for the WT protein, to eventually obtain similar numbers of mCherry-ZC3HC1 polypeptides within the same incubation volumes. However, the amounts of both recombinant versions in the soluble extracts always exceeded those of the endogenous ZC3HC1 nonetheless (see also Supplemental Information 3).

**Supplemental Figure S6. Quantification of the relative amounts of mCherry-tagged ZC3HC1 polypeptides remaining transiently and lastingly NE-associated upon the initial binding to the NE scaffolds of ZC3HC1 KO cells.**

Upon adding cell extracts containing mCherry-ZC3HC1 in excess and following the initial binding of such recombinant ZC3HC1 to the detergent-permeabilised NE scaffolds of the ZC3HC1 KO cells, we had recurrently noted that only a certain proportion of the added ZC3HC1 polypeptides remained lastingly NE-associated. In contrast, others appeared to only transiently interact with the NEs, with this observation illustrated in the following (A, B). Subsequent relative quantifications (C) revealed that the transiently and the more durably NE-associated ZC3HC1 polypeptides represented two subpopulations of about similar size.

**(A)** Schematic depiction of the incubation of TX-100-permeabilised HeLa ZC3HC1 KO cells, cleared of soluble TPR polypeptides, with the mCherry-tagged WT version of ZC3HC1 from digitonin-permeabilised HEK293T cells, obtained after high-speed centrifugation and cleared of TPR, followed by washes in assembly buffer and the cells' incubation therein for 12.5 h.

**(B)** Fluorescence microscopy of the detergent-extracted ZC3HC1 KO cells that were first incubated with the WT ZC3HC1-containing cell extract for 30 min, here with an extract in which the concentration of mCherry-ZC3HC1 was 8× higher than in experiments presented in Figure 2, followed by three quick washes with assembly buffer, and the cells' further incubation therein. Specimens were briefly inspected at indicated time points, thereby keeping exposure to laser light as short as possible. The monochromatic images are also shown colour-graded to display differences in pixel intensities via a colour LUT. Note that after having washed the cells and then having kept them in the buffer for only a short time, a certain amount of mCherry-ZC3HC1 had been lost again from the NE. Such an observation was made recurrently, and in the examples presented here, such signal diminishment is apparent when comparing the micrograph for the "loaded" specimen with the one acquired 30 min after washing. From that point on, however, the remaining NE-associated mCherry was found lastingly positioned there, as it is evident when comparing the pictures taken after 30 min and up to 12.5 h. When inspecting the colour-graded images, the amount of mCherry released from the NE and the other remaining NE-associated both appeared to represent about half of the total amount initially attached to the NE. Bar, 10 µm.

**(C)** Quantification of signal yields for mCherry-ZC3HC1 at the NEs of the ZC3HC1 KO cells prior to and at different time points after cell extract removal and incubation in assembly buffer only. Randomly chosen NE segments for quantifications via ImageJ were from essentially all labelled nuclei (n = 115 in total, with at least 13 nuclei per time point) seen in equatorial view within randomly chosen images of these cells. The graph displays the NEs' relative signal intensity values measured at the indicated time points, with the arithmetic mean values marked by black squares, with the one for the first time point (-5 min) set to 100%, and with the standard deviations provided. Note that the mean signal yield for the mCherry-ZC3HC1 present at the KO cells' NEs in the presence of an excess of mCherry-ZC3HC1, while soluble TPR polypeptides were absent, was about twice as high as after the same NEs had been washed with assembly buffer and incubated therein for longer times. While work aiming at a detailed explanation for such an observation goes beyond the scope of the current study, we here consider it possible that such a step-wise loss of a specific amount of mCherry-ZC3HC1 on the one hand, and the persistent NE-association of a second proportion, on the other hand, points to at least two, then similarly large pools of ZC3HC1 polypeptides possibly engaging in different types of interactions at the NB. Future work will need to clarify whether these

ZC3HC1 subpopulations might bind with differing affinities to different sites of the NPC-anchored TPR polypeptides of pool T1, either naturally or merely as a consequence of having treated the NEs with TX-100, or whether ZC3HC1 polypeptides can homodimerise. In such latter case, one could imagine the one pool tightly bound directly to the TPR 1 polypeptides, and the second associated to the first one only loosely in the absence of T2 polypeptides.

**Supplemental Figure S7. Illustration of the sfGFP$^{\mathbf{L9mIAA7}}$-tag and the subcellular distribution of *At*AFB2 in an HCT116 master cell line.**

**(A)** Crystal structure of the sfGFP (https://www.rcsb.org/structure/2B3P; Pédelacq *et al*, 2006), shown with and without an AID-tag (here mIAA7), the latter schematically depicted not to scale. The nt coding sequence for mIAA7 has been inserted in-frame into the sfGFP sequence coding for loop 9 (L9), between aa175 and aa176, with this position having already been shown suitable for peptide insertions earlier (Abedi *et al*, 1998; Wang *et al*, 2014).

**(B)** Integration of each one of two different *At*AFB2 expression cassettes into the AAVS1 safe-harbour locus within the *PPP1R12C* gene on chromosome 19 of the near-diploid HCT116 cell line, using the sgRNA/Cas9 vector T2 (Natsume *et al*, 2016; Addgene plasmid #72833), for targeting the AAVS1 locus. Under the control of the EF-1α promoter, the *At*AFB2 cassette, based on the formerly described pSH-EFIRES-P-*At*AFB2 cassette (Li *et al*, 2019; Addgene plasmid #129715), allowed for constitutive expression of the auxin signalling F-box 2 protein AFB2 gene from *Arabidopsis thaliana.* In addition, we had tagged the ORF for *At*AFB2 with the nt sequence for the Myc-tag epitope sequence EQKLISEEDL to allow for the chimeric protein's detection by antibodies (rabbit mAb 71D10). Furthermore, we had appended the nt sequence of the nuclear localisation signal (NLS) of c-Myc (Dang & Lee, 1988) to one of the two expression cassettes' *At*AFB2-Myc sequence to allow for the chimeric protein's nuclear import. The already present internal ribosome entry site (IRES; Li *et al*, 2019), from the encephalomyocarditis virus, allowed for the translation of the puromycin resistance (puroR) gene. The left (LHA) and right homology arms (RHA) required for cassette integration via homology-directed repair are indicated.

**(C)** IFM of three different HCT116 cell lines, with the one having the NLS-free *At*AFB2-Myc cassette homozygously inserted into both AAVS1 loci (micrographs on the left), the other harbouring the *At*AFB2-Myc-NLS cassettes in both loci (right side), and the third possessing both an NLS-free and an NLS-tagged version of *At*AFB2 (centre). Note that the NLS-lacking version of *At*AFB2 was largely excluded from the nucleus and distributed throughout the cytoplasm instead, while *At*AFB2 was near-exclusively located within the nuclear interior in the cell line expressing only the NLS-tagged version. By contrast, in the HCT116 cell line selected as the current study's master cell line (MCL) for all degron experiments, the *At*AFB2 polypeptides were distributed throughout the cytoplasm and nuclear interior, as a result of *At*AFB2 in this cell line being expressed both with and without the NLS. Bar, 10 µm.

**Supplemental Figure S8. Characterisation of an HCT116 cell line expressing sfGFP$^{L9mIAA7}$-tagged TPR.**

**(A)** Schematic depiction of the human *TPR* gene and the nucleotide and deduced aa sequences corresponding to exon 1, which in the current study was the target of CRISPR/Cas9n-conferred insertion of the ORF for sfGFP in combination with the AID-tag mIAA7. The sequences of the pair of sgRNAs used for site-specific knock-in are shown in blue lettering, with the PAM additionally highlighted in magenta. The deduced aa sequences of the non-tagged WT progenitor and the tagged progeny cell line at the transition between the sfGFP$^{L9mIAA7}$-tag and TPR are provided for comparison. As an aside, note that only the first and last four aa of the sfGFP sequence are shown, and the mIAA7 sequence has been inserted into loop 9 of sfGFP.

**(B)** Genotyping of the on-target integration of the sfGFP$^{L9mIAA7}$ ORF via PCR amplification of genomic regions, with two *TPR* alleles present in the nearly diploid HCT116 cell line. The PCR products stemmed from the tag-less HCT116 MCL progenitor cell line (0/2) and the progeny line in which both TPR alleles have been tagged (2/2). The arrowhead points at the PCR product with the homozygously integrated, 1029 bp-long sfGFP$^{L9mIAA7}$ ORF.

**(C)** IB of total cell extracts from the tag-less HCT116 MCL progenitor cell line and the same progeny line expressing sfGFP$^{L9mIAA7}$-TPR presented in S8B, with both TPR alleles tagged. SDS-PAGE and immunoblotting for TPR and GFP were like for Supplemental Figure S1C, with additional IB for ZC3HC1 performed for further comparison. Arrowheads mark the bands representing the sfGFP-tagged TPR polypeptides, and the asterisk marks the common TPR degradation product lacking most of its C-terminal domain.

**Supplemental Figure S9. Characterisation of HCT116 cell lines expressing sfGFP$^{AID}$-tagged ZC3HC1.**

**(A)** Schematic depiction of the human *ZC3HC1* gene and the nucleotide and deduced aa sequences corresponding to exon 10, which in the current study was the target of CRISPR/Cas9n-conferred insertion of the ORF for sfGFP in combination with an AID-tag. The sequences of the pair of sgRNAs used for site-specific knock-in are shown in blue lettering, with the PAM additionally highlighted in magenta. Furthermore provided are the aa sequences at the transition between ZC3HC1 and the sfGFP$^{AID}$-tag of two of the eventually tagged progeny cell lines, with one having the mIAA7 degron sequence appended to sfGFP's C-terminus and the other with the mIAA7 sequence inserted into loop 9 of sfGFP. Again, only the first and last four aa of the sfGFP sequence are shown.

**(B)** Genotyping of the on-target integration of the sfGFP$^{L9mIAA7}$ ORF via PCR amplification of genomic regions, with two *ZC3HC1* alleles present in the nearly diploid HCT116 cell line. The

PCR products stemmed from the tag-less HCT116 MCL progenitor cell line (0/2) and the two progeny lines in which both ZC3HC1 alleles have been tagged (2/2) with either sfGFP$^{CTmIAA7}$ or sfGFP$^{L9mIAA7}$. The arrowhead points at the PCR products with the homozygously integrated, 981 bp-long sfGFP$^{CTmIAA7}$ ORF and the 1029 bp-long sfGFP$^{L9mIAA7}$ ORF.

**(C)** IB of total cell extracts from the tag-less HCT116 WT cell line, the HCT116 MCL progenitor, and the two progeny lines expressing either ZC3HC1-sfGFP$^{CTmIAA7}$ or ZC3HC1-sfGFP$^{L9mIAA7}$, each with both *ZC3HC1* alleles tagged and also presented in S9B. Immunoblotting was with an antibody targeting the Myc-tag of the correspondingly tagged *At*AFB2 protein and with antibodies for ZC3HC1, with additional IB for TPR performed for further comparison.

**Supplemental Figure S10. Progression of auxin-induced degradation of TPR and ZC3HC1 in homozygous cell lines expressing different versions of the sfGFP$^{AID}$-tag.**

(**A**) Fluorescence microscopy of cells of the HCT116 cell line expressing all TPR polypeptides N-terminally tagged with sfGFP$^{L9mIAA7}$. Images were acquired from cells that had been harvested at indicated time points before and after the addition of auxin, with such cells then briefly fixed with FA, permeabilised with TX-100, and DNA-stained with Hoechst 33342. The micrographs with the GFP signals are shown colour-graded, with any colouring in green corresponding to no signal at all, to facilitate the assessment of the degree of degron-mediated elimination of the sfGFP$^{L9mIAA7}$-tagged polypeptides. Note that almost complete elimination of the NE-associated sfGFP$^{L9mIAA7}$-tagged TPR polypeptides was already observed 60 min after adding auxin, with essentially complete degradation by 90 min, except for some rare laggards (here marked by white arrowheads). Further note that cells presented in S10A and S10C were analysed in parallel. Bar, 25 µm.

**(B)** Crystal structure of the sfGFP (see also Supplemental Fig S7A) without an AID-tag, and with the schematically depicted AID-tag mIAA7 appended to either sfGFP's C-terminus or inserted into loop 9.

**(C)** Fluorescence microscopy of cells of the HCT116 cell line expressing all ZC3HC1 polypeptides C-terminally tagged with sfGFP and the AID sequence of mIAA7 either appended to the CT of sfGFP or inserted into sfGFP's loop 9. Treatment of cells and image acquisition was like in S10A. Note that elimination of the NE-associated sfGFP$^{L9mIAA7}$-tagged ZC3HC1 polypeptides after the addition of auxin was more rapidly achieved with the mIAA7 sequence inserted into the loop 9 of sfGFP than when appended to its CT. While substantial amounts of the ZC3HC1-sfGFP$^{CTmIAA7}$ polypeptides were still visible after incubation with auxin for 120 min, the sfGFP$^{L9mIAA7}$-tagged version of ZC3HC1 was almost quantitatively degraded at this time point. Bar, 25 µm.

**Supplemental Figure S11. Auxin-induced TPR degradation in the homozygous sfGFP$^{L9mIAA7}$-TPR cell line and the effects on other TPR binding partners.**

IFM of cells from the HCT116 progenitor MCL expressing the naturally tag-free TPR and of cells from the homozygous HCT116 progeny line, in which all TPR polypeptides were N-terminally tagged with sfGFP$^{L9mIAA7}$. Cells had been co-cultured as mixed populations on the same coverslip, cell cycle-synchronised, and then treated with either DMSO for control or with auxin for different lengths of time, with cells shown here having been harvested after incubation of four hours. In order to make some of the antibodies' epitopes better accessible, cells had been pre-extracted with TX-100 in the presence of $MgCl_2$, the latter for maintaining NB integrity in the control cells. To allow for the comparability of the results, we treated all specimens shown here in the same manner, which, in turn, also meant that the soluble nuclear pools of all NB components released upon TPR degradation were no longer present. Specimens were then double-labelled with combinations of antibodies for TPR, ZC3HC1 and the other TPR binding partners MAD1, GANP, SENP1, and NUP153. Note first, the complete elimination of NE-associated GFP and of essentially all TPR immunostaining in those cells expressing TPR-sfGFP$^{L9mIAA7}$, which was accompanied by ZC3HC1 being no longer detectable at the NE, while the progenitor cell's TPR and ZC3HC1 immunostainings remained unaffected. Note further that MAD1, GANP, and SENP1 polypeptides, too, had been detached en masse from the NE upon such TPR degradation, while NE-association of NUP153 appeared mostly unaffected at this time point. In contrast to ZC3HC1 and MAD1, though, traces of immunolabelling for GANP and SENP1 at the NEs were still visible at this time point, several hours after the quantitative removal of all TPR. Bar, 10 µm.

**Supplemental Figure S12. Temporarily maintained positioning at the NE of some NB-associated proteins, exemplified by GANP, in the absence of TPR.**

The rapid degron-mediated elimination of TPR and other NPC-associated proteins allows, in principle, for unveiling a hierarchy of complex interactions between NB and NPC components because this approach of eliminating a target protein, unlike its KD by RNAi or KO by CRISPR/Cas9-methodology, allows for a close spatiotemporal correlation between the target protein's removal and the effects this has on other proteins. For example, while RNAi of TPR is known to eventually also result in all NPC-associated GANP being no longer positioned there, demonstrating that TPR plays a central role in keeping GANP placed at the nuclear periphery (e.g., Umlauf *et al*, 2013; Wickramasinghe *et al*, 2014; our unpublished data), the degron-mediated elimination of TPR allows for unveiling further interactions, beyond those with TPR, that contribute to GANP's positioning at the NE, as demonstrated in the following. **(A)** IB of cell extracts obtained from sfGFP$^{L9mIAA7}$-TPR HCT116 cells, after these had been treated with auxin, or with DMSO only, for one hour. Similar numbers of cells of the two differently treated populations had then been fractioned in parallel, resulting each in a fraction of soluble proteins (S) and a corresponding pellet fraction (P) also containing the NPCs and normally, i.e. in the absence of auxin, the complement of NB proteins. The IBs shown here for TPR, ZC3HC1, and NUP153, and the Ponceau S-stained membrane are identical to the ones presented in Figure 4B. Note that the same membrane initially used for the immunolabelling of TPR and ZC3HC1 was reused for the additional immunolabelling of GANP. Further note that at this time point of the treatment with auxin, when TPR was already no longer detectable and when ZC3HC1 had been quantitatively released into solution, GANP was still present, almost exclusively, within the NPC-NB-enriched pellet fraction.

**(B)** IFM of cells from the HCT116 progenitor MCL expressing the naturally tag-free TPR and of cells of the homozygous HCT116 progeny line, in which all TPR polypeptides were N-terminally tagged with sfGFP$^{L9mIAA7}$. Cells had been co-cultured as mixed populations on the same coverslip and cell cycle-synchronised. They were then harvested after having been treated, or not, with auxin for one, two and four hours, with auxin having been added first, second and last to those cells that were to be incubated for four, two, and one hour, respectively. All specimens of this time course were harvested and processed together and then analysed in parallel, using identical microscope settings. Note the nearly complete elimination of NE-associated GFP and TPR immunostaining, accompanied by ZC3HC1 no longer detectable at the NE after already one hour of auxin-treatment. GANP, by contrast, was still located primarily at the NE at this time point, with some variation in the intensities of NE-associated immunolabelling for GANP also notable in those cells only expressing the naturally non-tagged version of TPR. GANP's gradual detachment from the NE was only notable at later time points, indicating that other proteins, apart from TPR, also contribute to keeping GANP positioned at the NE. Bar, 10 µm.

### Supplemental Tables

#### Supplemental Table S1: Antibodies.

| Antibody (target region) | Source |
|---|---|
| Guinea pig polyclonal anti-*Hs*NUP153 (21–36) | (Ferrando-May *et al*, 2001) |
| Guinea pig polyclonal anti-*Hs*NUP153 (1459–1475) | (Krull *et al*, 2010) |
| Guinea pig polyclonal anti-*Hs*TPR (2063–2084) | (Cordes *et al*, 1997) |
| Guinea pig polyclonal anti-*Hs*ZC3HC1 (307–355) | (Gunkel *et al*, 2021) |
| Mouse monoclonal anti-*Hs*LMNB2, clone X223 (96–226) | (Höger *et al*, 1990; Schumacher *et al*, 2006) |
| Mouse monoclonal anti-*Hs*TPR, clone 203-37 (1462–1500) | (Cordes *et al*, 1997; Hase *et al*, 2001; Gunkel *et al*, 2021) |
| Mouse monoclonal anti-*Hs*ZC3HC1, clone B-10 (1–60*) | Santa Cruz Biotechnology (Dallas, TX, USA; sc-365058)<br>* epitope mapped to aa 1-60 (Gunkel *et al*, 2021) |
| Mouse monoclonal anti-*Hs*MAD1 (481–718*), clone BB3-8 | Merck Millipore (Burlington, MA, USA; MABE867)<br>* region mapped in the course of this study |
| Rabbit monoclonal anti-*Hs*SENP1, clone EPR3844 | Abcam (Cambridge, United Kingdom; ab108981) |
| Rabbit monoclonal anti-Myc-tag, clone 71D10 | Cell Signaling Technology (Danvers, MA, USA; #2278) |
| Rabbit polyclonal anti-*Hs*GANP (1000–1012) | This study |
| Rabbit polyclonal anti-*Hs*NUP153 (50–100) | Abcam (ab84872) |
| Rabbit polyclonal anti-*Hs*TPR (77–94) | (Gunkel *et al*, 2021) |
| Rabbit polyclonal anti-*Hs*TPR (93–110) | This study |
| Rabbit polyclonal anti-*Hs*TPR (381–398) | This study |
| Rabbit polyclonal anti-*Hs*TPR (390–407) | This study |
| Rabbit polyclonal anti-*Hs*TPR (2147–2163, S2155P) | This study |
| Rat monoclonal anti-GFP, clone 3H9 | ChromoTek (Planegg-Martinsried, Germany; 3h9) |
| Donkey polyclonal anti-rat IgG (H&L, minimal cross reactions) conjugated with HRP | Jackson ImmunoResearch (Cambridgeshire, United Kingdom; 712-035-150) |
| Donkey polyclonal anti-guinea pig IgG (H&L, minimal cross reactions) conjugated with Alexa488, Cy3, Cy5 or HRP | Jackson ImmunoResearch (706-545-148, 706-165-148, 706-175-148 or 706-035-148) |
| Donkey polyclonal anti-mouse IgG (H&L, minimal cross reactions) conjugated with Alexa488, Cy3, Cy5 or HRP | Jackson ImmunoResearch (715-545-150, 715-165-150, 715-175-150 or 715-035-150) |
| Donkey polyclonal anti-rabbit IgG (H&L, minimal cross reactions) conjugated with Alexa488, Cy3, Cy5 or HRP | Jackson ImmunoResearch (711-545-152, 711-165-152, 711-175-152 or 711-035-152) |
| FluoTag-X4 anti-RFP/mCherry sdAb conjugated with Atto565 | NanoTag Biotechnologies (Göttingen, Germany; N0404) |

#### Supplemental Table S2: Cell lines.

| Cell line | Source |
|---|---|
| HCT116 | RRID:CVCL_0291 (CCL-247; ATCC, Manassas, VA, USA) |
| HCT116 *At*AFB2-Myc | This study |
| HCT116 *At*AFB2-Myc-NLS | This study |
| HCT116 *At*AFB2-Myc/atAFB2-Myc-NLS = MCL | This study |
| HCT116 sfGFP$^{L9mIAA7}$-TPR | This study |
| HCT116 ZC3HC1-sfGFP$^{L9mIAA7}$ | This study |
| HCT116 ZC3HC1-sfGFP$^{CTmIAA7}$ | This study |
| HEK293T | RRID:CVCL_0063 (CRL-3216; ATCC) |
| HeLa P2 | (Gunkel *et al*, 2021) |
| HeLa P2 ZC3HC1 KO | (Gunkel *et al*, 2021) |
| HeLa P2 ZC3HC1 KO TPR-sfGFP | This study |
| HeLa P2 ZC3HC1 KO sfGFP-TPR | This study |

**Supplemental Table S2:** Hela cells of subline Hela P2, all CRISPR/Cas9n-edited progeny lines of HeLa P2, and HEK293T cells were cultured in high-glucose DMEM (D6429, Sigma-Aldrich, St. Louis, MO, USA) with 10% FBS (P40-37500; PAN-Biotech, Aidenbach, Germany). For the HEK293T cells, 2 mM L-glutamine (GIBCO, Grand Island NY, USA) was additionally added to a final concentration of 6 mM. HCT116 cells and the CRISPR/Cas9-edited progeny lines were grown in McCoy's 5A medium (M9309, Sigma-Aldrich) with 10% FBS. All lines were cultivated in a humidified atmosphere with 5% $CO_2$, at a temperature of 37°C.

#### Supplemental Table S3: Mammalian expression vectors.

| Expression construct | Backbone (resistance) | Source |
|---|---|---|
| Dox-inducible mCherry-*Hs*ZC3HC1(1-502) | pTetOne (Amp) | This study |
| Dox-inducible mCherry-*Hs*ZC3HC1(1-502 \| C429S) | pTetOne (Amp) | This study |
| mCherry-*Hs*ZC3HC1(1-502) | pEYFP-C1 (Kan) | This study |
| mCherry-*Hs*ZC3HC1(1-502 \| C429S) | pEYFP-C1 (Kan) | This study |
| *Hs*ZC3HC1(1-502)-EGFP | pEGFP-N1 (Kan) | This study |
| *Hs*ZC3HC1(1-502 \| C429S)-EGFP | pEGFP-N1 (Kan) | This study |
| h*Sp*Cas9n(BB)-2A-Puro | PX462 V2.0 (Amp) | (Ran *et al*, 2013), gift from Feng Zhang (Addgene plasmid # 62987) |
| Integration vector *At*AFB2-Myc_IRES_puroR | pSH-EFIRES-P (Amp) | Modified Addgene plasmid #129715, gift from Elina Ikonen (Li *et al*, 2019) |
| Integration vector *At*AFB2-Myc-NLS_IRES_puroR | pSH-EFIRES-P (Amp) | Modified Addgene plasmid #129715, gift from Elina Ikonen (Li *et al*, 2019) |
| Integration vector TPR-sfGFP | pQE80N (Kan) | This study |
| Integration vector sfGFP-TPR | pQE80N (Kan) | This study |
| Integration vector sfGFP$^{L9mIAA7}$-TPR | pQE80N (Kan) | This study |
| Integration vector ZC3HC1-sfGFP$^{L9mIAA7}$ | pQE80N (Kan) | This study |
| Integration vector ZC3HC1-sfGFP$^{CTmIAA7}$ | pQE80N (Kan) | This study |
| sgRNA NT1 (*Hs*TPR) / h*Sp*Cas9n(BB)-2A-Puro | PX462 V2.0 (Amp) | This study |
| sgRNA NT2 (*Hs*TPR) / h*Sp*Cas9n(BB)-2A-Puro | PX462 V2.0 (Amp) | This study |
| sgRNA CT1 (*Hs*TPR) / h*Sp*Cas9n(BB)-2A-Puro | PX462 V2.0 (Amp) | This study |
| sgRNA CT2 (*Hs*TPR) / h*Sp*Cas9n(BB)-2A-Puro | PX462 V2.0 (Amp) | This study |
| sgRNA NT1 (*Hs*ZC3HC1) / h*Sp*Cas9n(BB)-2A-Puro | PX462 V2.0 (Amp) | (Gunkel *et al*, 2021) |
| sgRNA NT2 (*Hs*ZC3HC1) / h*Sp*Cas9n(BB)-2A-Puro | PX462 V2.0 (Amp) | (Gunkel *et al*, 2021) |
| sgRNA CT1 (*Hs*ZC3HC1) / h*Sp*Cas9n(BB)-2A-Puro | PX462 V2.0 (Amp) | This study |
| sgRNA CT2 (*Hs*ZC3HC1) / h*Sp*Cas9n(BB)-2A-Puro | PX462 V2.0 (Amp) | This study |
| sgRNA T2 (*Hs*AAVS1 safe-harbour) / pSpCas9(BB) | PX330 (Amp) | (Natsume *et al*, 2016), gift from Masato Kanemaki (Addgene plasmid # 72833) |

#### Supplemental Table S4: sgRNA Sequences

| Description | Vector backbone | gRNA sequence (sense) | Purpose |
|---|---|---|---|
| *Hs*AAVS1 sgRNA T2 | PX330 | GGGGCCACTAGGGACAGGAT | safe-harbour locus integration |
| *Hs*TPR sgRNA NT1 | PX462 V2.0 | gTCTCGGATTGCTGATCAGCA | sfGFP integration (NT, Exon 1) |
| *Hs*TPR sgRNA NT2 | PX462 V2.0 | GGGGCGGCATGAGAAATTTA | sfGFP integration (NT, Exon 1) |
| *Hs*TPR sgRNA CT1 | PX462 V2.0 | gCTCCTCTCCCTCCCATTGCA | sfGFP integration (CT, Exon 51) |
| *Hs*TPR sgRNA CT2 | PX462 V2.0 | gCAGAGGAAATATTAATTAAA | sfGFP integration (CT, Exon 51) |
| *Hs*ZC3HC1 sgRNA NT1 | PX462 V2.0 | GCAGTAGTTCGCTCCCCAGA | knock-out (Exon 1) |
| *Hs*ZC3HC1 sgRNA NT2 | PX462 V2.0 | GCAAACGCTTGTCCCTCACA | knock-out (Exon 1) |
| *Hs*ZC3HC1 sgRNA CT1 | PX462 V2.0 | GATTCCCACTGCCGAAATATT | sfGFP integration (CT, Exon 10) |
| *Hs*ZC3HC1 sgRNA CT2 | PX462 V2.0 | GAAGATACTCCAGCGCCTTCC | sfGFP integration (CT, Exon 10) |

**Supplemental Table S4:** Oligos (purchased by Sigma-Aldrich; including overhangs not shown here that are compatible with the digested vector) were annealed and inserted into a BbsI-digested bicistronic vector for gRNA/Cas9n according to the published protocol (Ran *et al*, 2013). PAM sequence is not included here. For higher expression levels of guide RNAs with the human U6 promoter, a 'G' was inserted at the transcription start site if not naturally occurring there. The AAVS1 sgRNA sequence was published earlier, and the plasmid was obtained via Addgene from Masato Kanemaki (Natsume *et al*, 2016; see also Supplemental Table S3).

#### Supplemental Table S5: Primers for genotyping.

| Target | Forward primer | Reverse primer |
|---|---|---|
| *Hs*TPR-NT | TGGCAGACAATTACCTCGGACGGTC | CAAGGACGGAAAGAGCATCTCACTTAC |
| *Hs*TPR-CT | CCATTTACCCTTTTTCGGTTTGTTAAATG | CATTGATGTGCCTAGTAGAACAGCAAGAG |
| *Hs*ZC3HC1-NT | CCCTGCTGAATTTGGACTGACAGCG | CTGCGGTTGGGAATCTGGTACACC |
| *Hs*ZC3HC1-CT | GTCCTAGACATATCCAGATCATTGTGGG | CTCAGTGACGAAAAACATGGCATATCAGAG |

**Supplemental Table S5:** The oligos used for genotyping of the generated cell lines were purchased from Sigma-Aldrich. Restriction enzyme cleavage sites (not shown here) were added in order to subclone PCR products for downstream sequencing.

Supplemental Figure S1

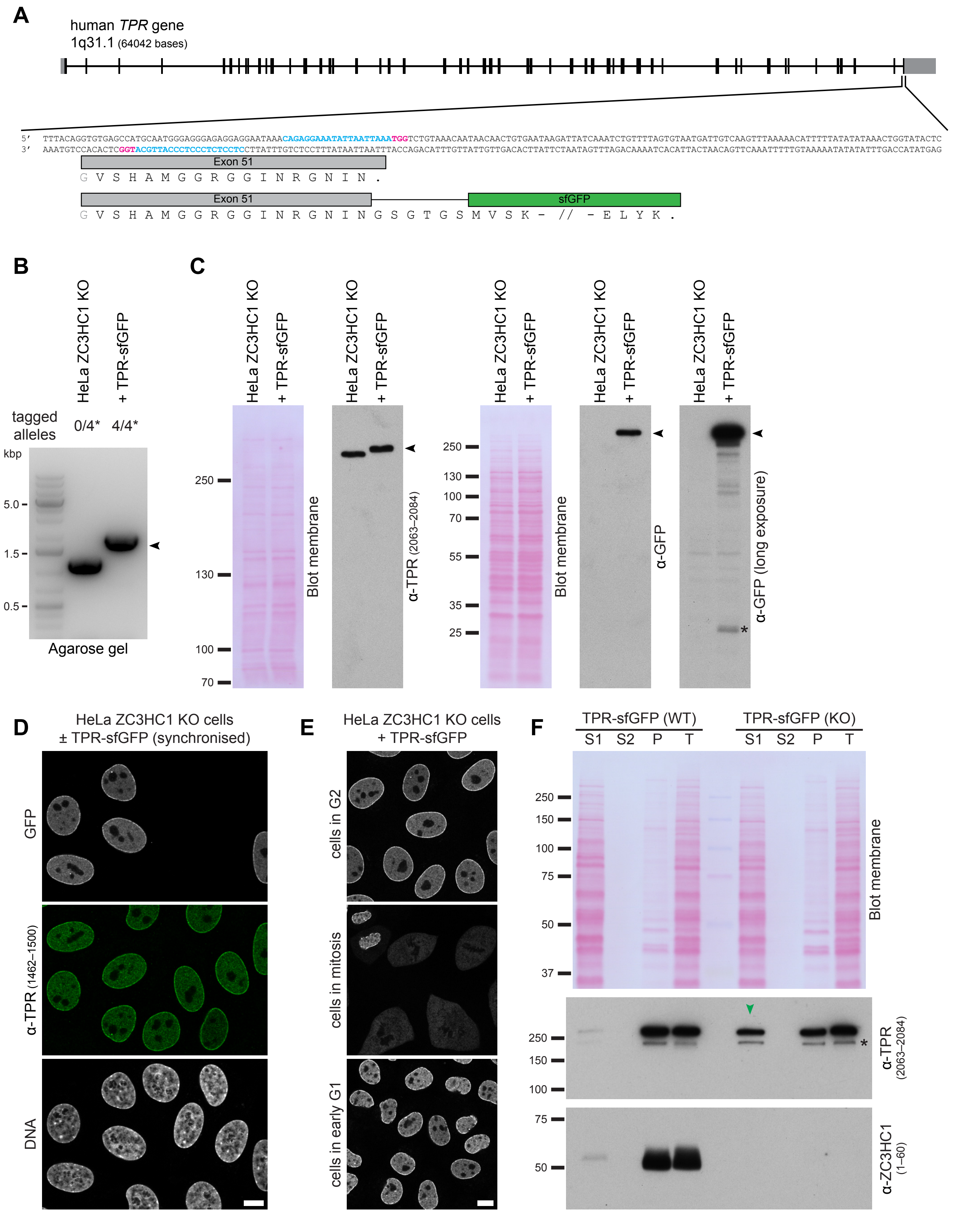

Supplemental Figure S2

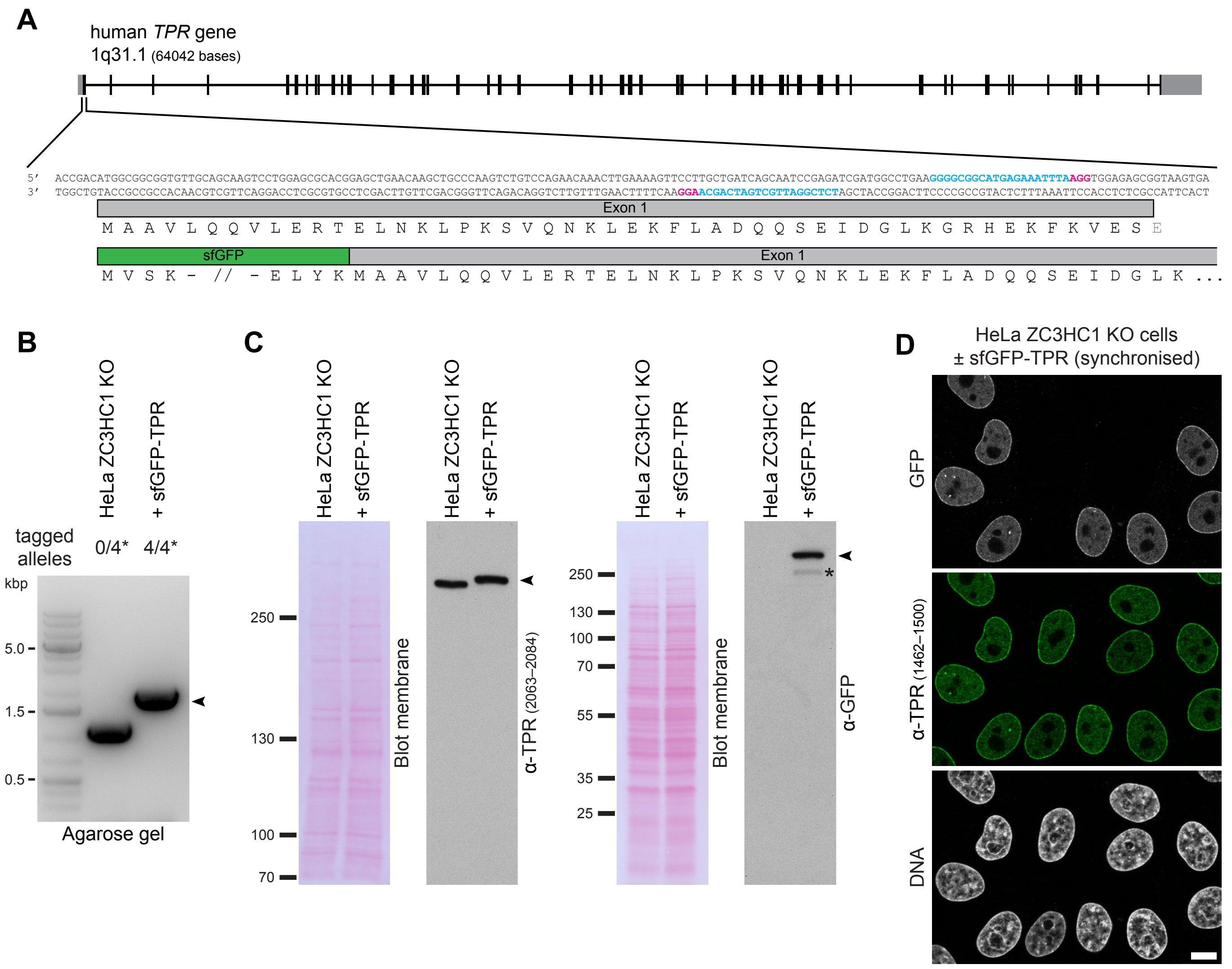

Supplemental Figure S3

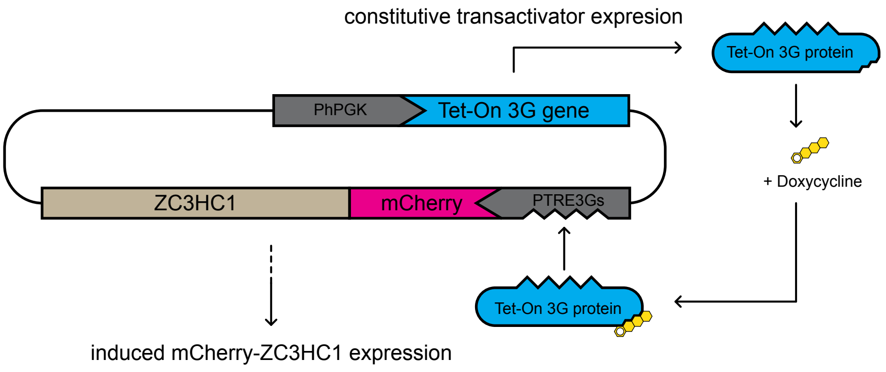

### Supplemental Figure S4

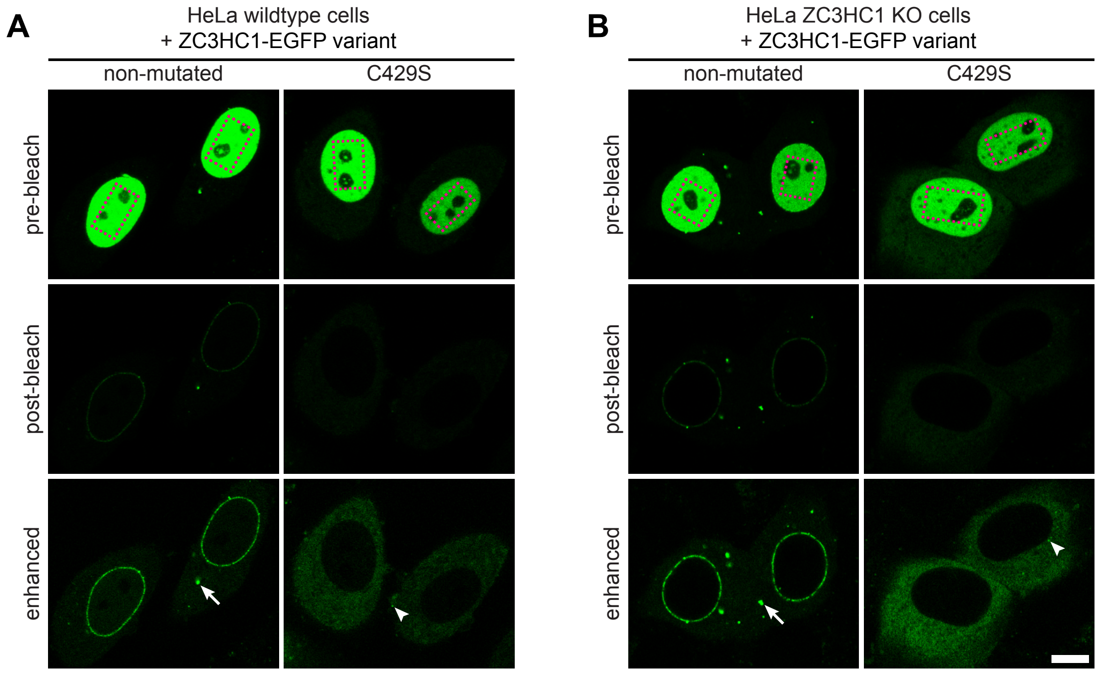

### Supplemental Figure S5

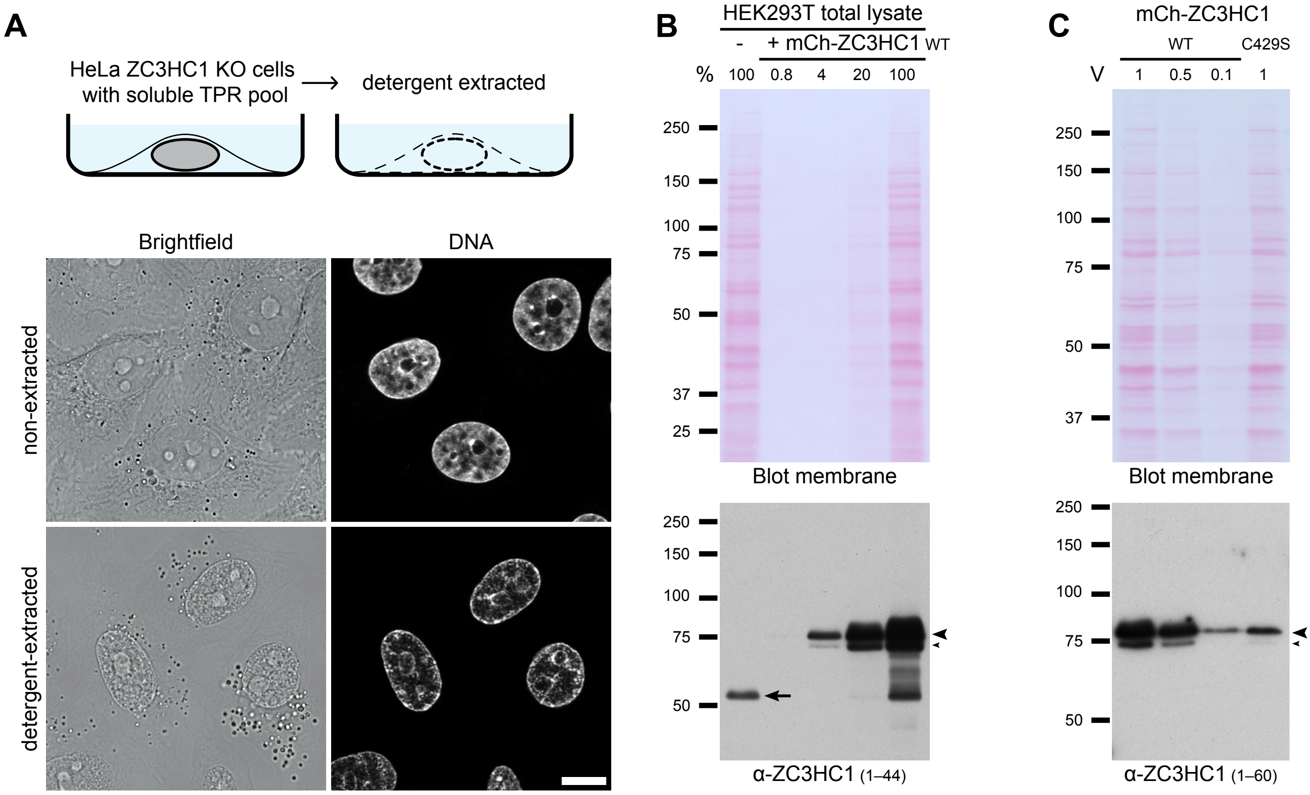

Supplemental  Figure S6

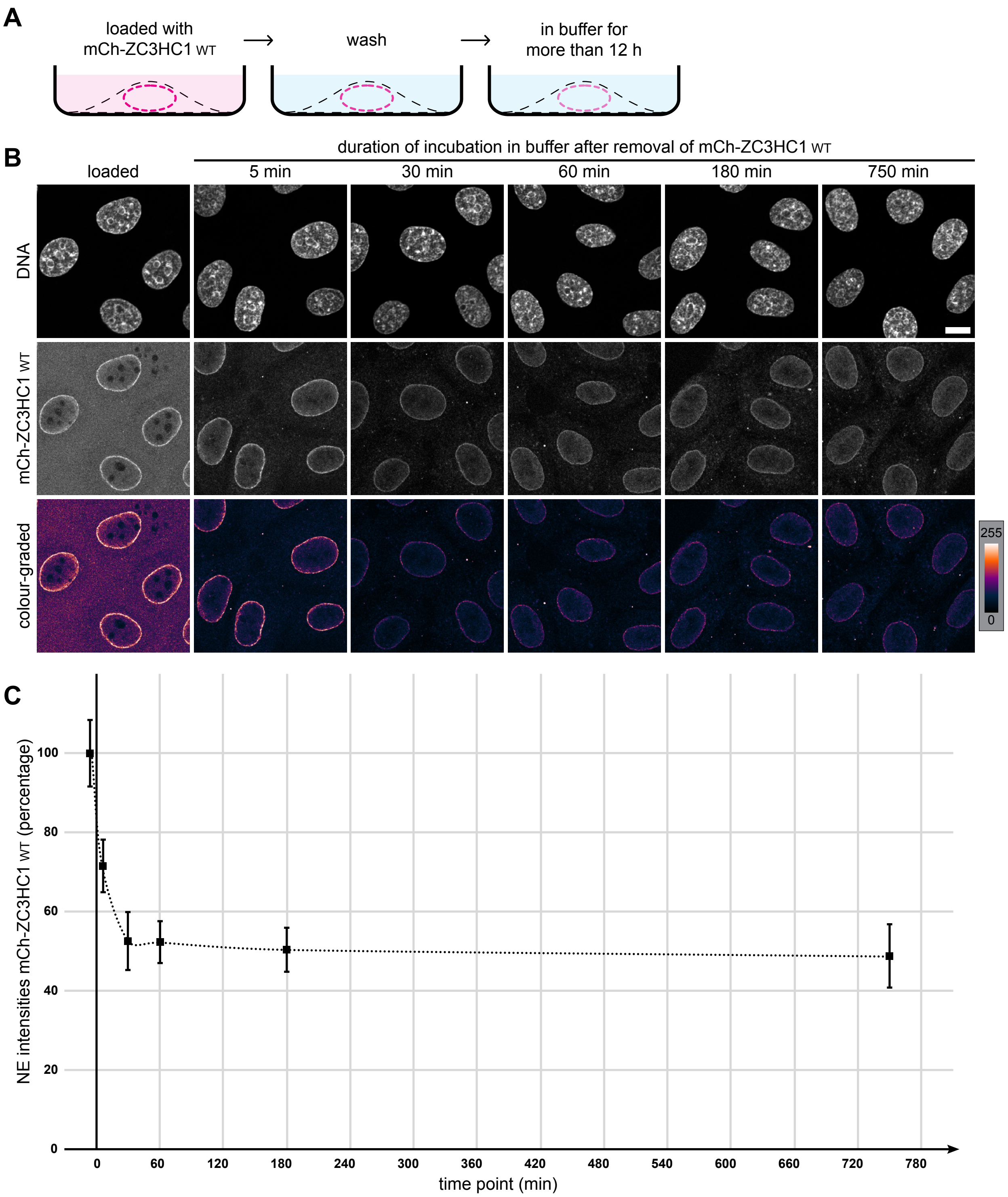

#### Supplemental Figure S7

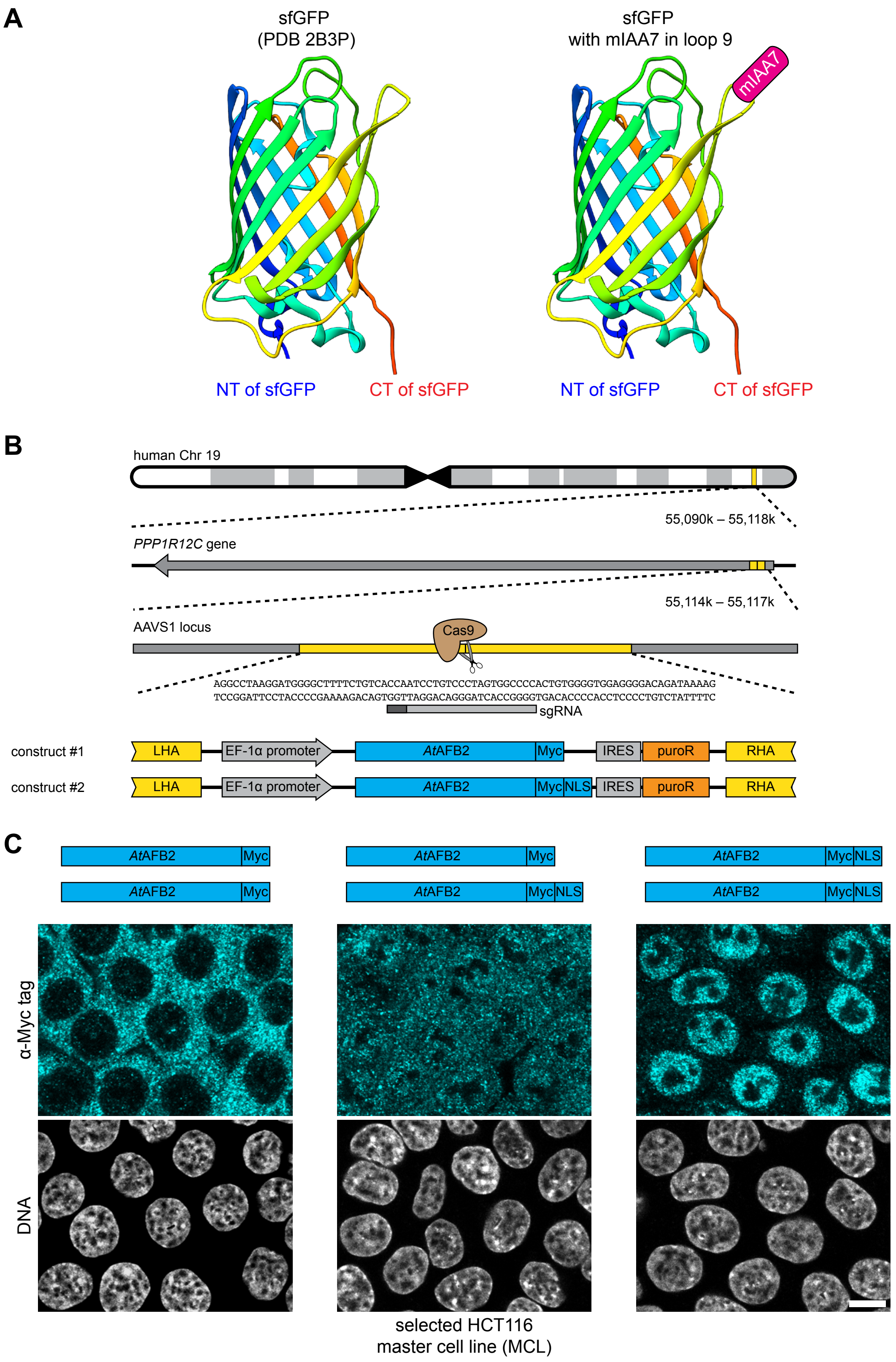

Supplemental Figure S8

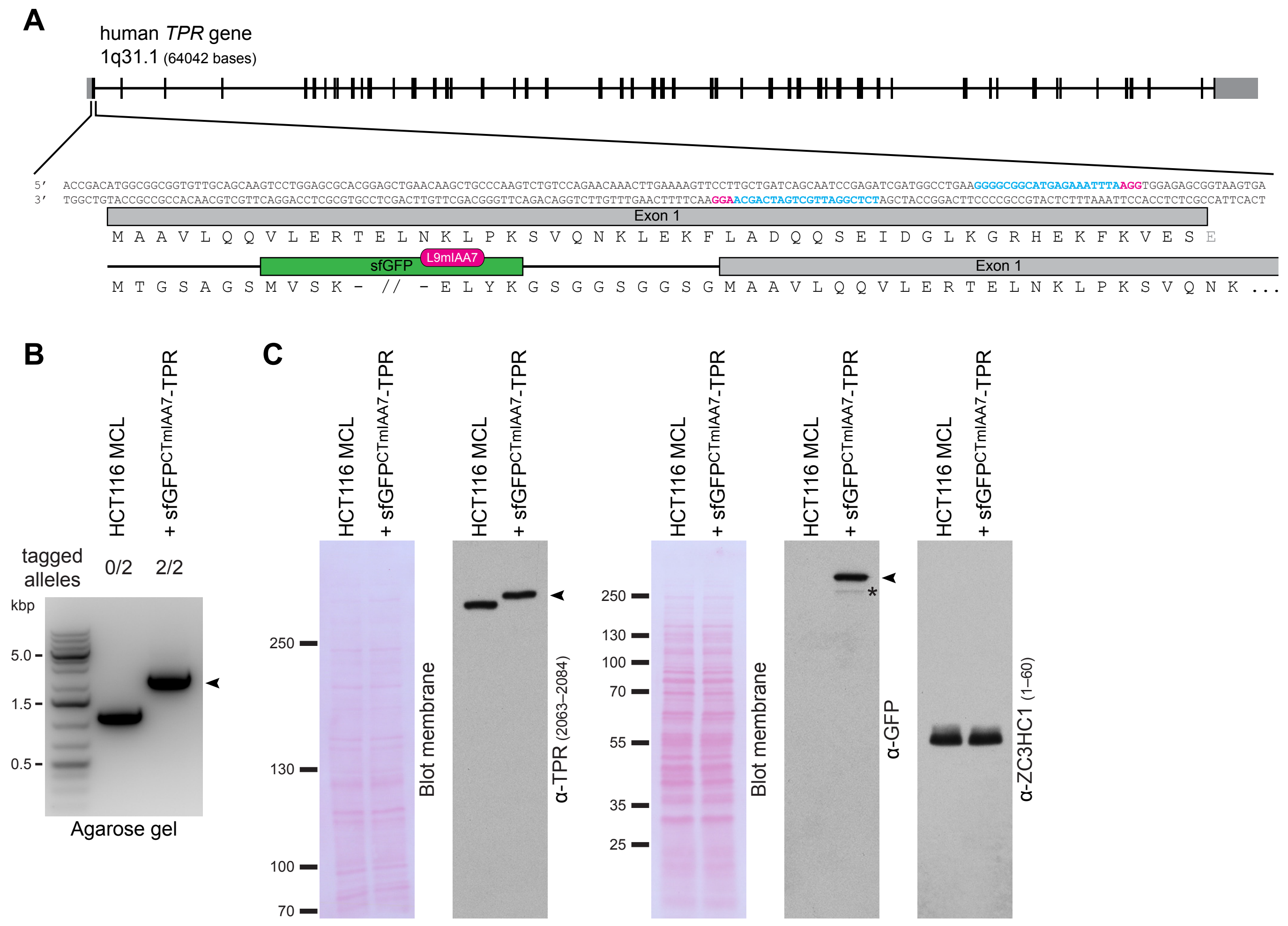

Supplemental Figure S9

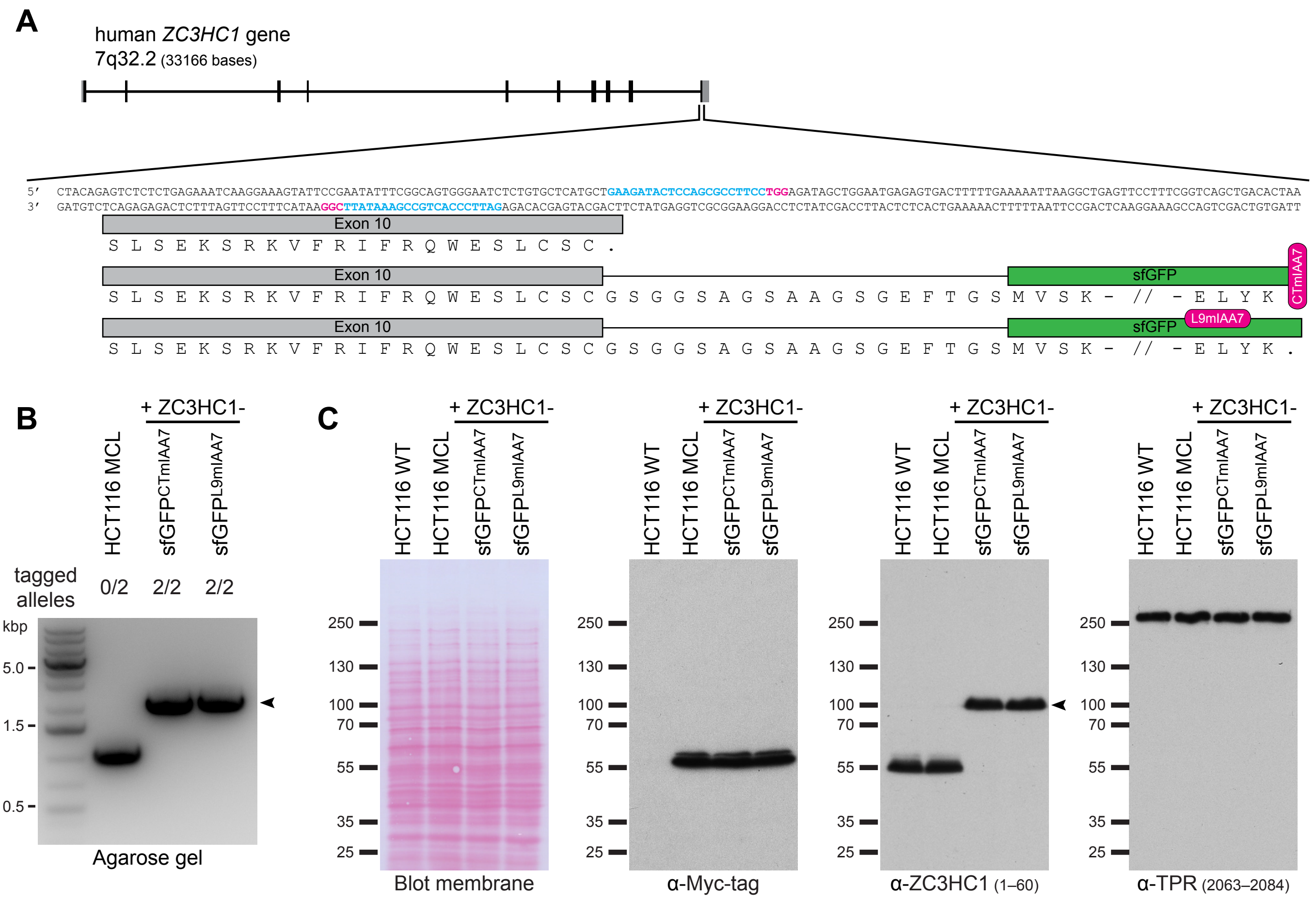

Supplemental Figure S10

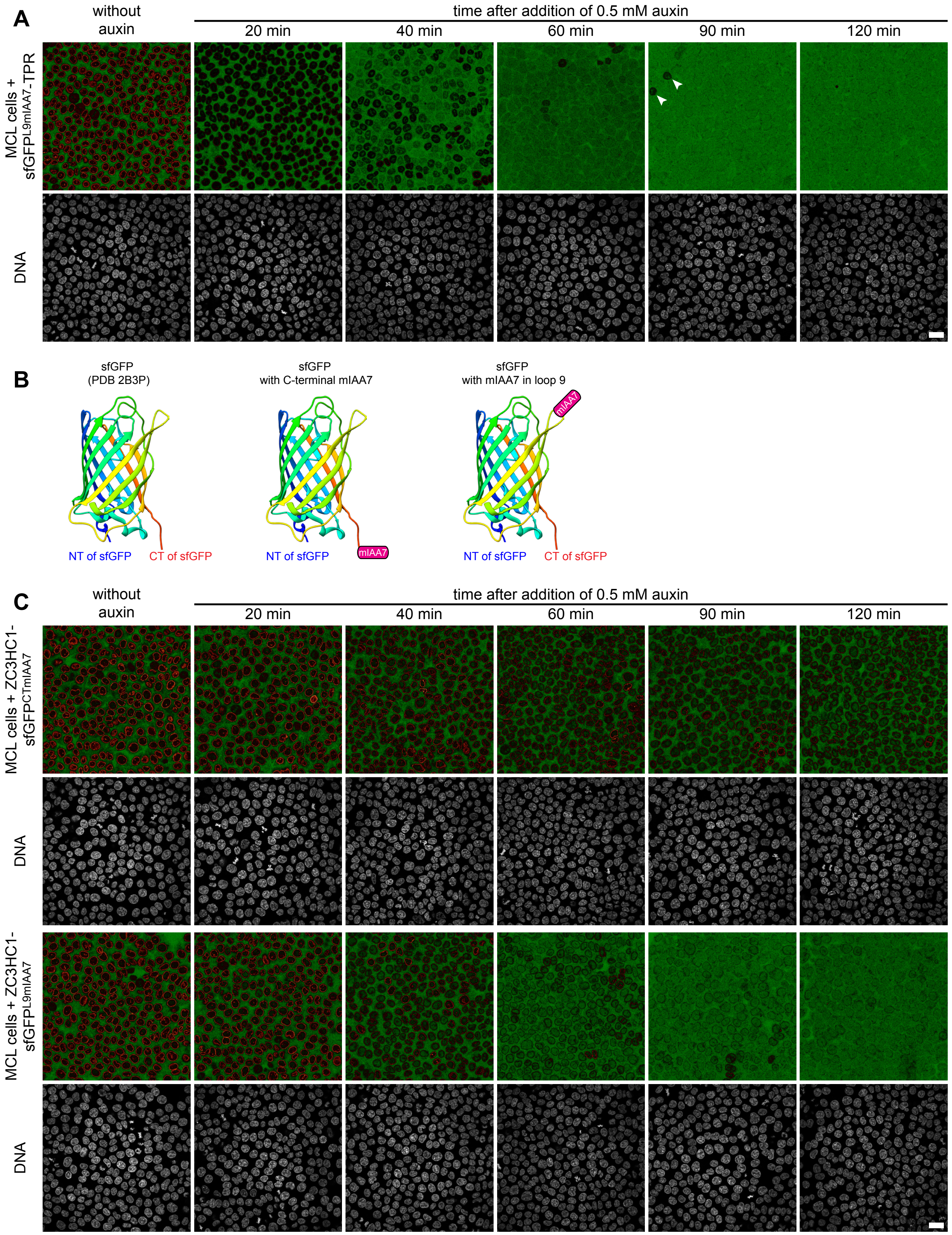

### Supplemental Figure S11

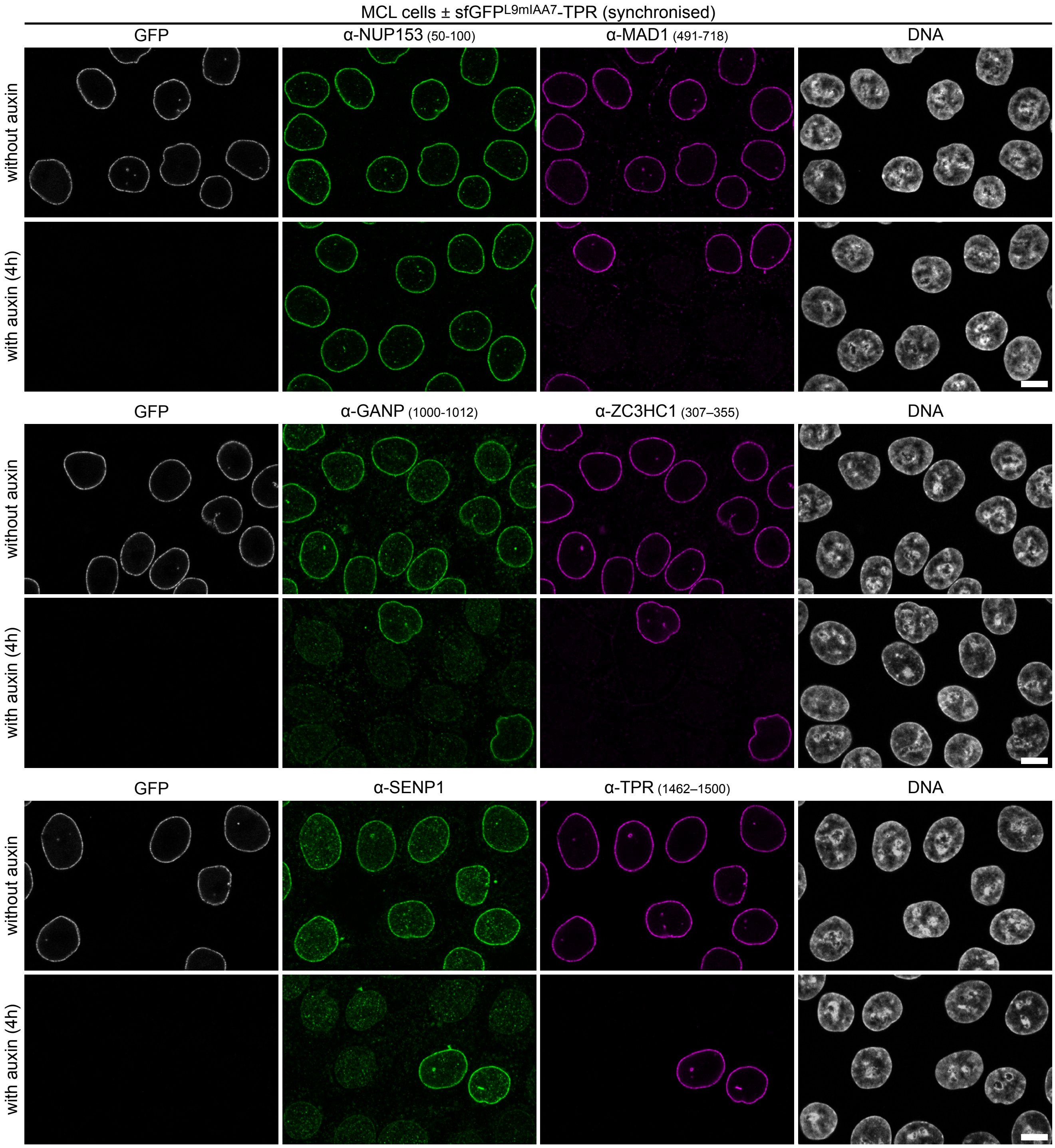

### Supplemental Figure S12

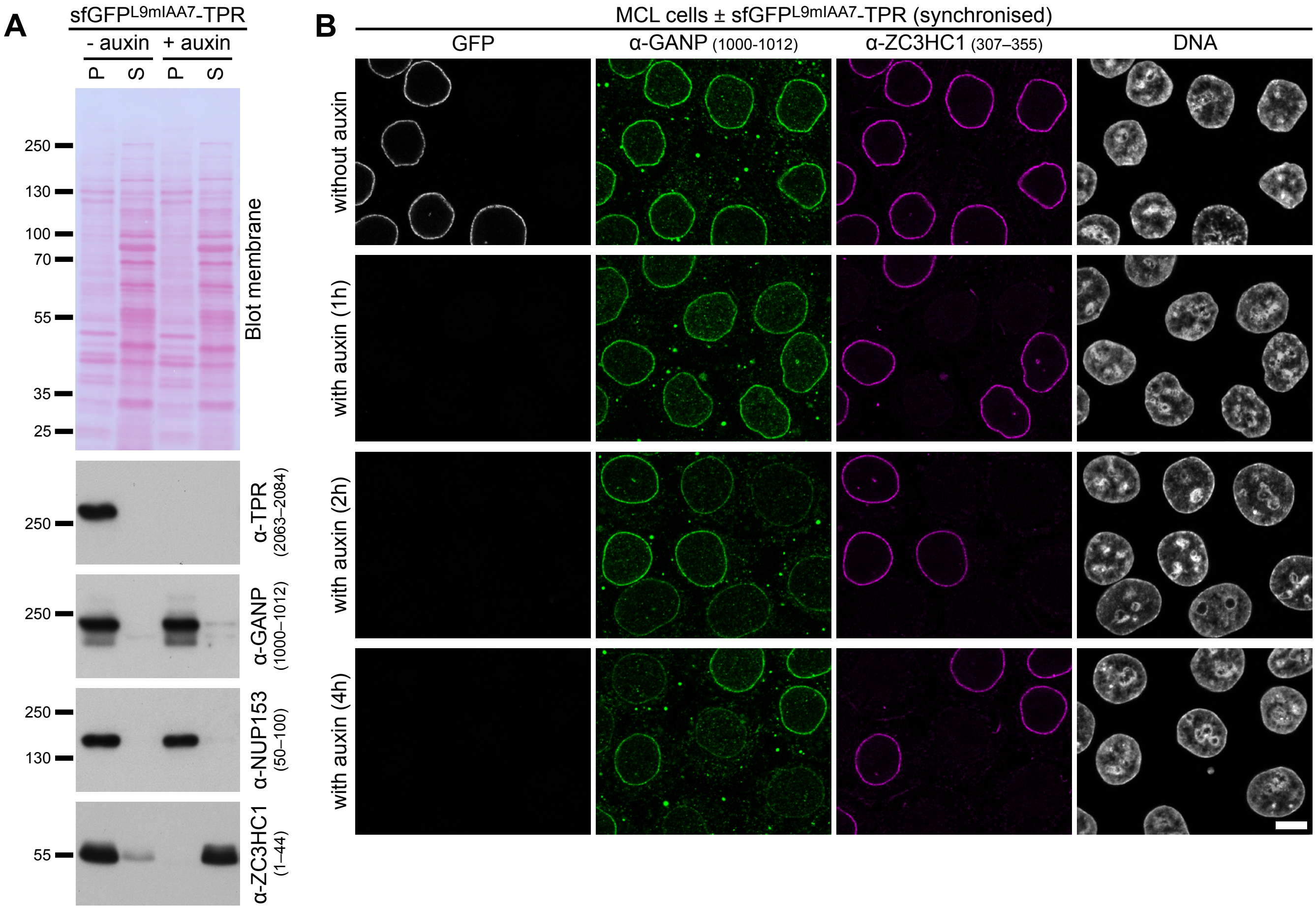
